# VirusImmu: a novel ensemble machine learning approach for viral immunogenicity prediction

**DOI:** 10.1101/2023.11.23.568426

**Authors:** Jing Li, Zhongpeng Zhao, ChengZheng Tai, Ting Sun, Lingyun Tan, Xinyu Li, Wei He, HongJun Li, Jing Zhang

## Abstract

**Background:** The viruses threats provoke concerns regarding their sustained epidemic transmission, making the development of vaccines particularly important. In the prolonged and costly process of vaccine development, the most important initial step is to identify protective immunogens. Machine learning (ML) approaches are productive in analyzing big data such as microbial proteomes, and can remarkably reduce the cost of experimental work in developing novel vaccine candidates.

**Results:** We intensively evaluated the immunogenicity prediction power of eight commonly-used ML methods by random sampling cross validation on a large dataset consisting of known viral immunogens and non-immunogens we manually curated from the public domain. XGBoost, kNN and RF showed the strongest predictive power. We then proposed a novel soft-voting based ensemble approach (VirusImmu), which demonstrated a powerful and stable capability for viral immunogenicity prediction across the test set and external test set irrespective of protein sequence length. VirusImmu was successfully applied to facilitate identifying linear B cell epitopes against African Swine Fever Virus as confirmed by indirect ELISA in vitro.

**Conclusions:** VirusImmu exhibited tremendous potentials in predicting immunogenicity of viral protein segments. It is freely accessible at https://github.com/zhangjbig/VirusImmu.

## Introduction

Vaccines are critical for disease prevention against emerging viral infections [1, 2]. Although many vaccines have been developed and are in use, infectious diseases are still major threats to people’s health, especially during new epidemic outbreaks [3, 4]. The use of recombinant proteins as immunogens is able to meet the demands of future vaccinology and is considered to be the most promising approach for novel vaccines’ development [5].

Immunogenicity is the property of molecules (proteins, lipids or carbohydrates, or combinations thereof) that can elicit specific antibodies against the pathogens [6, 7]. The immunogen triggering the protective immune response is the protective antigen [8]. As protective antigen can help vaccine developers create promising vaccine candidates [9], the identification of protective antigen capable of triggering a significant humoral immune system response is the most important first step in the design and development of new vaccines [3].

In the past decade, the methods for predicting the immunogenicity of protein antigens [3] fall into the two main categories (filtering and classifying) [1]. The filtering-based prediction methods select the most likely vaccine candidates using a serious of filters such as subcellular localization, adhesion probability, topology and sequence similarity with human proteins with the typical examples of Vaxign [10], ANTIGENpro [11], Vacceed [12], Jenner-predict [13], VacSol [14], and Protectome analysis [15]. The classifying prediction approaches applied machine learning (ML) methods to classify a protein as immunogen/non-immunogen [16]. The predicted candidates ranked in top positions are useful for preclinical confirmatory assays [17].

The most representative method of classifying prediction is VaxiJen[18], which proposed an alignment-independent method for antigen prediction based on auto cross covariance (ACC) transformation of protein sequences into uniform equal-length vectors. Wold’s z-scales were used to describe the main physicochemical properties of the amino acids building the tested proteins, which were then classified as protective antigens or non-antigens by partial least squares (PLS)-based discriminant analysis. VaxiJen is one of the few tools currently allowing classification based solely on the physiochemical properties of protein sequences [17]. VaxiJen was then improved by utilizing support vector machines (SVM) to derive the prediction model [1, 19]. Besides, six supervised ML methods were also applied to enhance the performance of the model [3] with focuses on bacterial immunogenicity prediction leaving immunogens’ prediction for viruses hardly untouched.

To overcome these limitations, we proposed an ensemble machine learning approach (VirusImmu) for viral immunogenicity prediction. Unlike VaxiJen only using a single traditional regression algorithm [18] or simply based on majority voting [20], our VirusImmu adopted a soft voting approach to construct the best ensemble model based on careful evaluation of predictive power of eight commonly-used machine learning methods. VirusImmu achieved better predictive power on the test set and external test set. Additionally, we generated VirusImmu-based software for Windows and Linux operating systems, respectively. We also presented a real application of VirusImmu in identification of linear B cell epitopes against African Swine Fever Virus (ASFV). These results provide extensive and powerful evidence about the immunogenicity prediction power of VirusImmu.

## Results

### Performance evaluation of eight machine learning models

The feature selection, performance evaluation and new model development are illustrated in Figure 1A. We curated a dataset including a total of 100 antigens and 100 non-antigens from public domains, which were characterized by Z- and E-descriptors under different lags (L), followed by transforming them into ACC terms (Figure 1B-C). For feature selection, we applied a 10-fold cross validation strategy upon randomly extracted 80% of the transformed dataset, and the process was repeated 100 times. The importance distribution of each ACC term was determined by counting its frequency of being selected by Random Forest (RF) as a feature in the 100 randomized datasets (Figure 1D). To determine the optimal features, we used a varying number of features ranked on the top according to the sorted importance distribution as input to the eight commonly-used machine learning models (PLSR, SVM, LR, XGBoost, Ada, kNN, MLP and RF), respectively. The immunogenicity prediction power for each group of features was evaluated by averaging the AUC values of all the eight models across the 100 randomized datasets. The 22 ACC terms from E-descriptors under L=8 achieving the best performance were determined as the optimal features (Figure 1E). All the analyses below were performed based on the 22 features.

**Figure 1.**
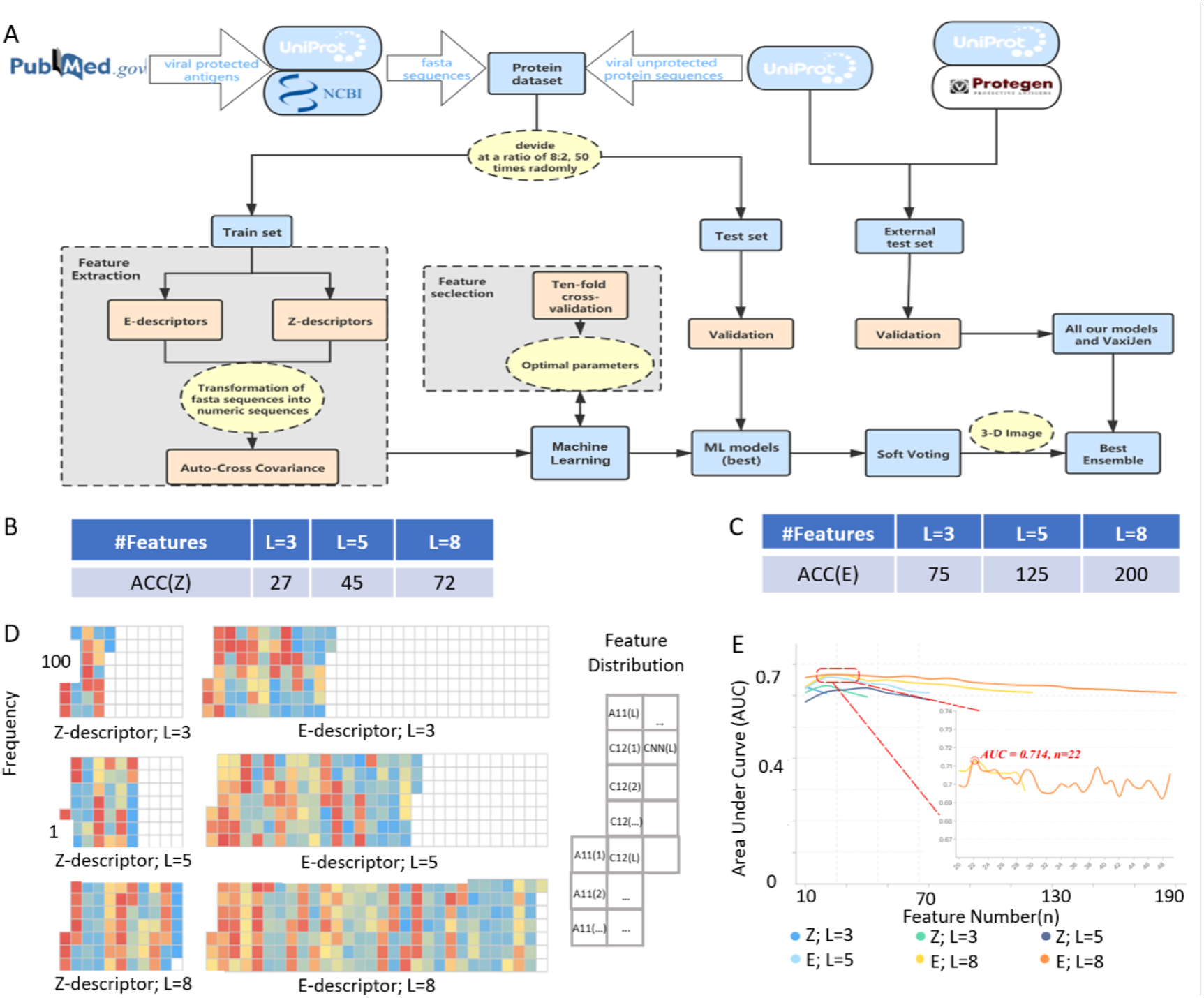
The process of collecting datasets, building models and the feature selection (FS) of present study. (A) Workflow of machine learning (ML) model’s development in the present study. The viral protein dataset was randomly divided into training set and test set at a ratio of 8:2. The training set was used for developing models, the test set and external test sets for validation. (B-C) ACC transformed results with different descriptors (Z,E) and *L*(3,5,8) for each protein. Z and E are Z-descriptor and E-descriptor, respectively. (D)Heat map of the feature importance distribution in different descriptors and lags (*L*) in training sets. (E) The average AUC value across 100 test sets varies with the number of top ACC terms in different descriptors and lags (*L*).

We then conducted intensive experiments to evaluate the performance of the eight machine learning models in predicting immunogenicity of antigens through a random sampling-cross validation strategy. A total of 50 rounds’ randomizations were performed with each round having the dataset divided into training set and test set at a ratio of 8:2. The training set was applied to train each model, and the trained model was then evaluated for immunogenicity prediction on the test set. The average ROC statistics across 50 rounds’ randomizations (Table 1) demonstrated that RF had the most powerful predictive power (AUC=0.770; Accuracy=0.665; Precision=0.628; Recall=0.830; F1=0.712), followed by XGBoost (AUC=0.745; Accuracy=0.669; Precision=0.651; Recall=0.748; F1=0.692) and kNN (AUC=0.708; Accuracy=0.650; Precision=0.644; Recall=0.680; F1=0.658) with all the other five models having AUC less than 0.7 (varying from 0.629 to 0.684). The average ROC curves of kNN, RF, and XGBoost (confidence interval of 95%) on the test set were shown in Figure 2A-C (Table 1).

**Figure 2.**
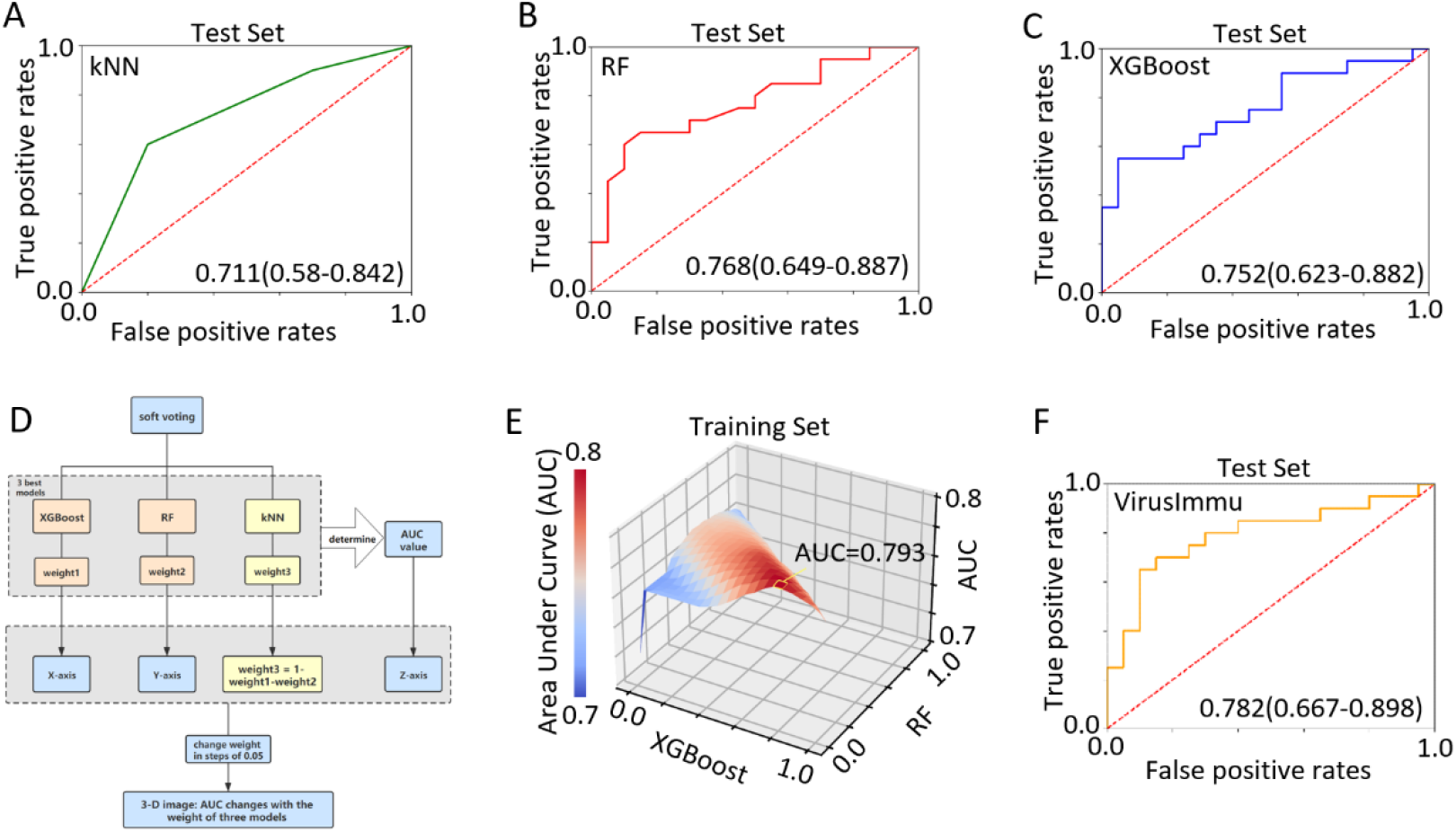
Three-dimensional image of AUC values varying on different soft voting ensemble classifiers. (A) The average ROC curve of kNN (AUC=0.711), (B) RF (AUC=0.768) and (C) XGBoost (AUC=0.752) on test sets. (D) The principle and progress of the 3-D image. (E) The optimal soft voting ensemble classifier appears on the weight of 0.05: 0.75: 0.20 (XGBoost: RF: kNN), with the highest AUC of 0.793 in the training set. The X axis represents the weight for XGBoost model, the Y axis represents the weight for RF model, and the Z axis is the AUC value on different soft voting models. The weight of kNN model can be directly derived when the other two models’ weights are fixed, as the sum of the weights for the three models is 1. (F) The average ROC curve of our ensemble model (VirusImmu) (AUC=0.782) on test sets.

**Table 1:**
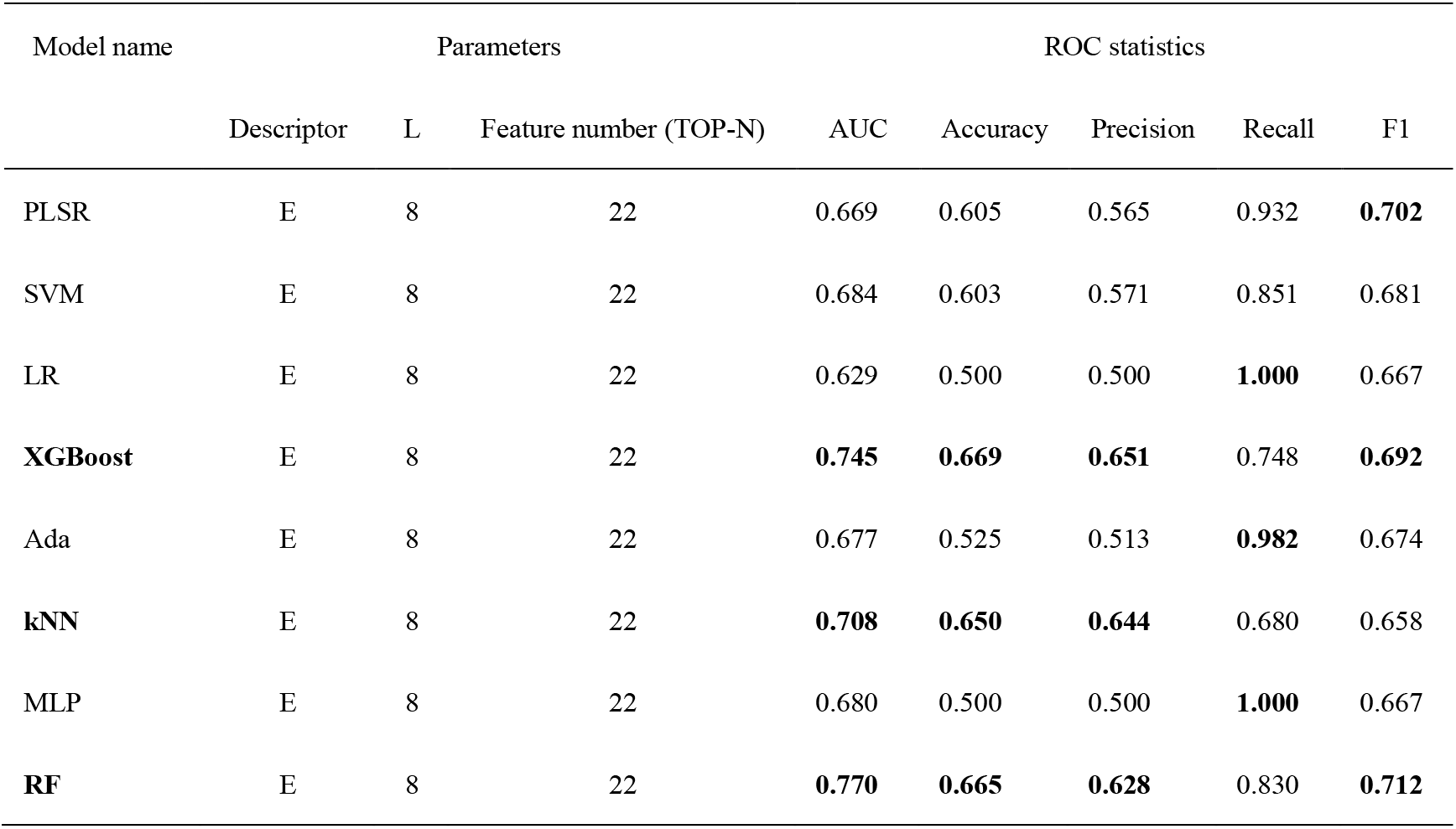
Average ROC statistics of eight commonly-used machine learning models.

| Model name | Parameters |  |  | ROC statistics |  |  |  |  |
| --- | --- | --- | --- | --- | --- | --- | --- | --- |
|  | Descriptor | L | Feature number (TOP-N) | AUC | Accuracy | Precision | Recall | F1 |
| PLSR | E | 8 | 22 | 0.669 | 0.605 | 0.565 | 0.932 | **0.702** |
| SVM | E | 8 | 22 | 0.684 | 0.603 | 0.571 | 0.851 | 0.681 |
| LR | E | 8 | 22 | 0.629 | 0.500 | 0.500 | **1.000** | 0.667 |
| **XGBoost** | E | 8 | 22 | **0.745** | **0.669** | **0.651** | 0.748 | **0.692** |
| Ada | E | 8 | 22 | 0.677 | 0.525 | 0.513 | **0.982** | 0.674 |
| **kNN** | E | 8 | 22 | **0.708** | **0.650** | **0.644** | 0.680 | 0.658 |
| MLP | E | 8 | 22 | 0.680 | 0.500 | 0.500 | **1.000** | 0.667 |
| **RF** | E | 8 | 22 | **0.770** | **0.665** | **0.628** | 0.830 | **0.712** |

### VirusImmu outperforms other commonly-used models

To improve the predictive power of models for immunogenicity, we constructed a soft-voting ensemble classifier (VirusImmu) based on the top three models (RF, XGBoost and kNN). The predictions from RF, XGBoost and kNN were weighted and merged to get the sum of weighted probabilities. To determine the weights for RF, XGBoost and kNN, we enumerated all possible (a total of 232) weights with each weight increasing from 0 to 1 at the incremental step of 0.05 (Figure 2D) and evaluated the performance of the model under different weights using ROC analysis. The optimal soft-voting ensemble model (VirusImmu) obtained in the training set (weight: 0.05 for XGBoost, 0.75 for RF, and 0.20 for kNN, AUC=0.793) (Figure 2E, S1) achieved better performance (average AUC:0.782) than each individual model (average AUC: 0.745 for XGBoost, 0.770 for RF, 0.708 for kNN) (Figure 2F) on test sets.

VaxiJen is one of the few methods using the physiochemical properties of protein sequences to predict immunogenicity. Unlike our VirusImmu, Vaxijen adopted a single traditional regression algorithm or simply majority voting. Therefore, we compared the performance of our VirusImmu with VaxiJen. The average ROC curves demonstrated that VirusImmu outperformed VaxiJen (confidence interval of 95%) in the test set (AUC=0.782 for VirusImmu, AUC=0.75 for VaxiJen) (Figure 3A).

**Figure 3.**
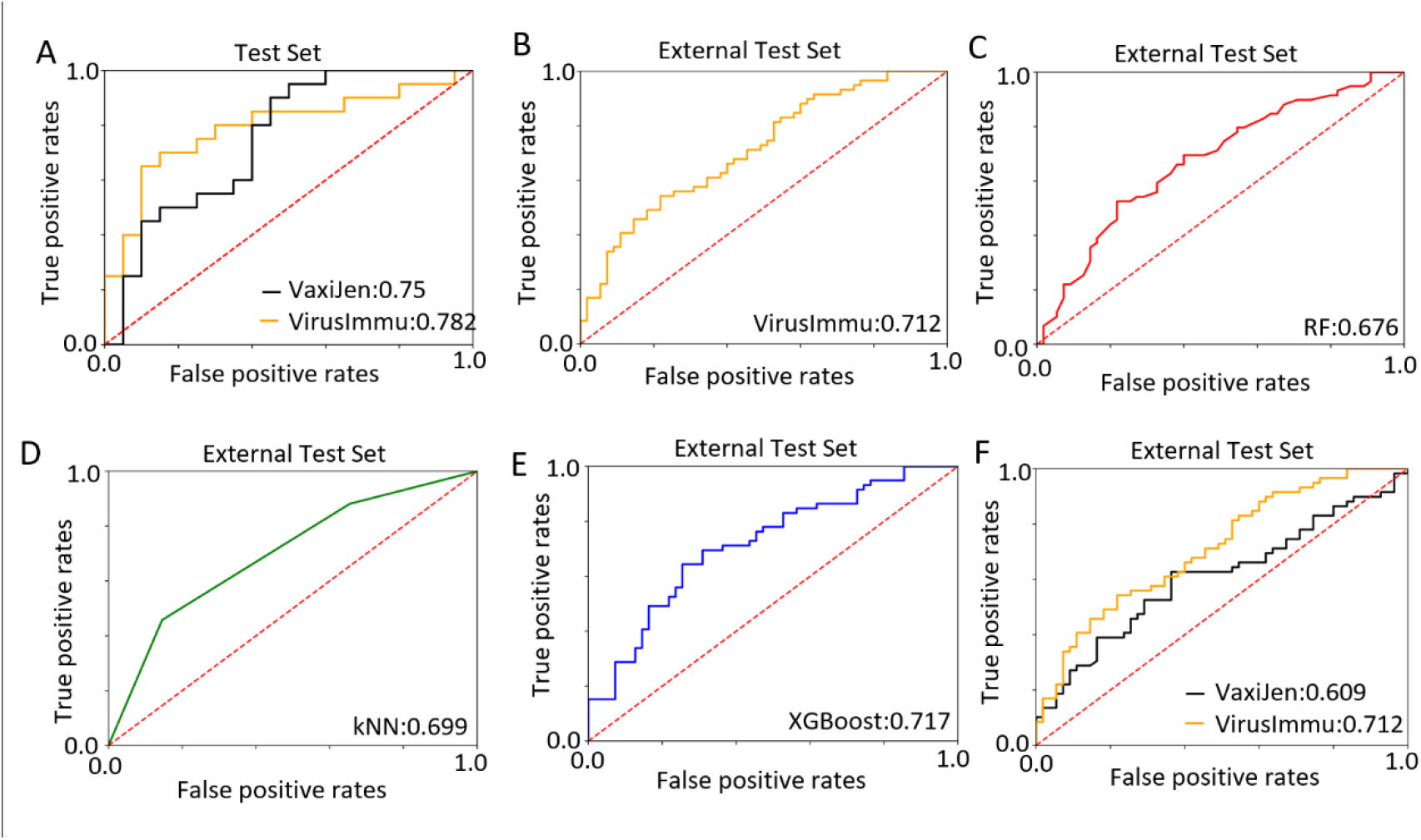
The performance comparisons among VirusImmu and other models. (A)The average ROC curves of our VirusImmu, and VaxiJen in test set, after extracting 20 positive antigens and 20 negative proteins for 50 times randomly. (B-E) The ROC curves of VirusImmu (B), RF (C), kNN (D) and XGBoost (E) in the external test set.(F)The ROC curves of VirusImmu and VaxiJen on the external test set.

To further validate the performance of VirusImmu, we independently collected an external test set containing 59 antigens and 54 non-antigens. The ROC curves revealed that VirusImmu (AUC=0.712) (Figure 3B) outperformed RF (AUC=0.676) (Figure 3C) and kNN (AUC=0.699) (Figure 3D) with similar performance as XGBoost (AUC=0.717) (Figure 3E). VaxiJen had the worst performance on the external test set (AUC=0.609) (Figure 3F). In short, VirusImmu produces more stable predictions for protein immunogenicity than eight commonly-used ML prediction methods and VaxiJen in both the test sets and external test set (Figure 2,3 and S2A). We also compared the performance of VirusImmu with that of two recently released prediction methods, and NetBCE [21] and EpiDope [22]. NetBCE only predicted the immunogenicity for protein sequence smaller than 24 amino acids, however, VirusImmu were able to be applied upon both short and long protein sequence segments. EpiDope combined Embedding from Language Models (ELMo) Deep Neural Networks (DNN) and Long Short Term Memory (LSTM) DNN [22], achieving AUC of 0.667 (Figure S2B) worse than VirusImmu (AUC=0.712).

### VirusImmu achieves better robustness

To test the robustness of all models, we performed 50 rounds’ random sampling with each round using about 30% samples from antigen and non-antigens in the external test set. VirusImmu achieved better performance than VaxiJen in AUC and F1 metrics (Figure 4 A-B). As the predictive power of models may probably be affected by protein sequence length, we divided external test set into five groups according to protein sequence length at the incremental step of 200 bp (Figure 4C), followed by performing 50 rounds’ random samplings. For protein sequence length less than 200 bp, VirusImmu achieved good performance (median AUC=0.665), second to kNN (median AUC=0.725), but much better than VaxiJen (P<0.001) (median AUC=0.546), XGBoost (median AUC=0.615), and RF (median AUC=0.592) (Figure 4D). For 200-400 bp or 400-600 bp proteins, XGBoost achieved the best performance (median AUC = 0.762 and 0.655), followed by VirusImmu (median AUC=0.724 and 0.616), much better than kNN, RF, and VaxiJen (P<0.001) (Figure 4E-F). For 600-800 bp proteins, VirusImmu outperformed all other models with median AUC of 0.868 (P<0.001) (Figure 4G). For proteins with sequence length larger than 800bp, all models achieved similar performances with VaxiJen slightly better than other models (Figure 4H).

**Figure 4.**
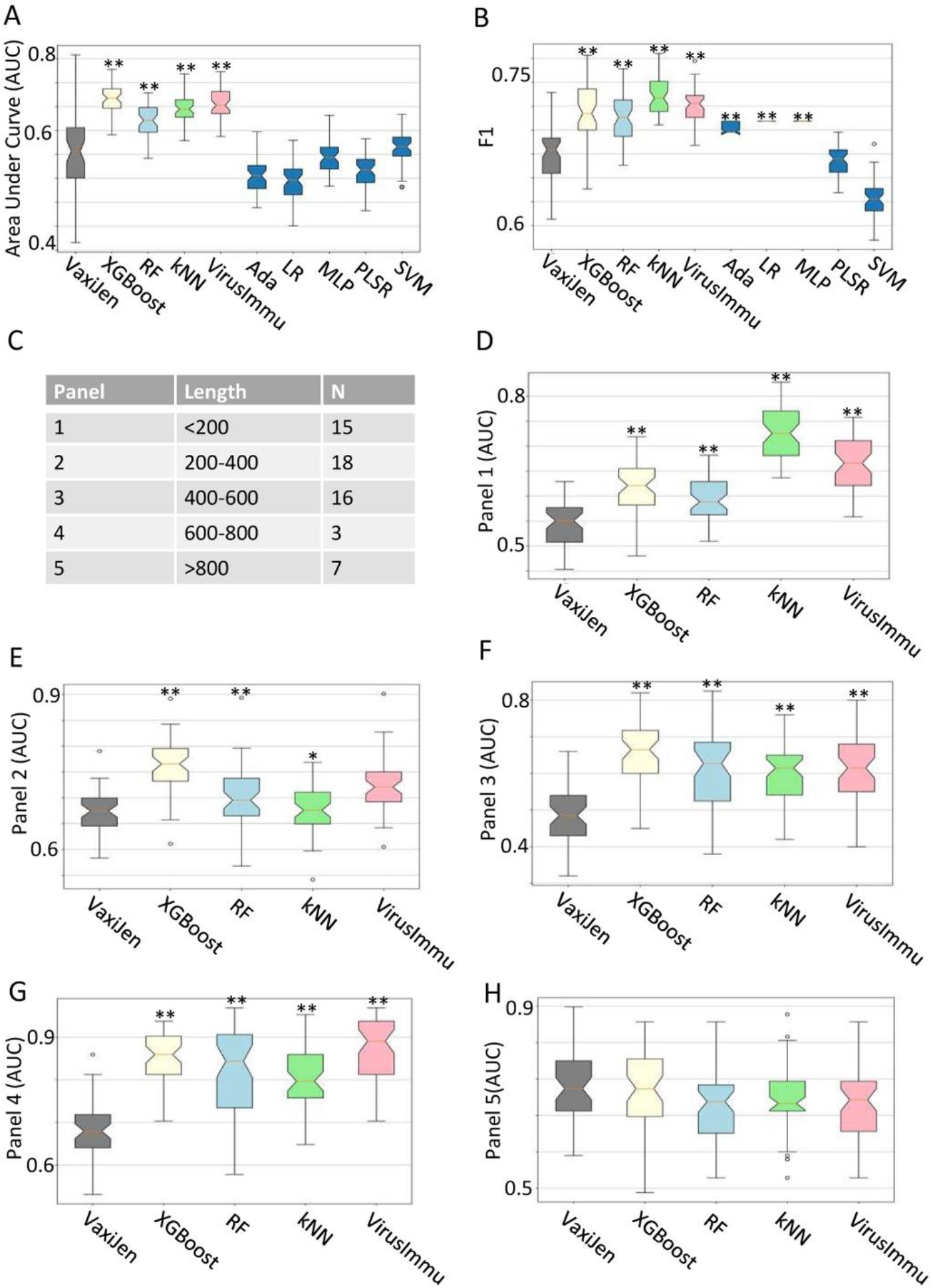
Robustness comparisons of VirusImmu and other models. (A) Box-plots of AUC values in external test set. (B) Box-plots of F1 in external test set. (C) Five groups divided by length of the external validation set, N is the number of proteins of every panel. (D-H) Box-plots of AUC on VirusImmu, XGBoost, RF, kNN and VaxiJen in the external test subset consisting of proteins with sequence length < 200 (D), 200-400 (E), 400-600 (F), 600-800 (G) and > 800 (H). *: P<0.01; **: P<0.001; Wilcoxon test is performed between each of our model and VaxiJen.

Both XGBoost and VirusImmu achieved good performances (the top two) in external validation data, with XGBoost having slightly better AUC (Figure 4A) but worse F1 (Figure 4B) than that of VirusImmu. XGBoost also performed worse for <200 bp and 600-800 bp proteins than VirusImmu. As most epitopes are protein segments with length less than 200, VirusImmu have better application scenario than XGBoost.

To further prove the reliability of VirusImmu, we selected epitopes for SARS-CoV-2 in published literatures[23–26] to verify the immunogenicity prediction ability of VirusImmu. The results showed that 14 of the total 15 epitopes involved in the four literatures were predicted as antigens by VirusImmu (Table S5), verifying VirusImmu’s good performance for viral protein immunogenicity prediction. To facilitate the use of VirusImmu, we also developed VirusImmu-based software for Windows and Linux operating systems, respectively, which are freely available for downloading at ‘http://www.oncoimmunobank.cn/software/item/virusimmu’.

### VirusImmu facilitates identifying peptide vaccine candidates for African Swine Fever Virus (ASFV)

As there is no effective vaccine or treatment for ASFV, it is required to identify protective antigens. ASFV pp220 polyprotein essential for virus structural integrity [27] was found containing epitopes that can induce robust immune responses in pigs, suggesting it will be promising candidate in vaccine development [27].

To identify antigenic epitopes, we predicted 1376 B-cell linear epitope candidates from pp220 protein using 17 most popular methods including BCPred, Immune Epitope Database (IEDB) server (Bepipred2.0, Bepipred, Emini Surface Accessibility, Kolaskar & Tongaonkar Antigenicity, Chou & Fasma Beta-Turn, Karplus & Schulz Flexibity, and Parker Hydrophilicity), AAP, FBCPred, Bcepred (Hydrophilicity, Flexibility, Accessibility, Turns, Exposed Surface, Polarity, and Antigenic Propensity), and ABCpred (Figure 5A, Table S6). Stringent criteria were used to filter out epitopes having antigenicity predicted by VaxiJen ≤ 1.3, resulting in 29 epitopes remained, of which 12 epitopes were classified as non-allergen and non-toxin (Figure 5A). VirusImmu predicted eight of the 12 epitopes to be antigenic (Figure 5A, Table S7).

**Figure 5.**
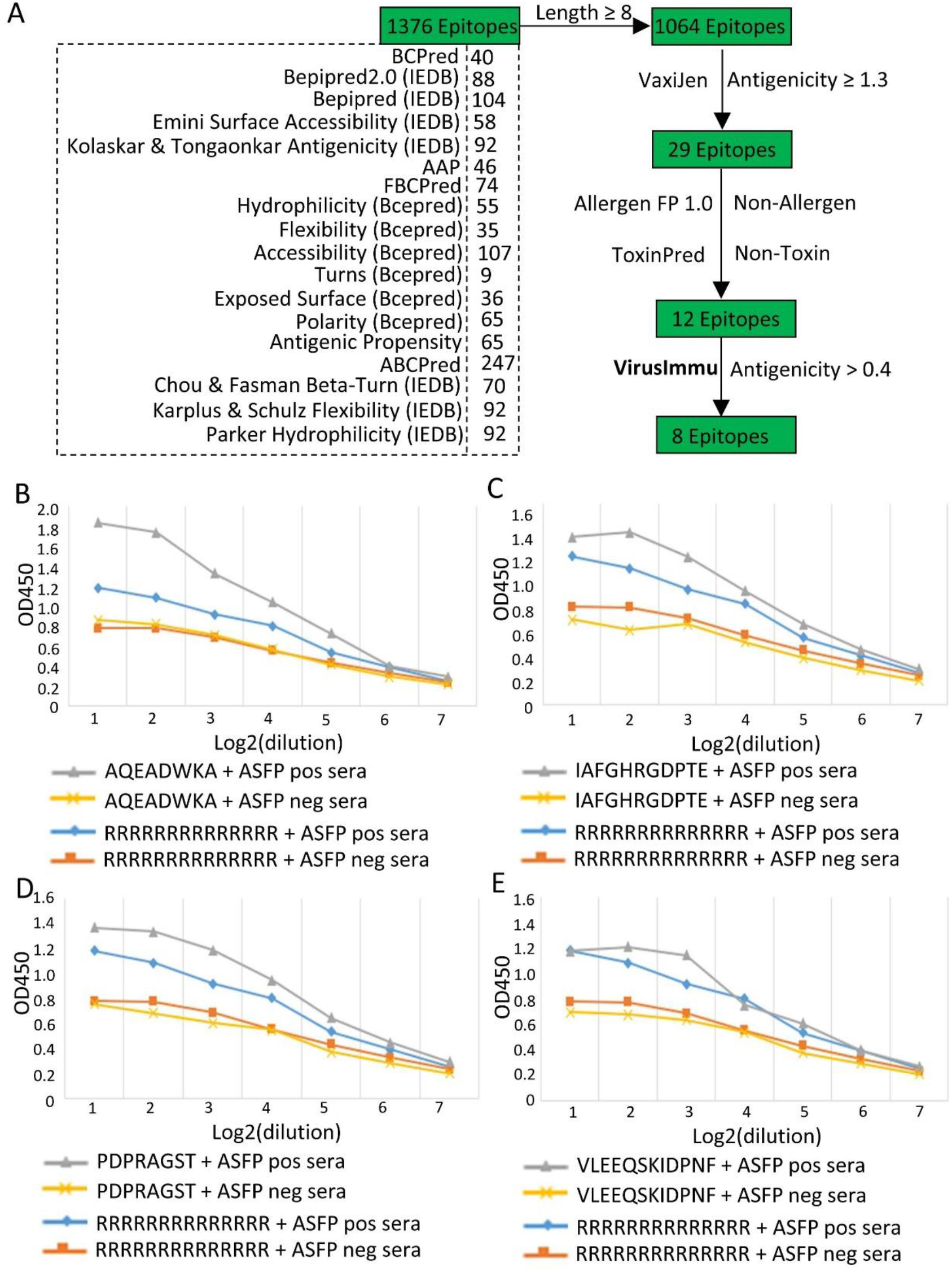
Measurements of antigenic B cell epitopes binding to antibody. (A) Framework of antigenic linear B-cell epitopes identification. (B-D) The binding affinity assessed by ELISA between sera antibodies from pigs infected with ASFV SY18ΔMGF/ΔCD2v and linear B-cell epitopes of AQWADWKA (B), IAFGHRGDPTE (C) and PDPRAGST (D) predicted by VirusImmu as antigenic. (E) The binding affinity assessed by ELISA between sera antibodies from pigs infected with ASFV SY18ΔMGF/ΔCD2v and linear B-cell epitopes (VLEEQSKIDPNF) predicted by VirusImmu as non-antigenic. y-axis is the average OD450 (Optical Density) value of two replicates under each dilution. x-axis is the dilution.

To confirm the binding of the eight epitopes with serum IgG antibodies against ASFV, we collected pooled sera from five ASFV infected pigs and five healthy pigs, respectively. Indirect ELISA assay confirmed that seven antigenic linear B-cell epitopes (Figure 5B-D, S3A-D) but one (Figure S3E) were reacting specifically and dose dependently with sera antibodies from the ASFV infected pigs but not in healthy ones, whereas the arbitrary control peptides (‘RRRRRRRRRRRRRR’) had no effects. The epitope predicted by VirusImmu as non-antigenic (‘VLEEQSKIDPNF’) exhibited no specific binding with sera antibodies either (Figure 5E). These results provide a powerful example of VirusImmu application in real scenarios.

## Discussion

Experimentally validating all epitopes predicted by different tools is simply not feasible. The current tools, such as VaxiJen [18], use traditional classification methods with low accuracy and stability. Machine learning has been proven to provide higher performance [28]. Nevertheless, the selection of the optimal features and coding strategies, and a more comprehensive external test set, are required for stably predicting immunogenicity of diverse viral proteins.

In this study, we proposed an ensemble machine learning approach (VirusImmu) for virus antigenicity prediction, achieving promising results in both internal and external validation sets. VirusImmu significantly outperforms existing prediction tools that used classifiers to predict viral protein’s immunogenicity, with higher accuracy (AUC=0.782) and greater robustness in external test sets. Protein length was considered and validated, and our model showed more stable predictions on viral protein datasets of various length compared to previous model [18]. Users have no need to consider the length of viral proteins, which is critical to develop vaccines for virus.

VirusImmu exhibited powerful immunogenicity prediction and huge potentials in real life applications. We applied VirusImmu to help identify B-cell epitopes against AFSV. Of 12 linear B-cell epitopes were predicted as antigenic by the widely used immune-informatics methods, eight were also predicted as antigenic by VirusImmu. Indirect ELISA assays confirmed that seven of the eight linear B-cell epitopes predicted by VirusImmu as antigens reacted specifically and dose dependently with sera antibodies from the pigs infected with ASFV, suggesting that they may induce antibodies against ASFV which needs to be further confirmed by virus neutralization assay.

Despite these advances, classifying-based prediction tool still [18] has the drawback of low accuracy rate compared with the filtering-based method [29]. The lack of accuracy may be due to the fact that the data is not comprehensive enough and the associated machine learning algorithms are not well matched. VirusImmu adopted Z-descriptor and E-descriptor to describe the physicochemical properties of the residues, but there are also other descriptors such as CKSAAP [30–33] which may improve the performance. The future aim should be implemented with higher accuracy without throwing away proteins or restricting protein length. Therefore, our direction for endeavors is to expand the data and find more machine algorithm models for viral proteins’ immunogenic prediction.

## Conclusions

We successfully developed a machine learning ensemble approach (VirusImmu) in viral antigens’ immunogenicity prediction. VirusImmu, which is not based on sequence comparison and excludes the effect of protein sequence length, is applicable to the prediction of proteins and peptides with higher accuracy and greater universality compared to similar predictive tools. VirusImmu can be used to predict the immunogenicity of viral proteins, to select proteins with higher immunogenicity before epitopes prediction, and to predict specific epitopes’ immunogenicity, providing a more comprehensive tool for researchers in vaccine development. VirusImmu-based software and codes are freely accessible at ‘http://www.oncoimmunobank.cn/software/item/virusimmu’.

## Methods

### Dataset of training and test

Protein datasets were constructed by 100 antigens (Positive set) (Table S1) and 100 non-antigens (Negative set) (Table S2). Protected antigens were the validated protein antigens curated from the literatures, and the corresponding protein sequences were from UniProt and NCBI. Protein with complete fragments were preferred. Unprotected protein sequences (non-antigens) were randomly selected from Viral Bioinformatics Resource Center [18]. A BLAST was performed to confirm that non-antigens have no sequence identity to the antigens. A random sampling cross validation strategy was applied to derive test set from 20% of the positive and negative dataset. 50 randomizations were performed.

### Dataset of external evaluation

The external dataset was independently constructed (Table S3), consisting of 59 antigens and 54 non-antigens with the antigens’ sequences manually curated from UniProt and the Protegen database [34] and the non-antigens’ sequences selected randomly from UniProt in the same way of training set.

### Descriptors

Five E-descriptors [35] (Table S4) were used to characterize each protein, which was represented as a two-dimensional array (5 × *N*) by using the E-descriptors [35], where 5 is the number of descriptors and N is the protein length. Three Z-descriptors (3 × *N*) [36] were also used to represent protein and compared the difference between the two descriptors in the process of viral immunogenicity prediction.

### Auto-Cross Covariance (ACC)

An auto-cross covariance (ACC) transformation [37] was used to transform proteins to a uniform equal-length vector. Auto covariance *A*_*jj*_(*lag*) and Cross covariance *C*_*jk*_ (*lag*) were calculated by the following formulas:

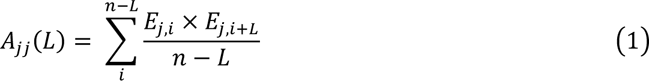

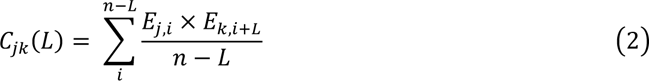

Index *j* and *k* refer to the the Z-scales (1, 2, 3) or E-scales (1, 2, 3, 4, 5), *n* is the number of amino acids in a virus protein sequence, index *i* indicates the position of amino acid (*i* = 1, 2, … n) and *L* is the lag (*L*= 1, 2, …*L*). A short range of lag (*L*), which are the lengths of the frame of contiguous amino acids, were used to calculated *A*_*jj*_ and *C*_*jk*_, to investigate the influence of close amino acid proximity on protein antigenicity.

### Feature selection

To identify and rank the most relevant features, each protein was transformed into a uniform vector, which consisted of ACC terms of different lengths. RF is used to calculate the Gini impurity of each ACC term, followed by sorting them in descending order and selecting the top 20 to record. We recursed the above process 100 times, calculated the frequency of each ACC term, and obtained the final feature importance distribution.

### Soft voting ensemble approach

Eight machine learning (ML) models were applied in the present study: Partial Least Squares (PLS) Regression (number of principal components = 2, maximum iterations = 100, iteration stop condition = 1e-06), K Nearest Neighbours (kNN) (number of neighbors = 2, weight mode = uniform, number of leaves = 30), Adaptive Boosting (Ada) (maximum iterations = 100, learning rate = 1.0.), Logistic Regression (LR) (maximum iterations = 1000, penalty = L2), Support Vector Machine (SVM) (error costs = 1, kernel function = RBF), Random Forest (RF) (maximum iterations = 100, feature selection method = gini, maximum depth = 6), Extreme Gradient Boosting (XGBoost) (base learning = tree, maximum depth = 6) and A Deep Learning Model (Multi-Layer Perceptron, MLP) (two hidden layers = 32 & 16, input layer size = input data size, output layer size = 2, activation function = relu, optimization function = adam, maximum iterations = 1000). A soft voting ensemble classifier was designed based on the best three models, in which predictions were weighted on the basis of classifier’s importance and merged to get the sum of weighted probabilities. Detailed information about these algorithms is also available in Supplementary Materials.

### Evaluation criteria

The ML models were trained by training set, validated by test set and then another layer of validation by the external test set. The predictive ability of the models was measured by the area under the Receiver Operating Characteristic (ROC) curve and statistics including accuracy, precision, sensitivity (Recall) and F1, which were calculated by the python package scikit-learn.

### Antigenic linear B-cell epitope prediction for pp220 protein of ASFV

A total of 17 widely used methods were used to predict linear B-cell epitopes for ASFV pp220 protein, including BCPreds [38], 7 methods in Immune Epitope Database (IEDB) server [39], AAP [40], FBCPred [41], 7 methods in Bcepred [42], and ABCpred [43]. VaxiJen [8] was employed to evaluate their antigenicity with stringent criteria to filter out epitopes having antigenicity score ≤ 1.3. Allergen FP 1.0 [44] and ToxinPred [45] were used to predict the allergenicity and toxicity. VirusImmu was applied to predict the antigenicity of the filtered epitopes.

### Animal experiments

Informed consent was obtained from the owners of pigs. All methods were carried out in accordance with relevant guidelines and regulations. All methods were carried out in accordance with the ARRIVE guidelines. Blood samples were collected and pooled for five six-month old healthy pigs that had no detectable antibodies against ASFV in Military Veterinary Institution, Changchun. Pooled sera of five pigs infected with ASFV SY18ΔMGF/ΔCD2v (10^4^ TCID_50_/mL, TCID_50_ represents Tissue Culture Infectious Dose) were provided by Jilin Heyuan Bioengineering Co., Ltd. The eight linear B-cell epitopes and two control peptides were synthesized by Scilight-Peptide Inc., Beijing, China. Indirect ELISA assay was performed to assess the binding of the eight epitopes with serum IgG antibodies against ASFV at the biosafety level 2 facilities in Beijing Institute of Microbiology and Epidemiology, China.

### VirusImmu-based software development

VirusImmu-based software was developed for Windows and Linux operating systems using PySide6 (6.5.0), respectively. Setup file for windows was created by Inno Setup6. Environment for Linux was created by Conda, with the yaml file provided. The software was tested on Windows 10 and Linux (Ubuntu 20.04.6 LTS).

## Abbreviations

ML: Machine learning
ACC: Auto cross covariance
PLS: Partial least squares regression
SVM: Support vector machines
kNN: K Nearest Neighbours
Ada: Adaptive Boosting
LR: Logistic Regression
RF: Random Forest
XGBoost: Extreme Gradient Boosting
MLP: Multi-Layer Perceptron
IEDB: Immune Epitope Database server
ROC: Receiver Operating Characteristic
ASFV: African Swine Fever Virus

## Declarations

### Ethical statements

Informed consent was obtained from the owners of pigs. All methods were carried out in accordance with relevant guidelines and regulations. All methods are reported in accordance with the ARRIVE guidelines (https://arriveguidelines.org). All animal experiments were approved by Ethics Committee of Beijing Institute of Microbiology and Epidemiology, China (the approval number is IACUC-IME-2022-016).

## Consent for publication

Not applicable.

## Availability of data and materials

All supporting data are included within the main article and its supplementary files. The custom codes can be accessed at github https://github.com/zhangjbig/VirusImmu.

## Authors’ contributions

JZ directed the project, interpreted results and wrote the manuscript. JL performed the data analyses and result presentation, and drafted the manuscript. ZPZ, LYT, XYL, and WH performed the biological experiments. TCZ optimized the codes, constructed the software and developed the Download webpage. HJL participated in directing the development of software and revising the manuscript. TS participated in the algorithm development. All named authors read and approved the final manuscript.

## Acknowledgements

We thank Dr. Ying Wang, Dr. Yun Yang and Dr. Wei Chen at Beihang University, Taicang Qingli Biotechnology Co., Ltd and Ling Wei Pharmaceutical Inc for providing valuable comments and discussion about bioinformatics analysis and experiments related to this work.

## References

1. Bowman BN, Mcadam PR, Vivona S, Zhang JX, Luong T, Belew RK, Sahota H, Guiney D, Valafar F, Fierer J: Improving reverse vaccinology with a machine learning approach. Vaccine 2011, 29(45):8156–8164.

2. Afrough B, Dowall S, Hewson R: Emerging viruses and current strategies for vaccine intervention. Clin Exp Immunol 2019, 196(2):157–166.

3. Mao HH, Chao S: Advances in Vaccines. Adv Biochem Eng Biotechnol 2020, 171:155–188.

4. Wang YB, Wang LP, Li P: Perspectives on novel vaccine development. Pol J Vet Sci 2018, 21(3):643–649.

5. Meinke A, Henics T, Nagy E: Bacterial genomes pave the way to novel vaccines. Current Opinion in Microbiology 2004, 7(3):314–320.

6. Rappuoli R, Bottomley MJ, D’Oro U, Finco O, De Gregorio E: Reverse vaccinology 2.0: Human immunology instructs vaccine antigen design. J Exp Med 2016, 213(4):469–481.

7. Gonzalez-Dias P, Lee EK, Sorgi S, de Lima DS, Urbanski AH, Silveira EL, Nakaya HI: Methods for predicting vaccine immunogenicity and reactogenicity. Hum Vaccin Immunother 2020, 16(2):269–276.

8. Immunogenicity Prediction by VaxiJen: A Ten Year Overview. Journal of Proteomics & Bioinformatics 2017, 10(11).

9. Liljeroos L, Malito E, Ferlenghi I, Bottomley MJ: Structural and Computational Biology in the Design of Immunogenic Vaccine Antigens. J Immunol Res 2015, 2015:156241.

10. Xiang Z, He Y: Vaxign: a web-based vaccine target design program for reverse vaccinology. Procedia in Vaccinology 2009, 1(1):23–292.

11. Magnan CN, Zeller M, Kayala MA, Vigil A, Randall A, Felgner PL, Baldi P: High-throughput prediction of protein antigenicity using protein microarray data. Bioinformatics 2010, 26(23):2936–2943.

12. Goodswen SJ, Kennedy PJ, Ellis JT: Vacceed: a high-throughput in silico vaccine candidate discovery pipeline for eukaryotic pathogens based on reverse vaccinology. Bioinformatics 2014(16):2381–2383.

13. Jaiswal V, Chanumolu SK, Gupta A, Chauhan RS, Rout C: Jenner-predict server: prediction of protein vaccine candidates (PVCs) in bacteria based on host-pathogen interactions. Bmc Bioinformatics 2013, 14(1):211.

14. Rizwan M, Naz A, Ahmad J, Naz K, Obaid A, Parveen T, Ahsan M, Ali A: VacSol: a high throughput in silico pipeline to predict potential therapeutic targets in prokaryotic pathogens using subtractive reverse vaccinology. BMC Bioinformatics 2017, 18(1):106.

15. Altindis E, Cozzi R, Palo BD, Necchi F, Mishra RP, Fontana MR, Soriani M, Bagnoli F, Maione D, Grandi G: Protectome Analysis: A New Selective Bioinformatics Tool for Bacterial Vaccine Candidate Discovery. Molecular & Cellular Proteomics 2015, 14(2):418–429.

16. Gonzalez-Dias P, Lee EK, Sorgi S, de Lima DS, Urbanski AH, Silveira EL, Nakaya HI: Methods for predicting vaccine immunogenicity and reactogenicity. Hum Vaccin Immunother 2020, 16(2):269–276.

17. Dalsass M, Brozzi A, Medini D, Rappuoli R: Comparison of Open-Source Reverse Vaccinology Programs for Bacterial Vaccine Antigen Discovery. Frontiers in Immunology 2019, 10:113-.

18. Flower DR, Doytchinova IA: VaxiJen: a server for prediction of protective antigens, tumour antigens and subunit vaccines. BMC Bioinformatics 2007, 8(1):4.

19. Ashley, Heinson, Yawwani, Gunawardana, Bastiaan, Moesker, Carmen, Hume, Elena, Vataga: Enhancing the Biological Relevance of Machine Learning Classifiers for Reverse Vaccinology. International Journal of Molecular Sciences 2017, 18(2):312.

20. Dimitrov I, Zaharieva N, Doytchinova I: Bacterial Immunogenicity Prediction by Machine Learning Methods. Vaccines (Basel) 2020, 8(4).

21. Xu H, Zhao Z: NetBCE: An Interpretable Deep Neural Network for Accurate Prediction of Linear B-cell Epitopes. Genomics Proteomics Bioinformatics 2022, 20(5):1002–1012.

22. Collatz M, Mock F, Barth E, Holzer M, Sachse K, Marz M: EpiDope: a deep neural network for linear B-cell epitope prediction. Bioinformatics 2021, 37(4):448–455.

23. Chen HZ, Tang LL, Yu XL, Zhou J, Chang YF, Wu X: Bioinformatics analysis of epitope-based vaccine design against the novel SARS-CoV-2. Infect Dis Poverty 2020, 9(1):88.

24. Crooke SN, Ovsyannikova IG, Kennedy RB, Poland GA: Immunoinformatic identification of B cell and T cell epitopes in the SARS-CoV-2 proteome. Sci Rep 2020, 10(1):14179.

25. Grifoni A, Sidney J, Zhang Y, Scheuermann RH, Peters B, Sette A: A Sequence Homology and Bioinformatic Approach Can Predict Candidate Targets for Immune Responses to SARS-CoV-2. Cell Host Microbe 2020, 27(4):671–680.e672.

26. Kar T, Narsaria U, Basak S, Deb D, Castiglione F, Mueller DM, Srivastava AP: A candidate multi-epitope vaccine against SARS-CoV-2. Sci Rep 2020, 10(1):10895.

27. Zajac MD, Sangewar N, Lokhandwala S, Bray J, Sang H, McCall J, Bishop RP, Waghela SD, Kumar R, Kim T et al: Adenovirus-Vectored African Swine Fever Virus pp220 Induces Robust Antibody, IFN-gamma, and CTL Responses in Pigs. Front Vet Sci 2022, 9:921481.

28. Incorporating Machine Learning into Established Bioinformatics Frameworks. International Journal of Molecular Sciences 2021, 22(6):2903.

29. Li G, Iyer B, Prasath S, Ni Y, Salomonis N: DeepImmuno: Deep learning-empowered prediction and generation of immunogenic peptides for T cell immunity. bioRxiv : the preprint server for biology:2020.2012.2024.424262.

30. Usman M, Khan S, Lee JA: AFP-LSE: Antifreeze Proteins Prediction Using Latent Space Encoding of Composition of k-Spaced Amino Acid Pairs. Sci Rep 2020, 10(1):7197.

31. Al-Saggaf UM, Usman M, Naseem I, Moinuddin M, Jiman AA, Alsaggaf MU, Alshoubaki HK, Khan S: ECM-LSE: Prediction of Extracellular Matrix Proteins Using Deep Latent Space Encoding of k-Spaced Amino Acid Pairs. Front Bioeng Biotechnol 2021, 9:752658.

32. Usman M, Khan S, Park S, Lee JA: AoP-LSE: Antioxidant Proteins Classification Using Deep Latent Space Encoding of Sequence Features. Curr Issues Mol Biol 2021, 43(3):1489–1501.

33. Muhammad U, Khan S, Park S, Wahab A: AFP-SRC:identification of antifreeze proteins using sparse representation classifier. Neural Computing and Applications 2022, 34(3):2275–2285.

34. Yang B, Samantha S, Xiang Z, He Y: Protegen: a web-based protective antigen database and analysis system. Nucleic Acids Research 2011(suppl_1):D1073–D1078.

35. New quantitative descriptors of amino acids based on multidimensional scaling of a large number of physical–chemical properties. Molecular modeling annual 2001, 7(12):445–453.

36. Hellberg S, Sjoestroem M, Skagerberg B, Wold S: Peptide quantitative structure-activity relationships, a multivariate approach. Journal of Medicinal Chemistry 1987, 30(7):1126–1135.

37. Wold S, Jonsson J, Sjrstrm M, Sandberg M, Rnnar S: DNA and peptide sequences and chemical processes multivariately modelled by principal component analysis and partial least-squares projections to latent structures. Analytica Chimica Acta 1993, 277(2):239–253.

38. El-Manzalawy Y, Dobbs D, Honavar V: Predicting linear B-cell epitopes using string kernels. J Mol Recognit 2008, 21(4):243–255.

39. Peters B, Sidney J, Bourne P, Bui HH, Buus S, Doh G, Fleri W, Kronenberg M, Kubo R, Lund O et al: The immune epitope database and analysis resource: from vision to blueprint. PLoS Biol 2005, 3(3):e91.

40. Chen J, Liu H, Yang J, Chou KC: Prediction of linear B-cell epitopes using amino acid pair antigenicity scale. Amino Acids 2007, 33(3):423–428.

41. El-Manzalawy Y, Dobbs D, Honavar V: Predicting flexible length linear B-cell epitopes. Comput Syst Bioinformatics Conf 2008, 7:121–132.

42. Saha S, Raghava GPS: BcePred: Prediction of continuous B-cell epitopes in antigenic sequences using physico-chemical properties. In: ICARIS 2004, LNCS3239: 2004 2004. Springer: 197–204.

43. Saha S, Raghava GP: Prediction of continuous B-cell epitopes in an antigen using recurrent neural network. Proteins 2006, 65(1):40–48.

44. Dimitrov I, Naneva L, Doytchinova I, Bangov I: AllergenFP: allergenicity prediction by descriptor fingerprints. Bioinformatics 2014, 30(6):846–851.

45. Gupta S, Kapoor P, Chaudhary K, Gautam A, Kumar R, Open Source Drug Discovery C, Raghava GP: In silico approach for predicting toxicity of peptides and proteins. PLoS One 2013, 8(9):e73957.

