## Supplementary Materials for "VirusImmu: a novel ensemble machine learning approach for viral immunogenicity prediction"

Figure S1. Four-dimensional image of AUC value varying on different soft voting ensemble classifiers
Figure S2. ROC curves of all models
Figure S3. Measurements of antigenic B cell epitopes binding to antibody.

Table S1. Protein datasets:100 antigens (Positive set)
Table S2. Protein datasets:100 non-antigens (Negative set)
Table S3. Dataset of external evaluation
Table S4. The representation of twenty naturally occurring amino acids by Z-descriptors and E-descriptors
Table S5. List of epitopes for SARS-CoV-2 as given in the literature, and the immunogenicity prediction results by VirusImmu.
Table S6. B-cell linear epitopes predicted for ASFV pp220 protein.
Table S7. Eight antigenic and four non-antigenic B cell linear epitopes predicted by VirusImmu for ASFV pp220 protein.

S1. Materials and Methods
S2. A more specific of description of relevant models

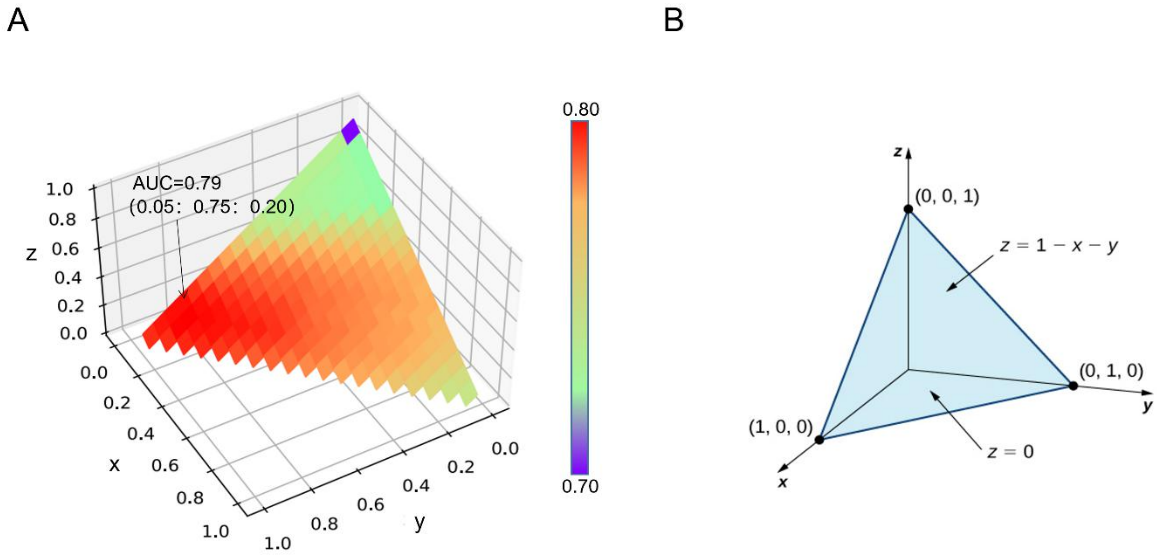


**Figure S1** (A) Four-dimensional image of AUC value varying on different soft voting ensemble classifiers. The principle of the 4-D image is the similar to the 3-D image, the X axis represents the weight of XGBoost model, the Y axis represents the weight of RF model, and the Z axis represents kNN model. The change in color as a fourth dimensional variable reflects the magnitude of the AUC value. The optimal soft voting ensemble classifier appears on the weight of 0.05: 0.75: 0.20 (XGBoost: RF: kNN), with the highest AUC value (0.793). (B) Since the weights of the three models always sum to 1 ($x + y + z = 1$), when they are used as independent variables of the four-dimensional image, the color changes reflecting the AUC values can only be displayed on one plane. This results in that, the four-dimensional image does not reflect the AUC changes as well as the three-dimensional image (Figure 2B), although all the variables are directly included.

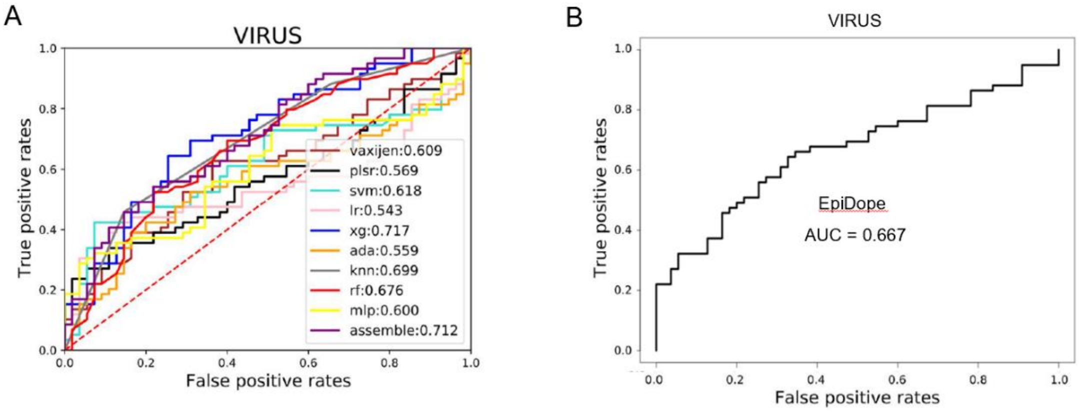


**Figure S2.** ROC curves of all models in present study, our optimal ensemble model, VaxiJen and EpiDope in external validation.

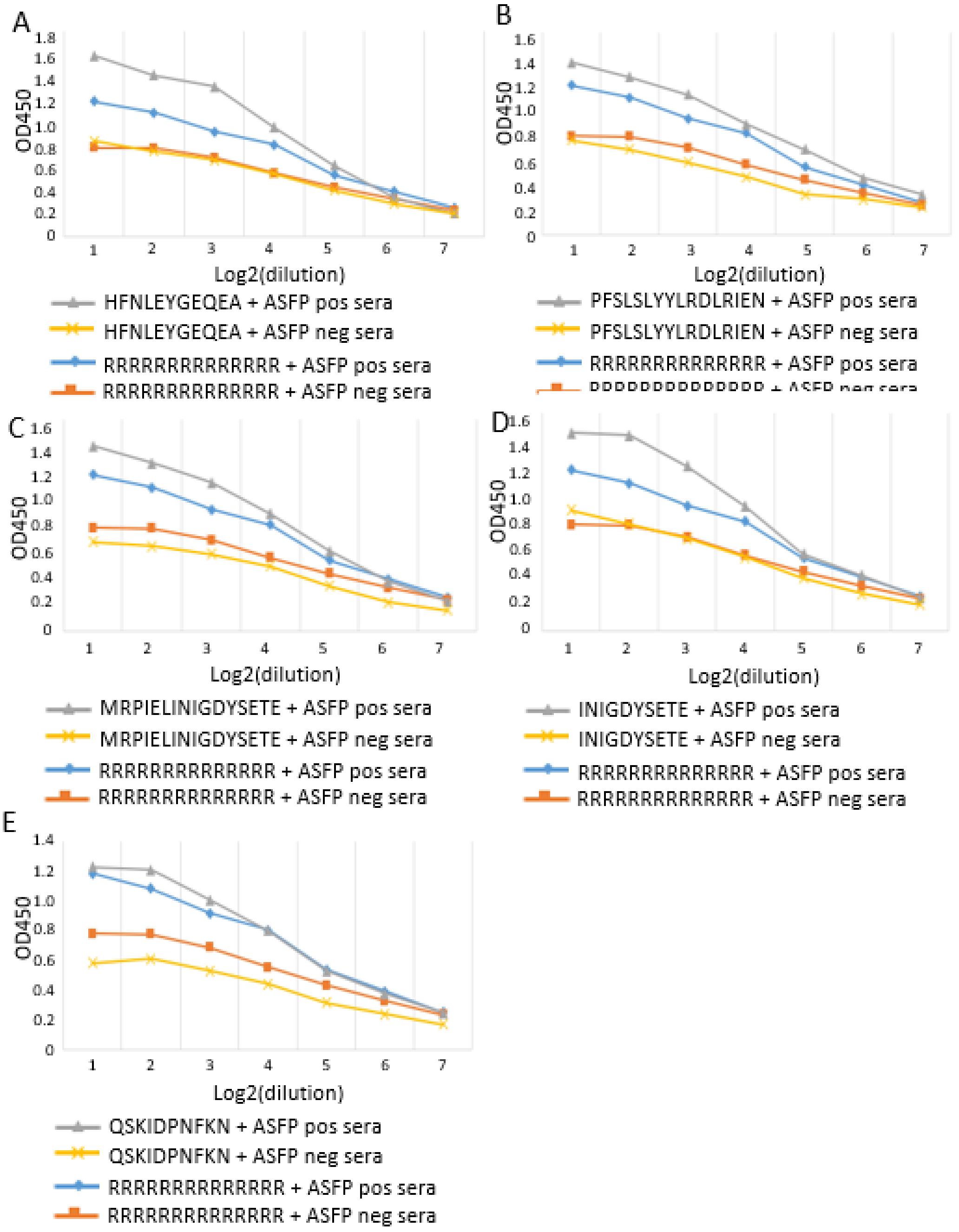


**Figure S3** Measurements of antigenic B cell epitopes binding to antibody. (A-E) The binding affinity assessed by ELISA between sera antibodies from pigs infected with ASFV SY18ΔMGF/ΔCD2v and linear B-cell epitopes of HFNLEYGEQEA (A), PFSLSLYYLRDLRIEN (B), MRPIELINIGDYSETE (C), INIGDYSETE (D) and QSKIDPNFKN (E).

**Table S1 Protein datasets:100 antigens (Positive set).**

| *Swiss-prot* | *Species* | *Protein* | *Reference* |
|---|---|---|---|
| P24030 | Aleutian mink disease parvovirus | Non-capsid protein NS-1 | Castelruiz et al., Vaccine, 2005, 23, 1225 |
| O12347 | Avian infectious bursal disease virus | Structural protein VP2 | Rong et al, Vaccine, 2005, 23, 4844 |
| Q5BMC7 | Avian orthoreovirus | Sigma C protein | Vasserman et al., Avian Dis., 2004, 48, 271 |
| P0C796 | Borna disease virus | Nucleoprotein | Henkel et al., J. Virol., 2005, 79, 314 |
| Q56I18 | Bovine ephemeral fever virus | Glycoprotein G | Walker, Curr. Top. Microbiol. Immunol., 2005, 292, 57 |
| P0CK29 | Bovine herpesvirus 1.1 | Glycoprotein D | Toussaint et al., Vet. Res., 2005, 36, 529 |
| P06167 | Bovine parainfluenza 3 virus | Hemagglutinin-neuraminidase | Schmidt et al., 2000, 74, 8922 |
| P09990 | Bovine parainfluenza 3 virus | Fusion glycoprotein | Schmidt et al., 2000, 74, 8922 |
| O10683 | Bovine respiratory syncytial virus | Major surface glycoprotein G | Brady et al., Vaccine, 22, 3762 |
| O10645 | Bovine viral diarrhea virus | Structural glycoprotein E2 | Liang et al., Vaccine, 2005, 23, 5252 |
| Q2LC28 | Classical swine fever virus | Envelope glycoprotein | Rau et al., Vet. Res., 2006, 37, 155 |
| Q68287 | Classical swine fever virus | NS3 polyprotein | Rau et al., Vet. Res., 2006, 37, 155 |
| Q07690 | Dengue virus type 2 | Envelope glycoprotein | Ocazionez et al., Vaccine, 2000, 19, 648 |
| O10247 | Dengue virus type 3 | Nonstructural protein 1 | Mellado-Sanchez et al., Viral Immunol., 2005, 18, 709 |
| Q2YH57 | Dengue virus type 3 | Envelope protein | Blaney et al., Am. J. Trop. Med. Hyg., 2004, 71, 811 |
| Q8V1K2 | Dengue virus type 3 | Premembrane | Blaney et al., Am. J. Trop. Med. Hyg., 2004, 71, 811 |
| O92944 | Dengue virus type 4 | Envelope protein | Guzman et al., Am. J. Trop. Med. Hyg., 2003, 69, 129 |
| Q04245 | Equid herpesvirus 1 | Glycoprotein D | Ruitenberg et al., Virus Res., 2001, 79, 125 |
| Q8AYT7 | Equine arteritis virus | GL envelope glycoprotein | Castillo-Olivares et al., J. Gen. Virol., 2001, 82, 2425 |
| O39843 | Foot-and-mouth disease virus | VP1 | Wong et al., Virology, 2000, 278, 27 |
| Q8JMH1 | Foot-and-mouth disease virus | 3A nonstructural protein | Grubman, Biologicals, 2005, 33, 227 |
| O92567 | Foot-and-mouth disease virus - type A | 3D protein | Garcia_Briones et al., Virology, 2004, 322, 264 |
| O89126 | Hepatitis C virus | E2 glycoprotein | Tarr et al., Hepatology, 2006, 43, 592 |

| | | | |
|---|---|---|---|
| P69618 | Hepatitis delta virus | Large delta antigen | Fiedler & Roggendorf, Intervirology, 2001, 44, 154 |
| O56048 | Hepatitis E virus | Capsid protein | Li et al., Vaccine, 2004, 22, 370 |
| Q19AX9 | Human coxsackievirus B3 | Capsid protein | Xu et al., Vaccine, 2004, 22, 3603 |
| Q6RXC2 | Human cytomegalovirus | UL123 | Wang et al., J. Clin. Virol., 2006, 35, 324 |
| Q6SW59 | Human cytomegalovirus | UL83 | Wang et al., J. Clin. Virol., 2006, 35, 324 |
| O56638 | Human cytomegalovirus | Glycoprotein B | Schleiss et al., J. Infect. Dis., 2003, 188, 1868 |
| Q6SW76 | Human cytomegalovirus | UL55 | Wang et al., J. Clin. Virol., 2006, 35, 324 |
| Q197M9 | Human enterovirus 71 | Structural protein VP1 | Wu et al., Vaccine, 2001, 20, 895 |
| Q6YFA9 | Human herpesvirus 1 | Glycoprotein D | BenMohamed et al., J. Virol., 2003, 77, 9463 |
| O37453 | Human herpesvirus 2 | Glycoprotein B | Heineman et al., Vaccine, 2004, 22, 2558 |
| Q5ICU7 | Human herpesvirus 2 | Glycoprotein D | Heineman et al., Vaccine, 2004, 22, 2558 |
| O40239 | Human immunodeficiency virus | Transmembrane glycoprotein gp41 | Decroix et al., Clin. Diagn. Lab. Immunol., 2003, 10, 1103 |
| O37033 | Human immunodeficiency virus 1 | Gag protein | Nabel, Vaccine, 2002, 20, 1945 |
| O09530 | Human immunodeficiency virus 1 | Gp120 | Parker et al., J. Virol., 2001, 75, 10906 |
| O36766 | Human immunodeficiency virus 1 | MA-p17 | Zimmerman et al., AIDS Res. Hum. Retroviruses, 1998, 14, 741 |
| Q14VH5 | Human immunodeficiency virus 1 | Pol protein | Nabel, Vaccine, 2002, 20, 1945 |
| O12164 | Human immunodeficiency virus type 1 | Envelope glycoprotein gp160 | Seaman et al., J. Virol., 2005, 79, 2956 |
| P04326 | Human immunodeficiency virus type 1 | Protein Tat | Partidos et al., Eur. J. Immunol., 2004, 34, 3723 |
| P04583 | Human immunodeficiency virus type 1 subtype A | Envelope glycoprotein gp160 | Wang et al., Virology, 2006, 350, 34 |
| P03378 | Human immunodeficiency virus type 1 subtype B | Envelope glycoprotein gp160 | Wang et al., Virology, 2006, 350, 34 |
| P04581 | Human immunodeficiency virus type 1 subtype D | Envelope glycoprotein gp160 | Wang et al., Virology, 2006, 350, 34 |
| Q1A2Y0 | Human metapneumovirus | Fusion protein | Tang et al., Vaccine, 2005, 23, 1657 |
| P03129 | Human papillomavirus type 16 | Protein E7 | Cheung et al., Vaccine, 2004, 23, 629 |
| P36731 | Human papillomavirus type 3 | Major capsid protein L1 | Stanley, Rev. Med. Virol., 2006, 16, 139 |
| P16071 | Human parainfluenza 1 virus | Hemagglutinin-neuraminidase | Skiadopoulos et al., 2002, 20, 1846 |
| O55887 | Human parainfluenza virus 1 | Fusion glycoprotein | Skiadopoulos et al., 2002, 20, 1846 |

| | | | |
|---|---|---|---|
| P89316 | Human parvovirus B19 | VP1 and VP2 structural protein | Franssila et al., Vaccine, 2004, 22, 3809 |
| O91261 | Human poliovirus 1 | VP1 protein | Suzuki, Gene, 2004, 328, 127 |
| Q91UA3 | Human poliovirus 1 | VP3-VP1 capsid protein | Suzuki, Gene, 2004, 328, 127 |
| P24569 | Human respiratory syncytial virus | Non-structural protein 2 | Jin et al., Vaccine, 2003, 21, 3647 |
| O09635 | Human respiratory syncytial virus | Fusion protein | Tang et al., J. Virol., 2004, 78, 11198 |
| Q4KRX1 | Human respiratory syncytial virus | Non-structural protein 1 | Jin et al., Vaccine, 2003, 21, 3647 |
| P69359 | Human respiratory syncytial virus B | Small hydrophobic protein | Jin et al., Vaccine, 2003, 21, 3647 |
| Q4KRW2 | Human respiratory syncytial virus B | Matrix protein 2-2 | Jin et al., Vaccine, 2003, 21, 3647 |
| P11198 | Human rotavirus A | Outer capsid protein VP4 | Banyai et al., J. Med. Virol., 2005, 76, 414 |
| P11231 | Human rotavirus A | RNA-binding protein VP2 | Ogier et al., Eur. J. Immunol., 2005, 35, 2122 |
| P25187 | Human rotavirus A | Glycoprotein VP7 | Banyai et al., J. Med. Virol., 2005, 76, 414 |
| P16487 | Human rotavirus A | VP6 protein | Ogier et al., Eur. J. Immunol., 2005, 35, 2122 |
| Q1KVI3 | Human SARS coronavirus | S1 protein | Cavanagh, Avian Pathol., 2003, 32, 567 |
| Q64FG1 | Human SARS coronavirus | S2 protein | Cavanagh, Avian Pathol., 2003, 32, 567 |
| P59595 | Human SARS coronavirus | Nucleocapsid protein | See et al., J. Gen. Virol., 2006, 87, 641 |
| Q67148 | Influenza A virus | M2 protein | Fan et al., Vaccine, 2004, 22, 2993 |
| P09345 | Influenza A virus | Hemagglutinin | Hoffmann et al., Proc. Natl. Acad. Sci. USA, 2005, 102, 12915 |
| O09754 | Japanese encephalitis virus | Envelope protein | Arroyo et al., J. Virol., 2004, 78, 12497 |
| P89481 | Japanese encephalitis virus | Pre-M protein | Wu et al., Microbiol. Immunol. , 2004, 48, 585 |
| P35256 | Lake Victoria marburgvirus | Membrane-associated structural protein VP24 | Warfield et al., Expert Rev. Vaccines, 2005, 4, 429 |
| P35258 | Lake Victoria marburgvirus | Minor nucleoprotein VP30 | Warfield et al., Expert Rev. Vaccines, 2005, 4, 429 |
| P35259 | Lake Victoria marburgvirus | Polymerase complex protein VP35 | Warfield et al., Expert Rev. Vaccines, 2005, 4, 429 |
| P35260 | Lake Victoria marburgvirus | Matrix protein VP40 | Warfield et al., Expert Rev. Vaccines, 2005, 4, 429 |
| Q38L42 | Lake Victoria marburgvirus | Glycoprotein | Jones et al., Nat. Med., 2005, 11, 786 |
| P17332 | Lassa virus | Glycoprotein polyprotein | Lukashevich et al., J. Virol., 2005, 79, 13934 |
| P04935 | Lassa virus | Nucleocapsid protein | Lukashevich et al., J. Virol., 2005, 79, 13934 |

| | | | |
|---|---|---|---|
| P35971 | Measles virus | Hemagglutinin glycoprotein | Putz et al., Vaccine, 2004, 22, 4173 |
| Q27YE8 | Mopeia virus | Z protein | Lukashevich et al., J. Virol., 2005, 79, 13934 |
| P11210 | Murine cytomegalovirus | Immediate-early protein 1 | Ye et al., J. Virol., 2002, 76, 2100 |
| Q29YA6 | Muscovy duck reovirus | Sigma C protein | Kuntz-Simon et al., Vaccine, 2002, 20, 3113 |
| Q91E46 | Muscovy duck reovirus | Sigma B protein | Kuntz-Simon et al., Vaccine, 2002, 20, 3113 |
| P12554 | Newcastle disease virus | Hemagglutinin-neuraminidase | Claassen et al., Pharmeuropa Bio., 2004, 2004, 1 |
| O57135 | Phocid herpesvirus 1 | Glycoprotein B | Martina et al., Vaccine, 2003, 21, 2433 |
| O57136 | Phocid herpesvirus 1 | Glycoprotein D | Martina et al., Vaccine, 2003, 21, 2433 |
| Q77Q74 | Porcine circovirus 2 | ORF-1 | Blanchard et al., Vaccine, 2003, 31, 4565 |
| Q9YQS7 | Porcine circovirus 2 | ORF-2 | Blanchard et al., Vaccine, 2003, 31, 4565 |
| P33484 | Porcine parvovirus | Coat protein VP1 | Roic et al., J. Vet. Med. B Infect. Dis. Vet. Public Health, 2006, 53, 17 |
| P17280 | Simian immunodeficiency virus | Protein Rev | Negri et al., J. Gen. Virol., 2004, 85, 1191 |
| P17282 | Simian immunodeficiency virus | Gag polyprotein | Nakaya et al., J. Virol., 2004, 78, 9366 |
| P17285 | Simian immunodeficiency virus | Protein Tat | Negri et al., J. Gen. Virol., 2004, 85, 1191 |
| P17664 | Simian immunodeficiency virus | Protein Nef | Negri et al., J. Gen. Virol., 2004, 85, 1191 |
| Q88229 | Sin Nombre virus | Envelope glycoprotein G1 | Bharadwaj et al., J. Gen. Virol., 2002, 83, 1745 |
| Q71TT1 | Vaccinia virus | A33R | Hooper et al., Virology, 2003, 306, 181 |
| Q71TT2 | Vaccinia virus | L1R | Hooper et al., Virology, 2003, 306, 181 |
| Q9DS89 | West Nile virus | Envelope protein | Calvert et al., J. Gen. Virol., 2006, 87, 339 |
| P89237 | Western equine encephalitis virus | Envelope glycoprotein E1 | Das et al., Antiviral Res., 2004, 64, 85 |
| O11459 | Zaire ebolavirus | Membrane-associated protein VP24 | Warfield et al., Expert Rev. Vaccines, 2005, 4, 429 |
| Q6V1Q9 | Zaire ebolavirus | Polymerase cofactor VP35 | Warfield et al., Expert Rev. Vaccines, 2005, 4, 429 |
| P87671 | Zaire ebolavirus | Envelope glycoprotein | Jones et al., Nat. Med., 2005, 11, 786 |
| Q2PDK5 | Zaire ebolavirus | Matrix protein VP40 | Warfield et al., Expert Rev. Vaccines, 2005, 4, 429 |
| Q05323 | Zaire ebolavirus | Minor nucleoprotein VP30 | Warfield et al., Expert Rev. Vaccines, 2005, 4, 429 |

**Table S2 Protein datasets：100 non-antigens (Negative set).**

| *Species* | *Protein* |
|---|---|
| Amsacta moorei entomopoxvirus | AMV011 |
| Amsacta moorei entomopoxvirus | AMV012 |
| Amsacta moorei entomopoxvirus | AMV026 |
| Amsacta moorei entomopoxvirus | AMV033 |
| Amsacta moorei entomopoxvirus | AMV077 |
| Amsacta moorei entomopoxvirus | AMV129 |
| Amsacta moorei entomopoxvirus | AMV142 |
| Amsacta moorei entomopoxvirus | AMV154 |
| Amsacta moorei entomopoxvirus | AMV220 |
| Amsacta moorei entomopoxvirus | AMV227 |
| Amsacta moorei entomopoxvirus | AMV252 |
| Avian pneumovirus | small hydrophobic protein |
| Avian pneumovirus | SH |
| Barmah Forest virus | 6K protein |
| Bovine parainfluenza virus | large polymerase subunit L |
| Bovine papular stomatitis virus | ORF001 hypothetical protein |
| Bovine papular stomatitis virus | ORF005 hypothetical protein |
| Bovine papular stomatitis virus | ORF008 ankyrin repeat protein |
| Bovine papular stomatitis virus | ORF010 EEV maturation protein |
| Bovine papular stomatitis virus | ORF031 hypothetical protein |
| Bovine papular stomatitis virus | ORF034 hypothetical protein |
| Bovine papular stomatitis virus | ORF044 virion core protein |
| Bovine papular stomatitis virus | ORF053 poly-A polymerase small subunit |

| | |
|---|---|
| Bovine papular stomatitis virus | ORF082 hypothetical protein |
| Bovine papular stomatitis virus | ORF116 hypothetical protein |
| Bovine papular stomatitis virus | ORF120 hypothetical protein |
| Convict Creek virus | RNA-dependent RNA polymerase |
| Camelpox virus | CMP6L |
| Camelpox virus | CMP8R |
| Camelpox virus | CMP121R |
| Camelpox virus | CMP126.5aR |
| Camelpox virus | CMP131L |
| Camelpox virus | CMP187R |
| Camelpox virus | hypothetical protein |
| Camelpox virus | hypothetical protein |
| Camelpox virus | putative IMV membrane morphogenesis protein |
| Camelpox virus | truncated kelch-like protein |
| Canarypox virus | CNPV092 conserved hypothetical protein |
| Canarypox virus | CNPV236 ribonucleotide reductase small subunit |
| Canarypox virus | CNPV315 G protein-coupled receptor-like protein |
| Cowpox virus | unknown |
| Cowpox virus | V100 |
| Cowpox virus | V150 |
| Cowpox virus | B10R protein |
| Dengue virus type 2 | unknown |
| Dugbe virus | L protein |
| Ectromelia virus | EVM071 |
| Ectromelia virus | EVM140 |
| Fowlpox virus | ORF FPV085 |

| | |
|---|---|
| Goatpox virus | unknown |
| Human parainfluenza virus 1 | matrix protein |
| Japanese encephalitis virus | Genome polyprotein |
| Japanese encephalitis virus | Structural polyprotein |
| Mayaro virus | protein nsP3 |
| Mayaro virus | Structural polyprotein |
| Montana myotis leukoencephalitis virus | non-structural protein NS5 |
| Molluscum contagiosum virus | MC030R |
| Molluscum contagiosum virus | MC061R |
| Molluscum contagiosum virus | MC070R |
| Molluscum contagiosum virus | MC099R |
| Molluscum contagiosum virus | MC104L |
| Molluscum contagiosum virus | MC113L |
| Molluscum contagiosum virus | MC114R |
| Molluscum contagiosum virus | MC138R |
| Molluscum contagiosum virus | MC139R |
| Molluscum contagiosum virus | MC141R |
| Molluscum contagiosum virus | MC146R |
| Molluscum contagiosum virus | MC147R |
| Molluscum contagiosum virus | MC162R |
| Monkeypox virus | Major core protein 4a precursor |
| Monkeypox virus | EEV maturation protein |
| Monkeypox virus | serine protease inhibitor-like SPI-1 protein |
| Melanoplus sanguinipes entomopoxvirus | ORF MSV001 62 amino acid repeat gene family protein |
| Melanoplus sanguinipes entomopoxvirus | ORF MSV083 hypothetical protein |
| Melanoplus sanguinipes entomopoxvirus | ORF MSV131 hypothetical protein |

| | |
|---|---|
| Melanoplus sanguinipes entomopoxvirus | ORF MSV234 hypthetical protein |
| Melanoplus sanguinipes entomopoxvirus | ORF MSV233 hypothetical protein |
| Melanoplus sanguinipes entomopoxvirus | ORF MSV240 leucine rich repeat gene family protein |
| Melanoplus sanguinipes entomopoxvirus | ORF MSV258 hypothetical protein |
| Mumps virus | P |
| Mumps virus | SH |
| Myxoma virus | m128L |
| Orf virus | ORF018 poly-A polymerase catalytic subunit PAPL |
| Orf virus | ORF097 DNA polymerase processivity factor |
| Orf virus | ORF116 hypothetical protein |
| Orf virus | ORF120 hypothetical protein |
| Orf virus | hypothetical protein |
| Orf virus | unnamed protein product |
| Orf virus | unnamed protein product |
| Orf virus | unknown |
| Orf virus | ORF028 hypothetical protein |
| Orf virus | ORF074 mRNA capping enzyme small subunit |
| Rabbitpox virus | RPV-017 |
| Rift Valley fever virus | Non-structural protein S, NSs |
| Rabbit fibroma virus | gp017L |
| Rabbit fibroma virus | gp073R |
| Rabbit fibroma virus | gp131R |
| Sheeppox virus | unknown |
| Swinepox virus | SPV049 hypothetical protein |
| Tamana bat virus | polyprotein |

**Table S3 Dataset of external evaluation: 59 antigens and 54 non-antigens. The length is the number of amino acids of every protein sequence.**

| ***Antigen*** | | | ***Non-antigen*** | | |
|---|---|---|---|---|---|
| ***swiss-prot*** | ***species*** | ***length*** | ***swiss-prot*** | ***species*** | ***length*** |
| P18541 | Lymphocytic choriomeningitis virus | 90 | P06821 | Influenza A virus | 97 |
| Q80928 | Human papillomavirus type 50 | 93 | Q8QV70 | Mumps virus | 171 |
| Q6IVU5 | Junin mammarenavirus | 94 | T1UCJ0 | Human metapneumovirus | 177 |
| A0A096Y4M5 | Peste-des-petits-ruminants virus | 99 | A0A4P2TYZ6 | Metapneumovirus sp | 179 |
| P06932 | Human papillomavirus 5 | 103 | A0A223FMT7 | Murmansk poxvirus | 209 |
| Q96702 | Cabbage leaf curl virus | 132 | A0A1P7U1P7 | Mumps orthorubulavirus | 224 |
| Q67618 | Tomato yellow leaf curl Sardinia virus | 134 | P0C998 | African swine fever virus | 246 |
| Q08590 | Indian cassava mosaic virus | 134 | A0A0C5AZI9 | African swine fever virus | 250 |
| A7LIS7 | Goatpox virus | 148 | A0A291NQV2 | Influenza A virus | 252 |
| Q77CG2 | Avian infectious bursal disease virus | 172 | A0A142CK82 | Brazilian marseillevirus | 270 |
| P03081 | Simian virus 40 | 174 | P23055 | Human parainfluenza 2 virus | 395 |
| A0MD33 | Porcine reproductive and respiratory syndrome virus | 183 | Q5UPG5 | Acanthamoeba polyphaga mimivirus | 421 |
| P0C567 | Murine polyomavirus | 195 | A0A0E3T7L7 | Bovine papular stomatitis virus | 426 |
| P0C6L3 | Hepatitis delta virus genotype I | 195 | A7IVP3 | Paramecium bursaria Chlorella virus | 472 |
| P09713 | Human cytomegalovirus | 199 | D3IZ23 | Pseudocowpox virus | 516 |
| K7XMM0 | Avian infectious bursal disease virus | 206 | A0A0A7MA42 | Parapoxvirus red deer | 521 |
| A8D6S6 | HBV | 212 | A0A1Z3GCX8 | Seal parapoxvirus | 523 |
| P0C573 | HBV | 212 | Q5UPJ9 | Acanthamoeba polyphaga mimivirus | 627 |
| P89951 | Gibbon hepatitis B virus subtype ayw3q | 212 | A0A097IVW4 | Cotia virus | 634 |
| P29996 | Hepatitis delta virus genotype I | 214 | A0A1V0QGS6 | Shearwaterpox virus | 635 |
| P09727 | Human cytomegalovirus | 215 | A0A3S7SWI8 | Moosepox virus GoldyGopher14 | 635 |
| A0A516L979 | Prokaryotic dsDNA virus sp | 225 | Q08FQ5 | Deerpox virus | 635 |

| | | | | | |
|---|---|---|---|---|---|
| P30019 | Hepatitis B virus genotype C subtype adr | 226 | A0A7D3V813 | Fadolivirus 1 | 640 |
| Q0PQY9 | Avian infectious bursal disease virus | 247 | Q5UPG7 | Acanthamoeba polyphaga mimivirus | 642 |
| J7K3A9 | Goose orthoreovirus | 321 | R4ZEN9 | Adoxophyes honmai entomopoxvirus 'L' | 645 |
| B0FDX7 | Orgyia leucostigma nucleopolyhedrovirus | 321 | R4ZEE7 | Mythimna separata entomopoxvirus 'L' | 646 |
| A0A1J0M549 | Avian orthoreovirus | 326 | O37319 | Heliothis armigera entomopoxvirus | 646 |
| Q8QV15 | Avian reovirus GEL03 97T | 326 | W6JPK4 | Anomala cuprea entomopoxvirus | 646 |
| Q83931 | Orthoreovirus S1 | 326 | Q9YW39 | Melanoplus sanguinipes entomopoxvirus | 647 |
| Q9J1B6 | ARV-176 | 326 | P24486 | Choristoneura biennis entomopoxvirus | 648 |
| Q4PIR7 | este-des-petits-ruminants virus | 335 | P29814 | Nucleoside triphosphatase I | 648 |
| P0DST1 | Variola virus | 354 | A0A1V0QGA0 | Shearwaterpox virus | 671 |
| O92921 | Hepatitis B virus genotype D | 389 | Q66810 | Tai Forest ebolavirus | 676 |
| P03077 | Murine polyomavirus | 421 | A0A6M6A941 | Orf virus | 718 |
| J9T568 | Kimberley virus | 434 | F1AWY5 | Orf virus | 723 |
| R4NFR6 | Influenza A virus | 465 | A0A2H4X2H1 | Flamingopox virus | 741 |
| Q6XT89 | Influenza A virus | 498 | D9J318 | Influenza A virus | 765 |
| Q8AZK7 | Epstein-Barr virus | 506 | A0A2H4UUX2 | Bodo saltans virus | 777 |
| P03524 | Rabies virus | 524 | E3T553 | Cafeteria roenbergensis | 777 |
| Q08823 | Peste-des-petits-ruminants virus | 525 | A0A6N1NK34 | Tupanvirus soda lake | 793 |
| Q4PIR6 | Peste-des-petits-ruminants virus | 546 | Q5UQU2 | Acanthamoeba polyphaga mimivirus | 817 |
| Q07F70 | Influenza A virus | 565 | Q5UPF8 | Acanthamoeba polyphaga mimivirus | 879 |
| V5R6W0 | Porcine parvovirus | 579 | Q87049 | Semliki forest virus | 1145 |
| A0A1D8QQE9 | Amdoparvovirus | 595 | Q9QSK4 | Invertebrate iridescent virus 6 | 1171 |
| A0A1S6JPN8 | Skunk amdoparvovirus | 641 | A0A7D5KF02 | Barmah forest virus | 1239 |
| A0A7L9B072 | Labrador amdoparvovirus 1 | 641 | A0A4Y5N1L5 | Semliki forest virus | 1253 |
| A0A0A0Q452 | Raccoon dog amdovirus | 641 | A0A0G2R660 | Middelburg virus | 1258 |
| Q89669 | Adelaide River virus | 660 | A0A4Y5MX58 | Semliki forest virus | 2432 |

| | | | | | |
|---|---|---|---|---|---|
| Q7T9D9 | Sudan ebolavirus | 676 | P08411 | Semliki forest virus | 2432 |
| P03070 | Simian virus 40 | 708 | A0A1J1DYJ4 | Getah virus | 2467 |
| Q80E47 | Tick-borne encephalitis virus | 721 | Q5Y389 | Getah virus | 2467 |
| P21428 | Influenza A virus | 759 | D7R979 | Chikungunya virus | 2474 |
| P16090 | Feline immunodeficiency virus | 856 | Q5XXP4 | Chikungunya virus | 2474 |
| P03203 | Epstein-Barr virus | 938 | A0A5P8KY96 | Ross River virus | 2493 |
| P03204 | Epstein-Barr virus | 992 | | | |
| P11223 | Avian infectious bronchitis virus | 1162 | | | |
| Q8JUX5 | Chikungunya virus | 1248 | | | |
| P0DTC2 | Severe acute respiratory syndrome coronavirus 2 | 1273 | | | |
| A0A513Q8L7 | Atypical porcine pestivirus | 1951 | | | |

**Table S4 The representation of twenty naturally occurring amino acids by Z-descriptors and E-descriptors.**

| *Amino acid* | *Z-descriptors* | | | *E-descriptors* | | | | |
|---|---|---|---|---|---|---|---|---|
| | ***Z1*** | ***Z2*** | ***Z3*** | ***E1*** | ***E2*** | ***E3*** | ***E4*** | ***E5*** |
| A | 0.07 | -1.73 | 0.09 | 0.01 | 0.13 | -0.48 | -0.04 | 0.18 |
| C | 0.71 | -0.97 | 4.13 | -0.13 | 0.17 | 0.07 | 0.57 | -0.37 |
| D | 3.64 | 1.13 | 2.36 | 0.30 | -0.06 | -0.01 | 0.23 | 0.16 |
| E | 3.08 | 0.39 | -0.07 | 0.22 | -0.28 | -0.32 | 0.16 | 0.30 |
| F | -4.92 | 1.30 | 0.45 | -0.33 | -0.02 | 0.07 | 0.00 | 0.21 |
| G | 2.23 | -5.36 | 0.30 | 0.22 | 0.56 | -0.02 | 0.02 | 0.11 |
| H | 2.41 | 1.74 | 1.11 | 0.02 | -0.18 | 0.04 | 0.28 | -0.02 |
| I | -4.44 | -1.68 | -1.03 | -0.35 | 0.07 | -0.09 | -0.20 | -0.11 |
| K | 2.84 | 1.41 | -3.14 | 0.24 | -0.34 | -0.04 | -0.33 | -0.03 |
| L | -4.19 | -1.03 | 0.98 | -0.27 | 0.02 | -0.27 | -0.27 | 0.21 |
| M | -2.49 | -0.27 | -0.41 | -0.24 | -0.14 | -0.16 | 0.32 | 0.08 |
| N | 3.22 | 1.45 | 0.84 | 0.26 | 0.04 | 0.12 | 0.12 | -0.06 |
| P | -1.22 | 0.88 | 2.23 | 0.17 | 0.29 | 0.41 | -0.22 | 0.38 |
| Q | 2.18 | 0.53 | -1.14 | 0.15 | -0.18 | -0.03 | 0.04 | -0.11 |
| R | 2.88 | 2.52 | -3.44 | 0.17 | -0.36 | 0.11 | -0.26 | -0.36 |
| S | 1.96 | -1.63 | 0.57 | 0.20 | 0.24 | -0.02 | -0.07 | -0.20 |
| T | 0.92 | -2.09 | -1.40 | 0.07 | 0.15 | -0.02 | -0.13 | -0.27 |
| V | -2.69 | -2.53 | -1.29 | -0.27 | 0.14 | -0.19 | -0.20 | -0.30 |
| W | -4.75 | 3.65 | 0.85 | -0.30 | -0.19 | 0.39 | 0.08 | 0.30 |
| Y | -1.39 | 2.32 | 0.01 | -0.14 | -0.06 | 0.43 | -0.10 | -0.09 |

**Table S5** List of epitopes for SARS-CoV-2 as given in the literature[1-5], and the immunogenicity prediction results by VirusImmu.

| Source | Coded Proteins | Epitopes | Immunogenicity |
|---|---|---|---|
| Chen et al. (2020) | Spike glycoprotein | LSPRWYFYY | 0.536598289 |
| | | RSRNSSRNS | 0.86297579 |
| | | IGYYRRATR | 0.656785306 |
| Crooke et al. (2020) | Membrane glycoprotein | EVTPSGTWL | 0.688347348 |
| | | KLDDKDPNFK | 0.490793404 |
| | | KTFPPTEPKKDKKKKADETQALPQ | 0.765427164 |
| Grifoni et al. (2020a) | Spike glycoprotein | DAVDCALDPLSETKCTLKSFTVEKGIYQTSN | 0.267523305 |
| | | VCGPKKSTNLVKNKCVNFNFNGLTGTGVLTESNKKFLPFQQF GRDIADTTDAVRDPQTLEILDITPCSFGGVSVI | 0.917424999 |
| | | GTNTSNQVAVLYQDVNCTEVPVAIHADQLTPTWRVYSTGS | 0.501377857 |
| Kar et al. (2020) | Spike glycoprotein | INITRFQTLLALHRS | 0.552521871 |
| | | GINITRFQTLLALHR | 0.666538778 |
| | | GWTFGAGAALQIPFA | 0.552509542 |
| | | FSYTESLAGKREMAII | 0.557510697 |
| | | HAGPGPGPY | 0.598804931 |
| | | KMGPGPGTRFA | 0.480148788 |

**Table S6. B-cell linear epitopes predicted for ASFV pp220 protein.**

| Method | Position | Epitope | AntigenicityScore (Vaxi |
|---|---|---|---|
| BCPred | 1390 | KLIKPPTDAGGRPLPGGELG | -0.1515 |
| | 1269 | TLFAWIVPYVGIPAGGGVRA | 0.2922 |
| | 607 | ARGHVGPNPGGAQEADWKAA | 0.9976 |
| | 361 | NSCAKKGGDEEKTPLDRRIE | 0.2712 |
| | 33 | RPFSGGGYFNGGGDKNPVQH | -0.2016 |
| | 1247 | SRVYPDPTTNATAPQDQNLV | 0.5635 |
| | 2124 | GSLYPTQFDYDEAGPGLAAG | 0.8026 |
| | 696 | SFPTSAGNDSNVFTDNAPAG | 0.0285 |
| | 1507 | SYWDNMAPRTYANVNDAANN | 0.4419 |
| | 1108 | EDPSLQQELGKVSYQELIRQ | 0.2541 |
| | 892 | GGADELEPEVIPEAAELYFR | 0.7627 |
| | 1969 | KIVSPKGQTRTLGSNRERER | 0.4212 |
| | 1056 | SDRFISPSTQPTPKWRPALY | 0.5392 |
| | 674 | GVRIIGRWFTEATGDTLAQV | -0.3466 |
| | 500 | FYTDIVQKKYGGGEDCECTR | 0.0099 |
| | 725 | IQQGRSVGTLRPVRASQAKN | 0.6678 |
| | 1443 | ANIQINNNPQPSERFEQYGR | 0.944 |
| | 2372 | ITEQNREEGPWSIVKQVGVG | 1.2156 |
| | 1819 | IENNEVYDPLLYPNLESGSP | 0.4663 |
| | 2205 | EVENMIQTPEIQNNPTPEVI | 0.3297 |
| | 2 | GNRGSSTSSRPPLSSEANLY | 0.559 |
| | 2272 | VIFNRHGIPVPHPRQILQTD | 0.3378 |
| | 2453 | GFSITGPSETFSDKQYDSDI | 0.4659 |
| | 90 | IEDILKDIKKQLPDPRAGST | 0.1879 |
| | 2050 | EVLNIPNRPPMNTREFMLKL | 0.3613 |
| | 136 | GQDKLIDTTDGAASICRQIV | 0.1939 |
| | 1083 | QTGMLQPNSQWDLVQKFRKQ | 0.5182 |
| | 870 | GIYDMFERPEPVYKLIPTRM | -0.0437 |
| | 2072 | NPYVSVSITQYGNELMSKGS | 0.6345 |
| | 2335 | PRWLGPSTGRVARAPVRMAP | 0.5537 |
| | 1946 | KTLQDVISFVESSYQEEQIN | 0.488 |
| | 1033 | MNQTNYSILPDEDGYTQSSQ | 0.382 |
| | 1218 | LIDWLGRKHGAISEIRNPGL | 0.5959 |
| | 1722 | LRGIGGVFRPNVTLMPLGDA | 0.4031 |
| | 1534 | GSDYGIQNNRSMMMVFNQLV | 0.4548 |
| | 243 | LTPTQKELDKLQTDEVDIIK | 0.2519 |
| | 1900 | VNIRSFKTVMMYNENTFGGV | 0.5249 |
| | 533 | SKAARSQVDLNQAINTFMYY | -0.0728 |
| | 1130 | NELKKEHTDKIQIVSKLIQG | -0.0456 |
| | 197 | MVTEVTKAAPNEEVINAVTM | 0.3472 |
| Bepipred2.0 (IEDB) | 5 | GSSTSSRPPLSSEANL | 0.4951 |
| | 27 | HIQRQTRPFSGGGYFNGGGDKNPVQHIKDYHIDSVSSKAK | 0.5211 |
| | 82 | FKVDTKQP | -0.0514 |
| | 99 | KQLPDPRAGSTFVKNAEKQET | 0.5117 |
| | 131 | EFIDLGQDKLIDTTD | 0.9415 |
| | 163 | HGLRAEYLDVHGSIENTLE | 0.5933 |
| | 204 | AAPNE | |
| | 225 | LNEQNL | 1.3889 |
| | 242 | ILTPTQKELDKLQTDEVD | 0.2097 |
| | 263 | LLNDTNSVLGTKN | 1.0422 |
| | 305 | GLKVEQYLQSKNWAEFDKELDLKRFSGLVSAENIA | 0.4944 |
| | 357 | KI | |
| | 364 | AKKGGDEEKTPLDRRIEAQRLDRK | 0.3924 |
| | 399 | QA | |
| | 421 | LTPNIT | 1.0008 |
| | 455 | AAREE | |
| | 477 | QSKIDPNFKN | 1.951 |
| | 506 | QKKYGGGEDC | -0.1514 |
| | 529 | ELGLSKAARSQV | 0.3624 |
| | 563 | THNKQEFQSYEEN | 0.5918 |
| | 590 | QLDTEKNARINSPAVDLARGHVGPNPGGAQEADWK | 0.6369 |
| | 635 | D | |
| | 659 | VNNIDSIQT | 0.4122 |
| | 695 | ESFPTSAGNDSNVFTDNAPAGHYYE | 0.0082 |
| | 726 | QQGRSVGTLRPVRASQAKN | 0.6278 |
| | 772 | MLGGEELRQMVPMSP | 0.7419 |
| | 805 | GLKNLNQSEIGGQRV | 0.9055 |
| | 824 | TPEEAAQ | 0.2741 |
| | 841 | DALSTRWET | 0.6191 |
| | 874 | MFERPEPVYKL | -0.0459 |
| | 894 | ADELEPEVIPE | 0.6488 |
| | 924 | SFRDENVQIS | 1.5241 |
| | 957 | INIGDYSETE | 1.7685 |
| | 980 | HFNLEYGEQEA | 1.6057 |
| | 1012 | TEWEK | |
| | 1019 | R | |
| | 1024 | ARTMNDFGMMNQTNYSILPDEDGYTQSSQLLPSDRFISPSTQPTPKWRPALYNIDSVDVQTGMLQPNSQWI | 0.639 |
| | 1102 | Q | |
| | 1104 | SEMFEDPSLQQELGKVSYQ | 0.4358 |
| | 1137 | TD | |
| | 1148 | QGSESLADTDV | 0.4412 |
| | 1187 | NNIKGLDLD | 1.4392 |
| | 1208 | TQAANVNRANL | 0.7069 |
| | 1220 | DWLGRKHGAISEIRNPGLVVK | 0.8265 |
| | 1244 | VRLSRVYPDPTTNATAPQDQNL | 0.6182 |
| | 1281 | PAGGGVRAEQEL | 1.0413 |
| | 1298 | VD | |
| | 1323 | QVRFPETSTA | 0.5477 |
| | 1361 | PHIDKNIIQYYENRSNPG | -0.1056 |
| | 1389 | DKLIKPPTDAGGRPLPGGELGLEG | -0.0325 |
| | 1430 | VLQLRGGVQRRDAANIQINNNPQPSERFEQY | 0.779 |
| | 1474 | ENNSGLR | -0.2482 |
| | 1492 | SNLIRTNNAQEENTLSYWDNMAPRTYANVNDAANNLRRYRLYGSDYGIQNN | 0.281 |
| | 1562 | DAPS | |
| | 1578 | NGNFSQAVMELGYTHPDLARDNIAFGHRGDPTE | 0.9657 |
| | 1630 | NRQGLS | 0.6293 |
| | 1650 | KENYRANLP | 0.5371 |
| | 1680 | YTNVQLARPN | 0.414 |
| | 1706 | NNINVPMTGLSVGQA | 0.7042 |
| | 1723 | R | |
| | 1726 | GGV | |

| | | | |
|---|---|---|---|
| | 1738 | LGDAQNNTSDV | 0.769 |
| | 1763 | GSHTLA | -0.0932 |
| | 1785 | LETEEHFIQ | 0.1707 |
| | 1796 | MSRYNKEPL | 0.8388 |
| | 1816 | DLRIENNEVYDPLLYPNLESGSPEF | 0.7857 |
| | 1843 | LYGTRKLLGNDPVQLSDMPG | 0.0695 |
| | 1874 | VVAREQITPTR | 0.9289 |
| | 1913 | ENTFGG | 0.726 |
| | 1923 | SENRDD | 1.0067 |
| | 1960 | QEEQINHIHKIVSPKGQTRTLGSNRERERIFNLFDM | 0.4142 |
| | 2010 | PLANIYNYDYSF | 0.9465 |
| | 2033 | AEKVRSLNTAAPQPDIA | 0.4217 |
| | 2051 | VLNIPNRPPMNTR | 0.6027 |
| | 2084 | NELMSKGSAGYM | 0.3108 |
| | 2098 | IFRGDNALNMGRPKFLSD | 0.1409 |
| | 2127 | YPTQFDYDEAGPG | 0.7763 |
| | 2148 | REQWGQPLSE | -0.4091 |
| | 2200 | VS | |
| | 2204 | REVENMIQTPEIQNNPT | 0.27 |
| | 2229 | NWTQQY | 0.2473 |
| | 2247 | NIGQPNS | -0.0175 |
| | 2277 | HGIPVPHPRQILQTDDEAT | 0.2983 |
| | 2333 | N | |
| | 2335 | PRWLGPSTGRVARAPVRMAPRDMRHPISYTE | 0.6962 |
| | 2374 | EQNREEGPWSIVKQVGVGIQKPTLVQIG | 0.949 |
| | 2403 | DR | |
| | 2432 | SQFRNVLVSPDHIINPSITEYGFSITGPSETFSDKQYDSD | 0.4544 |
| | 1 | MGNRGSSTSSRPPLSSE | 0.7977 |
| | 29 | QRQTRPFSGGGYFNGGGDKNPVQ | 0.1615 |
| | 61 | VSS | |
| | 85 | DTKQPI | -0.2417 |
| | 99 | KQLPDPRAGSTFVKNAEKQE | 0.5396 |
| | 140 | LIDTTDGAA | 0.7788 |
| | 175 | S | |
| | 177 | EN | |
| | 202 | TKAAPNEEV | 0.345 |
| | 244 | TPTQKELDKLQT | 0.1358 |
| | 269 | SVL | |
| | 317 | WAE | |
| | 363 | CAKKGGDEEKTPLDR | 0.2361 |
| | 398 | TQA | |
| | 454 | AAAREE | 1.0445 |
| | 477 | QSKIDPN | 1.3053 |
| | 507 | KKYGGGEDCECT | -0.1372 |
| | 535 | A | |
| | 537 | R | |
| | 539 | QV | |
| | 564 | H | |
| | 567 | QEFQSYEEN | 0.2488 |
| | 594 | EKNARINSP | 0.1297 |
| | 604 | VD | |
| | 609 | GHVGPNPGGAQEADWK | 1.2366 |
| | 683 | TEATGDT | -0.1499 |
| | 696 | SFPTSAGNDSNVFTDNAPAGHYY | -0.0083 |
| | 724 | EIQQGRSVG | 0.8322 |
| | 736 | PVRASQAK | 0.036 |
| | 811 | QSEIGG | 0.9251 |
| | 822 | ARTPEEAA | 0.2168 |
| | 844 | STRWE | |
| | 876 | ERPEPV | 1.189 |
| | 894 | ADELEPEVIPE | 0.6488 |
| | 961 | DYSET | |
| | 985 | YGEQEAT | 0.6934 |
| | 1025 | RT | |
| | 1039 | SILPDEDGYTQSSQLL | -0.0718 |
| | 1059 | FISPSTQPTPKWRP | 0.931 |
| | 1080 | VDV | |
| | 1086 | MLQPNSQW | 0.3992 |
| | 1107 | FEDPSLQQ | 0.4423 |
| | 1133 | KKEHT | |
| | 1151 | ESLADT | 0.1761 |
| | 1209 | QAANV | |
| | 1227 | G | |
| | 1231 | EI | |
| | 1238 | VV | |
| | 1249 | VYPDPTTNATAPQDQNL | 0.598 |
| | 1279 | GIPAGGGVRAE | 0.8753 |
| | 1327 | PETSTA | 0.1159 |
| Bepipred (IEDB) | 1372 | ENRSNPG | 1.2205 |
| | 1392 | IKPPTDAGGRPLPGGELGL | 0.172 |
| | 1437 | VQRRDA | 1.4034 |
| | 1445 | IQINNNPQPSERFE | 1.0219 |
| | 1474 | ENN | |
| | 1478 | GL | |
| | 1498 | NNAQEENTLS | 0.3815 |
| | 1511 | NMAPRTYANVNDAAN | 0.6209 |
| | 1537 | YGIQ | |
| | 1562 | DAPSG | |
| | 1581 | F | |
| | 1591 | T | |
| | 1593 | PDLA | |
| | 1603 | GHRGDPTEQ | 0.948 |
| | 1629 | TNRQG | |
| | 1709 | N | |
| | 1739 | GDAQNNTSDV | 0.5538 |
| | 1766 | TL | |
| | 1798 | RYNKE | |
| | 1804 | L | |
| | 1822 | NEV | |
| | 1833 | LESGSP | 0.4205 |
| | 1849 | LL | |
| | 1852 | N | |
| | 1854 | PVQLSDM | 1.5975 |

| | | | |
|---|---|---|---|
| | 1875 | V | |
| | 1879 | QITPT | |
| | 1915 | TF | |
| | 1924 | ENRDDKPI | 1.1109 |
| | 1958 | SYQEE | |
| | 1972 | SPKGQTRTLGSNRE | 0.3788 |
| | 2040 | NTAAPQPDIA | 0.1903 |
| | 2054 | IPNRPPMN | 0.0369 |
| | 2086 | LMSK | |
| | 2091 | SAG | |
| | 2104 | ALNM | |
| | 2112 | F | |
| | 2128 | PTQFDYDEAGPGLAAG | 0.7719 |
| | 2145 | QRGREQWGQPLSE | -0.0168 |
| | 2210 | IQTPEIQNNPTPEVIA | 0.3972 |
| | 2229 | NWTQQ | |
| | 2235 | R | |
| | 2248 | IGQPN | |
| | 2279 | IPVPHPRQILQTDDEAT | 0.1942 |
| | 2315 | D | |
| | 2337 | WLGPSTGRVAR | 0.4012 |
| | 2350 | VRMAPRDMRHPISYT | 1.1403 |
| | 2374 | EQNREEGPWS | 1.1622 |
| | 2391 | GIQ | |
| | 2403 | DR | |
| | 2444 | IIN | |
| | 2449 | IT | |
| | 2454 | FSITGPSETFSDKQYD | 0.6327 |
| Emini Surface Accessibility Prediction (IEDB) | 4 | RGSSTSSRPPL | 0.6741 |
| | 27 | HIQRQTRP | 1.1418 |
| | 44 | GGDKNP | -0.0492 |
| | 84 | VDTKQP | 0.2771 |
| | 96 | DIKKQLPDPRA | 0.3863 |
| | 223 | RLLNEQN | -0.1264 |
| | 245 | PTQKELDKLQ | 0.1381 |
| | 308 | VEQYLQSKNW | 0.2682 |
| | 319 | EFDKELD | -0.24 |
| | 350 | QTFNERHK | 1.0772 |
| | 366 | KGGDEEKTPLDRRIE | 0.5723 |
| | 382 | QRLDRK | 0.0928 |
| | 396 | KSTQAY | 0.0403 |
| | 454 | AAAREERET | 1.0504 |
| | 476 | EQSKIDPNFKNL | 1.7565 |
| | 505 | VQKKYG | 0.7456 |
| | 561 | NLTHNKQEFQSYEEN | 0.6851 |
| | 591 | LDTEKNARIN | 0.4422 |
| | 633 | EYDVKRRF | -0.0419 |
| | 740 | SQAKNI | 0.4954 |
| | 823 | RTPEEA | 0.1105 |
| | 845 | TRWETE | -0.0813 |
| | 874 | MFERPEPV | 0.2761 |
| | 962 | YSETEI | 1.0271 |
| | 984 | EYGEQEATK | 0.4015 |
| | 1011 | RTEWEKF | -1.0696 |
| | 1043 | DEDGYTQS | -0.7995 |
| | 1062 | PSTQPTPKWR | 0.8149 |
| | 1098 | KFRKQL | -0.5378 |
| | 1131 | ELKKEHTD | 0.9773 |
| | 1249 | VYPDPTTNATAPQDQ | 0.408 |
| | 1325 | RFPETS | 0.6457 |
| | 1370 | YYENRSN | 0.8193 |
| | 1436 | GVQRRD | 2.3403 |
| | 1448 | NNNPQPSERFEQY | 0.6385 |
| | 1472 | ALENNS | 0.2498 |
| | 1497 | TNNAQEENT | 0.4216 |
| | 1512 | MAPRTY | 0.792 |
| | 1526 | NLRRYRLY | -0.3504 |
| | 1538 | GIQNNR | 2.5596 |
| | 1605 | RGDPTE | 0.4403 |
| | 1628 | DTNRQG | -0.5434 |
| | 1648 | YLKENYRA | 0.9547 |
| | 1795 | YMSRYNKEP | 0.8402 |
| | 1943 | SLRKTL | -0.2707 |
| | 1957 | SSYQEEQ | 0.7061 |
| | 1973 | PKGQTR | 1.5669 |
| | 1980 | LGSNRERER | 1.2267 |
| | 2015 | YNYDYS | 1.2727 |
| | 2042 | AAPQPD | 0.2648 |
| | 2055 | PNRPPMNTR | 0.5485 |
| | 2129 | TQFDYDE | 0.8119 |
| | 2145 | QRGREQW | 0.9496 |
| | 2213 | PEIQNNPT | 0.7522 |
| | 2229 | NWTQQYRAR | 0.669 |
| | 2353 | APRDMR | 1.2441 |
| | 2372 | ITEQNREEG | 1.4008 |
| | 2461 | ETFSDKQYDS | 0.5386 |
| | 21 | YAKLQDH | 0.6546 |
| | 50 | VQHIKDY | 0.3823 |
| | 58 | IDSVSSK | 1.0029 |
| | 66 | KLRVIEG | -0.1782 |
| | 118 | ETVCKMI | -0.117 |
| | 148 | ASICRQIVLYINS | -0.1431 |
| | 166 | RAEYLDVHGS | 0.8299 |
| | 216 | MIEAVYRR | 0.6217 |
| | 258 | VDIIKL | -0.0867 |
| | 277 | GKVLSYTLCNLGIAASVA | 0.5893 |
| | 301 | LQKVGLKVEQYLQ | 0.7319 |
| | 329 | FSGLVSAE | 0.6821 |
| | 344 | AVNLLRQ | 0.3037 |
| | 410 | KIGIKLVKEIA | 0.9929 |
| | 442 | ALDLSLIGF | 1.4726 |
| | 466 | QFMLVKNV | 0.0219 |
| | 488 | YDSCSRLLQIIDFYTDIVQKK | 0.0253 |

| | | | |
|---|---|---|---|
| Kolaskar & Tongaonkar Antigenicity (IEDB) | 515 | CECTRV | -0.311 |
| | 522 | GAALTVEEL | 0.7204 |
| | 537 | RSQVDLN | 0.716 |
| | 551 | YYYYVAQIYSN | 0.1236 |
| | 577 | ATILGDA | -0.1752 |
| | 601 | SPAVDLARGH | 0.6259 |
| | 624 | KAAVSAIELEYD | 0.9439 |
| | 646 | GLDLYLK | 1.1805 |
| | 665 | IQTVQQML | 0.0686 |
| | 690 | LAQVFES | -0.373 |
| | 715 | GHYYEKVAAE | 0.4755 |
| | 731 | VGTLRPVRAS | 0.8587 |
| | 783 | PMSPLQIYKTLLEYIQHSALSVGLK | 0.6367 |
| | 817 | QRVALA | 1.0292 |
| | 830 | QRVYLSTVRVNDA | 0.4082 |
| | 864 | KIFIVLGIY | 0.6751 |
| | 878 | PEPVYKLI | -0.1473 |
| | 899 | PEVIPEAAELYFRLPRLAEFYQKLF | 0.1131 |
| | 943 | SGLIRI | -0.6259 |
| | 973 | EINVIYQHFN | 0.7566 |
| | 993 | KALIHFV | 0.1328 |
| | 1018 | QRIVQE | -1.1031 |
| | 1050 | SSQLLPSD | 0.1755 |
| | 1074 | LYNIDSVDVQ | 1.4288 |
| | 1093 | WDLVQKF | 0.1458 |
| | 1113 | QQELGKVSYQELIRQ | 0.2166 |
| | 1140 | IQIVSKLI | 0.6004 |
| | 1160 | KIFLFHETVITGLNLLSAIYVLLN | 0.2521 |
| | 1235 | PGLVVKE | 2.0531 |
| | 1246 | LSRVYPD | -0.1448 |
| | 1271 | FAWIVPYVGIP | 0.359 |
| | 1293 | AARYLVDN | -0.9954 |
| | 1304 | MQLLLTN | 0.4742 |
| | 1321 | MVQVRFP | 0.7158 |
| | 1332 | AQVHLDFTGLISLIDS | 0.8828 |
| | 1355 | FLNLLRPH | -0.2193 |
| | 1388 | IDKLIKP | -1.101 |
| | 1419 | KTYILLTKPYNVLQLRG | 0.3705 |
| | 1460 | YGRVFSRLVFYD | 0.1959 |
| | 1480 | RVEQVVLGDF | 0.8466 |
| | 1549 | FNQLVASYIARFY | -0.3525 |
| | 1566 | GKIYLNLINA | 0.1207 |
| | 1582 | SQAVMEL | 0.393 |
| | 1611 | QSVLLLSLGLMLQRL | 0.8232 |
| | 1634 | LSQHLISTLTEIPIYLK | 0.3569 |
| | 1665 | NILISQGELLKQFI | -0.4507 |
| | 1680 | YTNVQLAR | 0.5932 |
| | 1747 | DVVRKRLVAVIDG | 0.1364 |
| | 1772 | MEVLHELTDHPIYL | 0.2951 |
| | 1805 | MPFSLSLYYLRD | 1.2015 |
| | 1825 | YDPLLYPN | 0.4206 |
| | 1840 | FKLLYGTR | 0.6424 |
| | 1852 | NDPVQLS | 0.8431 |
| | 1872 | ETVVARE | 0.3788 |
| | 1888 | FYTHAIQALRFIVNIR | 0.3366 |
| | 1941 | VYSLRK | 0.9265 |
| | 1948 | LQDVISFVESS | 0.3377 |
| | 1965 | NHIHKIVSPK | -0.5117 |
| | 2007 | RSIPLANI | 1.3938 |
| | 2022 | EEIACLMYG | 1.5397 |
| | 2048 | IAEVLNI | 0.6255 |
| | 2067 | LKLLINPYVSVSIT | 0.6421 |
| | 2118 | FNKVLFGSLYPTQ | 0.2641 |
| | 2160 | NQALHELVRTI | -0.5558 |
| | 2173 | PQKLRVLRNI | -0.7875 |
| | 2221 | PEVIAA | 0.496 |
| | 2238 | VDTLIN | -0.5749 |
| | 2256 | DLIQTITPVTVRAQLGVIFN | 0.6296 |
| | 2278 | GIPVPHPRQI | 0.6701 |
| | 2304 | NIPAIIM | 0.2332 |
| | 2327 | LERYVFNVP | -0.7904 |
| | 2346 | ARAPVRM | 0.2801 |
| | 2382 | WSIVKQVGVGIQKPTLVQI | 0.646 |
| | 2420 | QRLLRLR | -1.0025 |
| | 2434 | FRNVLVSPD | 0.4284 |
| AAP | 2250 | QPNSMLDLIQTITPVTVRAQ | 0.63 |
| | 2135 | EAGPGLAAGIQRGREQWGQP | 0.5519 |
| | 209 | EVINAVTMIEAVYRRLLNEQ | -0.0277 |
| | 683 | TEATGDTLAQVFESFPTSAG | 0.0273 |
| | 1970 | IVSPKGQTRTLGSNRERERI | 0.5139 |
| | 2339 | GPSTGRVARAPVRMAPRDMR | 0.8375 |
| | 612 | GPNPGGAQEADWKAAVSAIE | 0.6723 |
| | 731 | VGTLRPVRASQAKNIRDLIG | 0.2906 |
| | 1238 | VVKENDVRLSRVYPDPTTNA | 0.5215 |
| | 878 | PEPVYKLIPTRMILGGADEL | 0.055 |
| | 44 | GGDKNPVQHIKDYHIDSVSS | 0.5687 |
| | 1843 | LYGTRKLLGNDPVQLSDMPG | 0.0695 |
| | 356 | HKILENSCAKKGGDEEKTPL | 0.0653 |
| | 1266 | VTETLFAWIVPYVGIPAGGG | 0.2171 |
| | 1125 | IRQAINELKKEHTDKIQIVS | 0.0729 |
| | 2277 | HGIPVPHPRQILQTDDEATQ | 0.2768 |
| | 1393 | KPPTDAGGRPLPGGELGLEG | 0.2797 |
| | 1495 | IRTNNAQEENTLSYWDNMAP | 0.5239 |
| | 1066 | PTPKWRPALYNIDSVDVQTG | 1.0301 |
| | 1434 | RGGVQRRDAANIQINNNPQP | 0.9754 |
| | 1819 | IENNEVYDPLLYPNLESGSP | 0.4663 |
| | 774 | GGEELRQMVPMSPLQIYKTL | 0.4203 |
| | 704 | DSNVFTDNAPAGHYYEKVAA | -0.0267 |
| | 1702 | IYYNNNINVPMTGLSVGQAA | 0.6286 |
| | 983 | LEYGEQEATKKALIHFVNEI | 0.4796 |
| | 2373 | TEQNREEGPWSIVKQVGVGI | 1.1333 |
| | 1360 | RPHIDKNIIQYYENRSNPGS | -0.3694 |
| | 2194 | TIREQLVSMRREVENMIQTP | 0.181 |

| | | | |
|---|---|---|---|
| | 487 | LYDSCSRLLQIIDFYTDIVQ | -0.2392 |
| | 2217 | NNPTPEVIAAAQNWTQQYRA | 0.3433 |
| | 823 | RTPEEAAQRVYLSTVRVNDA | 0.2166 |
| | 1730 | RPNVTLMPLGDAQNNTSDVV | 0.756 |
| | 2310 | MTPFTDLANDLRTFLETLER | -0.2917 |
| | 85 | DTKQPIEDILKDIKKQLPDF | -0.2025 |
| | 563 | THNKQEFQSYEENYATILGD | 0.4338 |
| | 1305 | QLLLTNIFEMTSSFNKMVQV | -0.081 |
| | 447 | LIGFYTNAAAREERETFLTQ | 0.4256 |
| | 2071 | INPYVSVSITQYGNELMSKG | 0.622 |
| | 2040 | NTAAPQPDIAEVLNIPNRPP | 0.3278 |
| | 1198 | QKSIIEWLRETQAANVNRAN | 0.4038 |
| | 960 | GDYSETEIRQLIKEINVIYQ | 0.3408 |
| | 1104 | SEMFEDPSLQQELGKVSYQE | 0.4148 |
| | 1871 | NETVVAREQITPTRFEHFYT | 0.6833 |
| | 1899 | IVNIRSFKTVMMYNENTFGG | 0.415 |
| | 251 | DKLQTDEVDIIKLLNDTNSV | -0.1093 |
| | 534 | KAARSQVDLNQAINTFMYYY | -0.1248 |
| | 1275 | VPYVGIPAGGGVRA | 0.4287 |
| | 1393 | KPPTDAGGRPLPGG | -0.1806 |
| | 361 | NSCAKKGGDEEKTP | 0.2924 |
| | 37 | GGGYFNGGGDKNPV | -0.3047 |
| | 1247 | SRVYPDPTTNATAP | 0.5071 |
| | 608 | RGHVGPNPGGAQEA | 0.7235 |
| | 702 | GNDSNVFTDNAPAG | -0.1909 |
| | 894 | ADELEPEVIPEAAE | 0.5688 |
| | 2130 | QFDYDEAGPGLAAG | 0.6084 |
| | 1953 | SFVESSYQEEQINH | 0.7172 |
| | 1514 | PRTYANVNDAANNL | 0.2121 |
| | 5 | GSSTSSRPPLSSEA | 0.4551 |
| | 263 | LLNDTNSVLGTKNF | 0.8925 |
| | 1827 | PLLYPNLESGSPEF | 0.7991 |
| | 136 | GQDKLIDTTDGAAS | 0.5063 |
| | 1110 | PSLQQELGKVSYQE | 0.7288 |
| | 2053 | NIPNRPPMNTREFM | 0.2817 |
| | 1726 | GGVFRPNVTLMPLG | 0.498 |
| | 1498 | NNAQEENTLSYWDN | 0.3819 |
| | 466 | QFMLVKNVLEEQSK | 0.0186 |
| | 2374 | EQNREEGPWSIVKQ | 0.8912 |
| | 1443 | ANIQINNNPQPSER | 1.2349 |
| | 1060 | ISPSTQPTPKWRPA | 1.0583 |
| | 2258 | IQTITPVTVRAQLG | 0.844 |
| | 728 | GRSVGTLRPVRASQ | 0.6941 |
| | 1076 | NIDSVDVQTGMLQP | 1.1248 |
| | 1556 | YIARFYDAPSGKIY | -0.0762 |
| | 1601 | AFGHRGDPTEQSVL | 0.9612 |
| | 394 | LNKSTQAYNDFLEN | -0.1801 |
| | 1130 | NELKKEHTDKIQIV | 0.0326 |
| | 1873 | TVVAREQITPTRFE | 1.1362 |
| | 2453 | GFSITGPSETFSDK | 0.2791 |
| | 2145 | QRGREQWGQPLSEY | -0.0733 |
| | 244 | TPTQKELDKLQTDE | -0.0763 |
| | 631 | ELEYDVKRRFYRAL | -0.2186 |
| | 2331 | VFNVPRWLGPSTGR | -0.3108 |
| FBCPred | 2276 | RHGIPVPHPRQILQ | 0.308 |
| | 1468 | VFYDALENNSGLRV | -0.2831 |
| | 2010 | PLANIYNYDYSFEE | 0.7738 |
| | 98 | KKQLPDPRAGSTFV | 0.6067 |
| | 1534 | GSDYGIQNNRSMMM | 0.701 |
| | 837 | VRVNDALSTRWETE | 0.7386 |
| | 314 | SKNWAEFDKELDLK | 0.8265 |
| | 679 | GRWFTEATGDTLAQ | -0.4725 |
| | 876 | ERPEPVYKLIPTRM | 0.1807 |
| | 1975 | GQTRTLGSNRERER | 0.5163 |
| | 916 | AEFYQKLFSFRDEN | 0.6364 |
| | 1180 | VLLNNFRNNIKGLD | 0.4792 |
| | 1409 | GLEGVNQIINKTYI | 0.359 |
| | 2243 | NFIGNIGQPNSMLD | -0.0788 |
| | 507 | KKYGGGEDCECTRV | -0.0761 |
| | 53 | IKDYHIDSVSSKAK | 1.29 |
| | 1039 | SILPDEDGYTQSSQ | -0.0341 |
| | 1373 | NRSNPGSFYWLEEH | 0.5856 |
| | 2081 | QYGNELMSKGSAGY | 0.4261 |
| | 1759 | GIIRGSHTLADSAM | 0.1664 |
| | 1782 | PIYLETEEHFIQNY | 0.2029 |
| | 1427 | PYNVLQLRGGVQRR | 0.5631 |
| | 1232 | IRNPGLVVKENDVR | 0.7565 |
| | 1991 | NLFDMNIIPINVNA | 1.041 |
| | 563 | THNKQEFQSYEENY | 0.6232 |
| | 1743 | NNTSDVVRKRLVAV | 0.4818 |
| | 194 | HERMVTEVTKAAPN | 0.3655 |
| | 2306 | PAIIMTPFTDLAND | 0.1772 |
| | 2100 | RGDNALNMGRPKFL | 0.6252 |
| | 1694 | LGANNDSVIYYNNN | 0.3964 |
| | 1315 | TSSFNKMVQVRFPE | -0.2985 |
| | 1676 | QFIQYTNVQLARPN | 0.583 |
| | 585 | AGRLMQLDTEKNAR | 0.1785 |
| | 1574 | NAFANGNFSQAVME | 0.1903 |
| | 2227 | AQNWTQQYRARVDT | -0.1793 |
| | 2291 | DDEATQWFMTNILN | -0.1257 |
| | 2427 | LNLELSQFRNVLVS | 0.6263 |
| | 113 | NAEKQETVCKMIAD | 0.1319 |
| | 1 | MGNRGSSTSSRPP | 0.9434 |
| | 42 | NGGGDKNPVQ | -0.2157 |
| | 59 | DSVSSKA | 0.7668 |
| | 102 | PDPRAGST | 1.4768 |
| | 112 | KNAEKQET | 0.325 |
| | 142 | DTTDGAAS | 0.996 |
| | 202 | TKAAPNEE | 0.5635 |
| | 251 | DKLQTDE | -0.9348 |
| | 360 | ENSCAKKGGDEEKTP | 0.3963 |
| | 397 | STQAYND | 0.5037 |
| | 453 | NAAAREERET | 0.8255 |

| | | | |
|---|---|---|---|
| | 475 | EEQSKID | 0.6964 |
| | 506 | QKKYGGGEDCECTR | 0.0423 |
| | 533 | SKAARSQ | 0.0266 |
| | 570 | QSYEENY | 0.4129 |
| | 590 | QLDTEKNAR | -0.0868 |
| | 614 | NPGGAQEAD | 0.4438 |
| | 683 | TEATGDT | -0.1499 |
| | 698 | PTSAGNDSNV | 0.6549 |
| | 709 | TDNAPAG | 0.784 |
| | 738 | RASQAKN | -0.5126 |
| | 822 | ARTPEEAAQR | 0.0209 |
| | 960 | GDYSETE | 1.0324 |
| | 984 | EYGEQEATKKA | 0.4039 |
| | 1023 | EARTMND | 0.467 |
| | 1042 | PDEDGYTQSSQ | -0.2356 |
| | 1061 | SPSTQPTPK | 0.6575 |
| Hydrophilicity (Bcepred) | 1133 | KKEHTDK | -0.4423 |
| | 1151 | ESLADTDVNK | 0.0805 |
| | 1206 | RETQAAN | 1.239 |
| | 1251 | PDPTTNATAPQDQN | 0.536 |
| | 1327 | PETSTAQ | 0.0448 |
| | 1371 | YENRSNPGS | 1.0775 |
| | 1393 | KPPTDAGGRP | -0.3661 |
| | 1435 | GGVQRRDAAN | 1.2361 |
| | 1448 | NNNPQPSER | 1.3001 |
| | 1496 | RTNNAQEENT | 0.4246 |
| | 1519 | NVNDAANN | 0.4087 |
| | 1603 | GHRGDPTEQSV | 0.9589 |
| | 1627 | KDTNRQG | -0.0847 |
| | 1650 | KENYRAN | 1.0582 |
| | 1695 | GANNDSV | -0.0391 |
| | 1739 | GDAQNNTSDV | 0.5538 |
| | 1797 | SRYNKEP | 1.2719 |
| | 1821 | ENNEVYD | 0.4586 |
| | 1922 | SENRDDKP | 0.7753 |
| | 1956 | ESSYQEEQ | 0.6747 |
| | 1972 | SPKGQTRT | 0.8322 |
| | 1981 | GSNRERER | 1.155 |
| | 2132 | DYDEAGPG | 0.7224 |
| | 2216 | QNNPTPE | 0.6367 |
| | 2288 | LQTDDEATQ | 0.2673 |
| | 2373 | TEQNREEGP | 1.0089 |
| | 2401 | GKDRFDT | -0.3055 |
| | 2460 | SETFSDKQYDSD | 0.49 |
| | 1 | MGNRGSSTSSR | 1.1239 |
| | 26 | DHIQRQT | -0.3173 |
| | 39 | GYFNGGGDK | -0.7908 |
| | 56 | YHIDSVSSK | 1.1 |
| | 109 | TFVKNAE | 0.1365 |
| | 361 | NSCAKKGGDEEK | 0.1986 |
| | 379 | IEAQRLD | 1.3572 |
| | 392 | EFLNKST | 0.6092 |
| | 453 | NAAAREER | 0.9973 |
| | 530 | LGLSKAA | 0.5932 |
| | 590 | QLDTEKN | -0.5688 |
| | 697 | FPTSAGN | 0.3996 |
| | 724 | EIQQGRSV | 0.7879 |
| | 747 | DLIGRSL | -0.1856 |
| | 1058 | RFISPST | 0.3324 |
| | 1094 | DLVQKFRKQL | -0.1859 |
| | 1130 | NELKKEHT | 0.2077 |
| Flexibility (Bcepred) | 1312 | FEMTSSF | 0.99 |
| | 1369 | QYYENRSNP | 0.582 |
| | 1431 | LQLRGGVQ | 0.8986 |
| | 1446 | QINNNPQPS | 1.1231 |
| | 1472 | ALENNSG | -0.0517 |
| | 1602 | FGHRGDPTE | 1.2233 |
| | 1624 | RLIKDTNR | -0.393 |
| | 1739 | GDAQNNT | 0.4761 |
| | 1794 | NYMSRYN | 0.1863 |
| | 1829 | LYPNLESGS | 0.8892 |
| | 1920 | NLISENRDD | 0.6665 |
| | 1969 | KIVSPKGQTRTLGSNRER | 0.3306 |
| | 2085 | ELMSKGS | 0.7621 |
| | 2142 | AGIQRGR | 1.448 |
| | 2200 | VSMRREV | 0.8442 |
| | 2336 | RWLGPST | 0.3195 |
| | 2370 | TYITEQNREE | 1.0008 |
| | 2454 | FSITGPS | 0.7248 |
| | 3 | NRGSSTSSRPPLSSE | 0.6628 |
| | 21 | YAKLQDHIQRQTRPFSG | 0.1519 |
| | 42 | NGGGDKNPVQHIKDYHIDS | 0.3433 |
| | 62 | SSKAKLRV | 0.6274 |
| | 82 | FKVDTKQPIEDILKDIKKQLPDPRAGSTFVKNAEKQETVCK | 0.1565 |
| | 137 | QDKLIDT | 0.3694 |
| | 178 | NTLENIK | 0.2193 |
| | 190 | IKQLHERMVTE | -0.1368 |
| | 202 | TKAAPNEE | 0.5635 |
| | 221 | YRRLLNEQNLQIN | 0.1067 |
| | 243 | LTPTQKELDKLQTDEV | 0.3683 |
| | 262 | KLLNDTN | -0.1856 |
| | 272 | GTKNFGK | -0.7891 |
| | 294 | ANKINKALQKV | -0.3321 |
| | 307 | KVEQYLQSKNWAEFDKELDLKR | 0.4882 |
| | 348 | LRQTFNERHKILENSCAKKGGDEEKTPLDRRIEAQRLDRKH | 0.3119 |
| | 392 | EFLNKSTQAYNDFLEN | 0.0717 |
| | 423 | PNITRLRD | 0.4768 |
| | 453 | NAAAREERETFLTQ | 0.5147 |
| | 471 | KNVLEEQSKIDPNFKNLYDS | 0.8343 |
| | 502 | TDIVQKKYGGGED | 0.319 |
| | 533 | SKAARSQVDLN | 0.3581 |
| | 559 | YSNLTHNKQEFQSYEENYATI | 0.6158 |
| | 589 | MQLDTEKNARINSP | -0.1172 |

| | | | |
|---|---|---|---|
| | 618 | AQEADWKA | 1.4251 |
| | 631 | ELEYDVKRRFYRALE | -0.0886 |
| | 650 | YLKNITKTF | -0.1566 |
| | 714 | AGHYYEKVAAEIQQGRS | 0.6235 |
| | 736 | RPVRASQAKNIRD | 0.39 |
| | 774 | GGEELRQM | 0.8976 |
| | 790 | YKTLLEY | -0.5026 |
| | 806 | LKNLNQSE | 0.9333 |
| | 821 | LARTPEEAAQRV | 0.0433 |
| | 840 | NDALSTRWETEDV | 0.8817 |
| | 872 | YDMFERPEPVYKLIPTR | -0.082 |
| | 894 | ADELEPEV | 0.4658 |
| | 911 | RLPRLAE | -0.0497 |
| | 923 | FSFRDENVQI | 2.0163 |
| | 960 | GDYSETEIRQL | 0.777 |
| | 982 | NLEYGEQEATKKAL | 0.8327 |
| | 999 | VNEINRRFG | 0.5254 |
| | 1009 | ITRTEWEKFQRI | 0.0801 |
| | 1022 | QEARTMND | 0.3491 |
| | 1032 | MMNQTNYSILPDEDGYTQSSQL | 0.3601 |
| | 1061 | SPSTQPTPKWRPALY | 0.6919 |
| | 1087 | LQPNSQWDLVQKFRKQLSEM | 0.2426 |
| | 1108 | EDPSLQQE | 0.6655 |
| | 1117 | GKVSYQELIRQAINELKKEHTDKIQ | 0.0129 |
| | 1154 | ADTDVNK | -0.5499 |
| | 1182 | LNNFRNNIKG | 0.0736 |
| | 1193 | DLDTIQK | -0.1589 |
| | 1203 | EWLRETQAAN | 0.405 |
| | 1230 | SEIRNPG | -0.4372 |
| Accessibility (Bcepred) | 1239 | VKENDVRLSRVYPDPTTNATAPQDQNLV | 0.5977 |
| | 1285 | GVRAEQELAARYLVDNQRIMQ | 0.2355 |
| | 1323 | QVRFPETSTAQ | 0.4602 |
| | 1351 | DTKYFLN | -0.3542 |
| | 1359 | LRPHIDKNIIQYYENRSNPGS | -0.3289 |
| | 1384 | EEHLIDKLIKPPTDAG | -0.4278 |
| | 1414 | NQIINKT | -0.2704 |
| | 1423 | LLTKPYNVLQ | -0.0093 |
| | 1434 | RGGVQRRDAANIQINNNPQPSERFEQYGR | 0.8653 |
| | 1470 | YDALENNSGLR | -0.139 |
| | 1493 | NLIRTNNAQEENTLSY | 0.2465 |
| | 1510 | DNMAPRTYANVN | 0.5463 |
| | 1523 | AANNLRRYRLYGSDYGIQNNRSMM | 0.3251 |
| | 1559 | RFYDAPSGK | 0.781 |
| | 1592 | HPDLARDN | -0.2998 |
| | 1603 | GHRGDPTEQSV | 0.9589 |
| | 1623 | QRLIKDTNRQGLSQH | -0.3739 |
| | 1646 | PIYLKENYRANLP | 0.8447 |
| | 1675 | KQFIQYT | -0.2976 |
| | 1703 | YYNNNIN | 0.4738 |
| | 1739 | GDAQNNTSDVVRKR | 0.5813 |
| | 1776 | HELTDHP | 0.2001 |
| | 1784 | YLETEEH | 0.299 |
| | 1792 | IQNYMSRYNKEPLMP | 0.4037 |
| | 1812 | YYLRDLRIENNEVYDP | 0.8018 |
| | 1847 | RKLLGND | -1.3827 |
| | 1866 | IMKNYNETV | 0.0046 |
| | 1876 | AREQITPTRFEH | 1.1304 |
| | 1901 | NIRSFKT | -0.3381 |
| | 1909 | MMYNENTFG | 0.2707 |
| | 1920 | NLISENRDDKPII | 0.4988 |
| | 1941 | VYSLRKTLQDV | 0.5651 |
| | 1955 | VESSYQEEQINH | 0.807 |
| | 1972 | SPKGQTRTLGSNRERERIFN | 0.4151 |
| | 2013 | NIYNYDYSFEE | 0.892 |
| | 2032 | SAEKVRSLNTAAPQPDIAE | 0.3342 |
| | 2052 | LNIPNRPPMNTREF | 0.5117 |
| | 2080 | TQYGNEL | 0.5016 |
| | 2105 | LNMGRPKFLSDQIFNK | 0.0403 |
| | 2127 | YPTQFDYDEAGP | 0.8035 |
| | 2143 | GIQRGREQWGQP | 0.3402 |
| | 2168 | RTIRIPQKLRVLRN | -0.2388 |
| | 2201 | SMRREVENM | 0.0645 |
| | 2211 | QTPEIQNNPTPEV | 0.557 |
| | 2227 | AQNWTQQYRARVDT | -0.1793 |
| | 2280 | PVPHPRQILQTDDEATQW | 0.2818 |
| | 2324 | LETLERYV | -0.4118 |
| | 2351 | RMAPRDMRHPISYTEN | 0.9003 |
| | 2370 | TYITEQNREEGPWS | 1.1354 |
| | 2393 | QKPTLVQIGKDRFDTRLIRN | 0.091 |
| | 2420 | QRLLRLR | -1.0025 |
| | 2430 | ELSQFRN | -0.2232 |
| | 2446 | NPSITEY | 0.9181 |
| | 2459 | PSETFSDKQYDSDIR | 0.4142 |
| | 263 | LLNDTNSVL | 0.2647 |
| | 561 | NLTHNKQ | 0.8996 |
| | 700 | SAGNDSNVFT | -0.3106 |
| | 1181 | LLNNFRNN | -0.1998 |
| Turns (Bcepred) | 1445 | IQINNNPQPS | 1.276 |
| | 1473 | LENNSGL | 0.1198 |
| | 1694 | LGANNDSVIYYNNNINVP | 0.215 |
| | 1740 | DAQNNTSDV | 0.5666 |
| | 2214 | EIQNNPTPE | 0.8057 |
| | 28 | IQRQTRP | 1.5005 |
| | 83 | KVDTKQPIEDILKDIKKQLPDPR | -0.1153 |
| | 111 | VKNAEKQET | 0.6654 |
| | 245 | PTQKELDKLQTDE | -0.1665 |
| | 309 | EQYLQSK | -0.111 |
| | 321 | DKELDLKR | 0.5464 |
| | 349 | RQTFNER | 0.2061 |
| | 365 | KKGGDEEKTPLDRRIEAQRLDRKH | 0.4514 |
| | 456 | AREERETF | 0.6645 |
| | 475 | EEQSKIDPNFK | 1.7792 |
| | 503 | DIVQKKY | 1.1319 |

| | | | |
|---|---|---|---|
| | 564 | HNKQEFQS | 0.7439 |
| | 590 | QLDTEKNAR | -0.0868 |
| | 633 | EYDVKRRFYRA | -0.4819 |
| | 807 | KNLNQSE | 0.801 |
| | 877 | RPEPVYK | 0.9711 |
| | 987 | EQEATKKA | 0.0128 |
| | 1010 | TRTEWEKFQR | -0.1942 |
| Exposed Surface (Bcepred) | 1065 | QPTPKWRP | 1.341 |
| | 1096 | VQKFRKQL | -0.7023 |
| | 1129 | INELKKEHTDKI | -0.1687 |
| | 1240 | KENDVRL | -0.0466 |
| | 1360 | RPHIDKN | -2.5275 |
| | 1369 | QYYENRSNP | 0.582 |
| | 1389 | DKLIKPP | -0.918 |
| | 1450 | NPQPSER | 1.8396 |
| | 1525 | NNLRRYR | -1.1041 |
| | 1626 | IKDTNRQG | 0.9863 |
| | 1648 | YLKENYRAN | 0.8444 |
| | 1796 | MSRYNKEPL | 0.8388 |
| | 1923 | SENRDDKPI | 0.8355 |
| | 1982 | SNRERER | 0.8826 |
| | 2171 | RIPQKLR | 0.3714 |
| | 2373 | TEQNREEGP | 1.0089 |
| | 2402 | KDRFDTR | 0.7054 |
| | 2464 | SDKQYDSD | 0.5731 |
| | 25 | QDHIQRQTR | -0.0278 |
| | 51 | QHIKDYHIDS | 0.888 |
| | 62 | SSKAKLRV | 0.6274 |
| | 90 | IEDILKDIKKQL | -0.3973 |
| | 110 | FVKNAEKQETVCK | 0.32 |
| | 162 | THGLRAEY | 0.898 |
| | 189 | AIKQLHERMVTEV | 0.0818 |
| | 203 | KAAPNEE | 0.3283 |
| | 217 | IEAVYRRLLNEQ | -0.2468 |
| | 246 | TQKELDKLQTDEVDIIK | -0.1421 |
| | 317 | WAEFDKELDLKRF | 0.0159 |
| | 336 | ENIAEFEKAVN | -0.0046 |
| | 349 | RQTFNERHKILENSCAKKGGDEEKTPLDRRIEAQRLDRKHILMEI | 0.3185 |
| | 404 | FLENVKKIG | 0.2677 |
| | 453 | NAAAREERETFLT | 0.5779 |
| | 470 | VKNVLEEQSKID | 0.2591 |
| | 513 | EDCECTR | 0.0911 |
| | 562 | LTHNKQEFQSYEE | 0.6479 |
| | 592 | DTEKNARIN | 0.4242 |
| | 631 | ELEYDVKRRFYRALE | -0.0886 |
| | 714 | AGHYYEKVA | 0.6422 |
| | 773 | LGGEELRQMV | 0.5197 |
| | 821 | LARTPEEAAQRV | 0.0433 |
| | 844 | STRWETEDV | 1.0057 |
| | 873 | DMFERPEPVYK | -0.1348 |
| | 894 | ADELEPEVIPE | 0.6488 |
| | 911 | RLPRLAE | -0.0497 |
| | 962 | YSETEIRQLIKEI | -0.1597 |
| | 983 | LEYGEQEATKKALIH | 0.4413 |
| | 999 | VNEINRRFG | 0.5254 |
| | 1009 | ITRTEWEKFQRIVQEAR | 0.2024 |
| | 1095 | LVQKFRKQLSEM | -0.2671 |
| Polarity (Bcepred) | 1128 | AINELKKEHTDKIQ | 0.0373 |
| | 1201 | IIEWLRETQ | 0.3878 |
| | 1220 | DWLGRKHGAISE | 0.6965 |
| | 1239 | VKENDVRL | 0.2154 |
| | 1285 | GVRAEQELAAR | 0.9368 |
| | 1359 | LRPHIDKN | -1.855 |
| | 1380 | FYWLEEHLIDKL | -0.0262 |
| | 1434 | RGGVQRR | 0.3044 |
| | 1452 | QPSERFEQYGR | 0.5909 |
| | 1525 | NNLRRYRLYG | -0.421 |
| | 1592 | HPDLARD | -0.8366 |
| | 1604 | HRGDPTE | 0.73 |
| | 1648 | YLKENYRAN | 0.8444 |
| | 1746 | SDVVRKRLVAV | 0.3918 |
| | 1771 | AMEVLHELTDHPIYLETEEHFIQN | 0.2256 |
| | 1796 | MSRYNKEPL | 0.8388 |
| | 1814 | LRDLRIENNEV | 1.0813 |
| | 1872 | ETVVARE | 0.3788 |
| | 1881 | TPTRFEHFYTHA | 0.3188 |
| | 1922 | ISENRDDKPII | 0.5101 |
| | 1956 | ESSYQEEQINHIHKIVSPK | 0.324 |
| | 1980 | LGSNRERERIFNL | 0.9255 |
| | 2031 | ISAEKVRSLN | 1.0119 |
| | 2062 | TREFMLKL | -0.4779 |
| | 2143 | GIQRGREQWG | 0.7861 |
| | 2162 | ALHELVRTIRIPQKLRVLRN | -0.2801 |
| | 2199 | LVSMRREVENMI | 0.4492 |
| | 2323 | FLETLERYV | -0.5682 |
| | 2353 | APRDMRHPI | 0.3582 |
| | 2372 | ITEQNREEGPWS | 1.4367 |
| | 2400 | IGKDRFDTRL | 0.2474 |
| | 2420 | QRLLRLRLNLE | -0.1004 |
| | 2465 | DKQYDSD | 0.5657 |
| | 50 | VQHIKDYHIDSVS | 1.0775 |
| | 149 | SICRQIVLYINSL | -0.2243 |
| | 169 | YLDVHGSI | -0.122 |
| | 258 | VDIIKLL | -0.493 |
| | 276 | FGKVLSYTLCNLG | 0.4053 |
| | 301 | LQKVGLKVEQYLQS | 0.6113 |
| | 410 | KIGIKLVKEI | 1.1069 |
| | 443 | LDLSLIGFY | 1.3831 |
| | 464 | LTQFMLVKNVLE | -0.3451 |
| | 489 | DSCSRLLQIIDFY | 0.1186 |
| | 514 | DCECTRVG | 1.9253 |
| | 549 | FMYYYYV | 0.532 |
| | 731 | VGTLRPV | 0.5076 |

| | | | |
|---|---|---|---|
| | 782 | VPMSPLQIYKTLLEYIQH | 0.1022 |
| | 830 | QRVYLSTVRV | 0.0137 |
| | 852 | VFFTFML | 0.5489 |
| | 864 | KIFIVLGIYD | 0.8184 |
| | 878 | PEPVYKLIPT | 0.1297 |
| | 897 | LEPEVIP | 1.4756 |
| | 930 | VQISMLP | 1.861 |
| | 974 | INVIYQHFN | 0.7172 |
| | 995 | LIHFVNE | 0.56 |
| | 1049 | QSSQLLPS | 0.3615 |
| | 1077 | IDSVDVQ | 1.7812 |
| | 1116 | LGKVSYQELI | 0.9111 |
| | 1140 | IQIVSKLIQG | 0.2006 |
| | 1158 | VNKIFLFHETVI | -0.0735 |
| | 1178 | IYVLLNNF | -0.5258 |
| | 1233 | RNPGLVVKE | 1.7014 |
| | 1243 | DVRLSRVYPD | 0.7288 |
| | 1265 | LVTETLF | 0.0021 |
| | 1273 | WIVPYVGIP | -0.1519 |
| Antigenic Propensity (Bcepred) | 1302 | RIMQLLLTNIF | -0.3127 |
| | 1321 | MVQVRFPE | 0.4886 |
| | 1333 | QVHLDFT | 2.4399 |
| | 1341 | LISLIDSL | 0.1449 |
| | 1353 | KYFLNLLR | -0.686 |
| | 1386 | HLIDKLI | -0.6454 |
| | 1421 | YILLTKPYNVLQLR | 0.6449 |
| | 1462 | RVFSRLVFYD | 0.0426 |
| | 1478 | GLRVEQVVLGDFR | 1.2714 |
| | 1547 | MVFNQLV | -0.3141 |
| | 1567 | KIYLNLI | -0.3397 |
| | 1609 | TEQSVLLLSLGLMLQRLI | 0.5795 |
| | 1633 | GLSQHLISTL | -0.1724 |
| | 1673 | LLKQFIQY | -1.2539 |
| | 1748 | VVRKRLV | 0.2329 |
| | 1774 | VLHELTD | -0.0936 |
| | 1806 | PFSLSLYYLRDL | 1.209 |
| | 1823 | EVYDPLLYPNL | 0.1967 |
| | 1896 | LRFIVNI | -0.2918 |
| | 1948 | LQDVISFVES | 0.1541 |
| | 1965 | NHIHKIVSP | -0.4582 |
| | 2026 | CLMYGIS | 1.3681 |
| | 2065 | FMLKLLINPYVSVSITQY | 0.5124 |
| | 2117 | IFNKVLFGSLYP | -0.0439 |
| | 2163 | LHELVRT | -0.5247 |
| | 2172 | IPQKLRVLRNIIVKNQLI | -0.538 |
| | 2260 | TITPVTVR | 1.3547 |
| | 2277 | HGIPVPHPRQI | 0.65 |
| | 2329 | RYVFNVPR | -0.7138 |
| | 2366 | NSVLTYITE | -0.1063 |
| | 2382 | WSIVKQVGVGIQKPTLVQIG | 0.8664 |
| | 2419 | IQRLLRLRL | -0.6359 |
| | 2433 | QFRNVLVSPDHII | 0.4361 |
| | 2225 | AAAQNWTQQYRARVDT | -0.0519 |
| | 173 | HGSIENTLENIKLLND | 0.2901 |
| | 1816 | DLRIENNEVYDPLLYP | 0.6771 |
| | 586 | GRLMQLDTEKNARINS | 0.1655 |
| | 34 | PFSGGGYFNGGGDKNP | -0.0141 |
| | 1593 | PDLARDNIAFGHRGDP | 1.0884 |
| | 1508 | YWDNMAPRTYANVNDA | 0.5863 |
| | 572 | YEENYATILGDAIAGR | 0.3358 |
| | 2348 | APVRMAPRDMRHPISY | 0.9017 |
| | 1876 | AREQITPTRFEHFYTH | 0.6396 |
| | 1720 | AALRGIGGVFRPNVTL | 0.1901 |
| | 2342 | TGRVARAPVRMAPRDM | 0.907 |
| | 2334 | VPRWLGPSTGRVARAP | 0.0052 |
| | 2127 | YPTQFDYDEAGPGLAA | 0.7828 |
| | 1226 | HGAISEIRNPGLVVKE | 0.7977 |
| | 512 | GEDCECTRVGGAALTV | 0.5756 |
| | 395 | NKSTQAYNDFLENVKK | 0.0477 |
| | 368 | GDEEKTPLDRRIEAQR | -0.1217 |
| | 2389 | GVGIQKPTLVQIGKDR | 0.5124 |
| | 1928 | DKPIITAGIGMNAVYS | 0.3928 |
| | 1584 | AVMELGYTHPDLARDN | 0.7543 |
| | 1220 | DWLGRKHGAISEIRNP | 0.5448 |
| | 1198 | QKSIIEWLRETQAANV | 0.3467 |
| | 1062 | PSTQPTPKWRPALYNI | 0.7105 |
| | 71 | EGIIRAIAKIGFKVDT | 0.0826 |
| | 2305 | IPAIIMTPFTDLANDL | 0.1007 |
| | 2167 | VRTIRIPQKLRVLRNI | -0.1596 |
| | 101 | LPDPRAGSTFVKNAEK | 0.5703 |
| | 87 | KQPIEDILKDIKKQLP | -0.501 |
| | 674 | GVRIIGRWFTEATGDT | -0.5241 |
| | 285 | CNLGIAASVANKINKA | 0.5823 |
| | 2396 | TLVQIGKDRFDTRLIR | 0.1259 |
| | 2197 | EQLVSMRREVENMIQT | 0.1636 |
| | 2114 | SDQIFNKVLFGSLYPT | 0.0899 |
| | 200 | EVTKAAPNEEVINAVT | 0.1814 |
| | 251 | DKLQTDEVDIIKLLND | -0.3966 |
| | 2212 | TPEIQNNPTPEVIAAA | 0.6342 |
| | 1675 | KQFIQYTNVQLARPNL | 0.2992 |
| | 1642 | LTEIPIYLKENYRANL | 0.8538 |
| | 1623 | QRLIKDTNRQGLSQHL | -0.4305 |
| | 1329 | TSTAQVHLDFTGLISL | 1.1067 |
| | 114 | AEKQETVCKMIADAIN | 0.2359 |
| | 810 | NQSEIGGQRVALARTP | 0.1627 |
| | 696 | SFPTSAGNDSNVFTDN | -0.134 |
| | 2265 | TVRAQLGVIFNRHGIP | 0.8161 |
| | 1890 | THAIQALRFIVNIRSF | 0.2221 |
| | 1244 | VRLSRVYPDPTTNATA | 0.6163 |
| | 1090 | NSQWDLVQKFRKQLSE | 0.1456 |
| | 376 | DRRIEAQRLDRKHILM | 0.4386 |
| | 2453 | GFSITGPSETFSDKQY | 0.4341 |
| | 230 | LQINILTNFIDNILTF | -0.3456 |

| | | | |
|---|---|---|---|
| | 188 | DAIKQLHERMVTEVTK | -0.0768 |
| | 1780 | DHPIYLETEEHFIQNY | 0.1239 |
| | 1602 | FGHRGDPTEQSVLLLS | 0.9858 |
| | 1205 | LRETQAANVNRANLID | 0.9688 |
| | 1006 | FGVITRTEWEKFQRIV | 0.4799 |
| | 476 | EQSKIDPNFKNLYDSC | 1.0795 |
| | 408 | VKKIGIKLVKEIALTF | 0.8344 |
| | 2290 | TDDEATQWFMTNILNI | 0.1234 |
| | 2146 | RGREQWGQPLSEYINQ | -0.1356 |
| | 2029 | YGISAEKVRSLNTAAP | 0.8117 |
| | 1389 | DKLIKPPTDAGGRPLP | -0.5919 |
| | 1364 | DKNIIQYYENRSNPGS | 0.149 |
| | 1022 | QEARTMNDFGMMNQTN | 0.9356 |
| | 819 | VALARTPEEAAQRVYL | 0.222 |
| | 540 | VDLNQAINTFMYYYYV | 0.0457 |
| | 2277 | HGIPVPHPRQILQTDD | 0.2944 |
| | 2056 | NRPPMNTREFMLKLLI | 0.032 |
| | 1984 | RERERIFNLFDMNIIP | 1.0773 |
| | 1905 | FKTVMMYNENTFGGVN | 0.6017 |
| | 1867 | MKNYNETVVAREQITP | 0.8145 |
| | 1765 | HTLADSAMEVLHELTD | 0.1429 |
| | 359 | LENSCAKKGGDEEKTP | 0.4408 |
| | 2245 | IGNIGQPNSMLDLIQT | 0.4114 |
| | 2204 | REVENMIQTPEIQNNP | 0.2896 |
| | 2037 | RSLNTAAPQPDIAEVL | 0.2069 |
| | 138 | DKLIDTTDGAASICRQ | 0.3492 |
| | 1267 | TETLFAWIVPYVGIPA | 0.1872 |
| | 826 | EEAAQRVYLSTVRVND | 0.2556 |
| | 771 | DMLGGEELRQMVPMSP | 0.7385 |
| | 2432 | SQFRNVLVSPDHIINP | 0.2434 |
| | 2283 | HPRQILQTDDEATQWF | 0.0268 |
| | 2255 | LDLIQTITPVTVRAQL | 0.7586 |
| | 1934 | AGIGMNAVYSLRKTLQ | 0.9579 |
| | 1855 | VQLSDMPGVQLIMKNY | 0.6915 |
| | 1691 | MGLLGANNDSVIYYNN | 0.6545 |
| | 164 | GLRAEYLDVHGSIENT | 1.0075 |
| | 1236 | GLVVKENDVRLSRVYP | 0.8823 |
| | 951 | MRPIELINIGDYSETE | 1.5232 |
| | 868 | VLGIYDMFERPEPVYK | 0.2146 |
| | 501 | YTDIVQKKYGGGEDCE | 0.2339 |
| | 311 | YLQSKNWAEFDKELDL | 0.9137 |
| | 214 | VTMIEAVYRRLLNEQN | 0.0716 |
| | 2136 | AGPGLAAGIQRGREQW | 0.7803 |
| | 1730 | RPNVTLMPLGDAQNNT | 0.7928 |
| | 1048 | TQSSQLLPSDRFISPS | 0.3611 |
| | 728 | GRSVGTLRPVRASQAK | 0.4482 |
| | 667 | TVQQMLDGVRIIGRWF | -0.5403 |
| | 641 | YRALEGLDLYLKNITK | 0.5554 |
| | 629 | AIELEYDVKRRFYRAL | 0.0834 |
| | 2186 | NQLIADLTTIREQLVS | 0.2033 |
| | 2078 | SITQYGNELMSKGSAG | 0.3724 |
| | 207 | NEEVINAVTMIEAVYR | 0.2953 |
| | 1835 | SGSPEFKLLYGTRKLL | 0.4916 |
| | 1822 | NEVYDPLLYPNLESGS | 0.4517 |
| | 1460 | YGRVFSRLVFYDALEN | -0.0763 |
| | 1395 | PTDAGGRPLPGGELGL | 0.4368 |
| | 1301 | QRIMQLLLTNIFEMTS | 0.0069 |
| | 1277 | YVGIPAGGGVRAEQEL | 0.7806 |
| | 12 | PPLSSEANLYAKLQDH | 0.6633 |
| | 989 | EATKKALIHFVNEINR | 0.1437 |
| | 615 | PGGAQEADWKAAVSAI | 0.7178 |
| | 608 | RGHVGPNPGGAQEADW | 1.1441 |
| | 564 | HNKQEFQSYEENYATI | 0.5863 |
| | 548 | TFMYYYYVAQIYSNLT | 0.3242 |
| | 2446 | NPSITEYGFSITGPSE | 0.8065 |
| | 2377 | REEGPWSIVKQVGVGI | 1.0867 |
| | 2323 | FLETLERYVFNVPRWL | -0.797 |
| | 1789 | EHFIQNYMSRYNKEPL | -0.1606 |
| | 1552 | LVASYIARFYDAPSGK | 0.2629 |
| | 1435 | GGVQRRDAANIQINNN | 1.121 |
| | 1323 | QVRFPETSTAQVHLDF | 0.8771 |
| | 1084 | TGMLQPNSQWDLVQKF | 0.3438 |
| ABCpred | 1040 | ILPDEDGYTQSSQLLP | -0.0241 |
| | 899 | PEVIPEAAELYFRLPR | 0.078 |
| | 888 | RMILGGADELEPEVIP | 0.6247 |
| | 43 | GGGDKNPVQHIKDYHI | 0.6343 |
| | 2012 | ANIYNYDYSFEEIACL | 1.0988 |
| | 1636 | QHLISTLTEIPIYLKE | 0.446 |
| | 1312 | FEMTSSFNKMVQVRFP | 0.1625 |
| | 95 | KDIKKQLPDPRAGSTF | 0.3562 |
| | 880 | PVYKLIPTRMILGGAD | -0.0093 |
| | 794 | LEYIQHSALSVGLKNL | 1.2513 |
| | 780 | QMVPMSPLQIYKTLLE | 0.331 |
| | 765 | AFARIGDMLGGEELRQ | 0.6836 |
| | 714 | AGHYYEKVAAEIQQGR | 0.6563 |
| | 683 | TEATGDTLAQVFESFP | -0.1431 |
| | 422 | TPNITRLRDALSRIND | -0.1618 |
| | 1536 | DYGIQNNRSMMMVFNQ | 0.8644 |
| | 1483 | QVVLGDFRLSNLIRTN | 0.9942 |
| | 1145 | KLIQGSESLADTDVNK | -0.0219 |
| | 722 | AAEIQQGRSVGTLRPV | 0.7366 |
| | 524 | ALTVEELGLSKAARSQ | 0.6424 |
| | 518 | TRVGGAALTVEELGLS | 1.0307 |
| | 346 | NLLRQTFNERHKILEN | -0.0451 |
| | 25 | QDHIQRQTRPFSGGGY | 0.2816 |
| | 2425 | LRLNLELSQFRNVLVS | 0.5167 |
| | 236 | TNFIDNILTPTQKELD | 0.453 |
| | 2089 | KGSAGYMSRIFRGDNA | 0.044 |
| | 2044 | PQPDIAEVLNIPNRPP | 0.4052 |
| | 2006 | MRSIPLANIYNYDYSF | 1.1486 |
| | 1919 | VNLISENRDDKPIITA | 0.6231 |
| | 1806 | PFSLSLYYLRDLRIEN | 1.3904 |
| | 1528 | RRYRLYGSDYGIQNNR | 0.6435 |

| | | |
|---|---|---|
| 1 | MGNRGSSTSSRPPLSS | 0.7636 |
| 734 | LRPVRASQAKNIRDLI | 0.4326 |
| 580 | LGDAIAGRLMQLDTEK | 0.1658 |
| 554 | YVAQIYSNLTHNKQEF | 0.4037 |
| 51 | QHIKDYHIDSVSSKAK | 1.1527 |
| 2369 | LTYITEQNREEGPWSI | 1.2794 |
| 1651 | ENYRANLPLFNKMFNI | 0.2508 |
| 130 | QEFIDLGQDKLIDTTD | 0.719 |
| 1252 | DPTTNATAPQDQNLVT | 0.4219 |
| 836 | TVRVNDALSTRWETED | 0.9187 |
| 432 | LSRINDMGTIALDLSL | 0.9818 |
| 332 | LVSAENIAEFEKAVNL | 0.4903 |
| 2105 | LNMGRPKFLSDQIFNK | 0.0403 |
| 1999 | PINVNALMRSIPLANI | 0.6151 |
| 1977 | TRTLGSNRERERIFNL | 0.508 |
| 1754 | VAVIDGIIRGSHTLAD | 0.1019 |
| 1405 | GGELGLEGVNQIINKT | 0.7027 |
| 845 | TRWETEDVFFTFMLKS | 0.9762 |
| 533 | SKAARSQVDLNQAINT | 0.0457 |
| 1958 | SYQEEQINHIHKIVSP | 0.3889 |
| 1945 | RKTLQDVISFVESSYQ | 0.4559 |
| 1466 | RLVFYDALENNSGLRV | 0.0493 |
| 1130 | NELKKEHTDKIQIVSK | 0.2641 |
| 1028 | NDFGMMNQTNYSILPD | 1.0107 |
| 997 | HFVNEINRRFGVITRT | 0.7762 |
| 649 | LYLKNITKTFVNNIDS | -0.0908 |
| 495 | LQIIDFYTDIVQKKYG | 0.5194 |
| 438 | MGTIALDLSLIGFYTN | 0.8414 |
| 968 | RQLIKEINVIYQHFNL | 0.3556 |
| 961 | DYSETEIRQLIKEINV | 0.3142 |
| 917 | EFYQKLFSFRDENVQI | 0.6313 |
| 747 | DLIGRSLSNFQALKNI | 0.1119 |
| 1683 | VQLARPNLMGLLGANN | 0.8125 |
| 1558 | ARFYDAPSGKIYLNLI | 0.219 |
| 154 | IVLYINSLTHGLRAEY | 0.4875 |
| 1350 | ADTKYFLNLLRPHIDK | -0.8505 |
| 122 | KMIADAINQEFIDLGQ | 0.6557 |
| 1174 | LLSAIYVLLNNFRNNI | -0.1204 |
| 2236 | ARVDTLINFIGNIGQP | -0.0874 |
| 1629 | TNRQGLSQHLISTLTE | -0.1138 |
| 1285 | GVRAEQELAARYLVDN | 0.2493 |
| 1074 | LYNIDSVDVQTGMLQP | 1.0224 |
| 456 | AREERETFLTQFMLVK | 0.2262 |
| 2357 | MRHPISYTENSVLTYI | 0.2089 |
| 2312 | PFTDLANDLRTFLETL | -0.4169 |
| 2179 | LRNIIVKNQLIADLTI | -0.0951 |
| 1337 | DFTGLISLIDSLMADT | 0.2057 |
| 1123 | ELIRQAINELKKEHTD | -0.085 |
| 2098 | IFRGDNALNMGRPKFL | 0.2734 |
| 596 | NARINSPAVDLARGHV | 0.2935 |
| 467 | FMLVKNVLEEQSKIDP | 0.5422 |
| 222 | RRLLNEQNLQINILTN | -0.0565 |
| 319 | EFDKELDLKRFSGLVS | -0.0137 |
| 2459 | PSETFSDKQYDSDIRI | 0.4595 |
| 2019 | YSFEEIACLMYGISAE | 1.124 |
| 1668 | ISQGELLKQFIQYTNV | 0.0009 |
| 1564 | PSGKIYLNLINAFANG | -0.3041 |
| 1295 | RYLVDNQRIMQLLLTN | -0.0942 |
| 704 | DSNVFTDNAPAGHYYE | -0.2865 |
| 1740 | DAQNNTSDVVRKRLVA | 0.2874 |
| 145 | DGAASICRQIVLYINS | -0.1405 |
| 929 | NVQISMLPELEGIFSG | 0.8341 |
| 267 | TNSVLGTKNFGKVLSY | 0.5281 |
| 1377 | PGSFYWLEEHLIDKLI | -0.1654 |
| 787 | LQIYKTLLEYIQHSAL | -0.112 |
| 2062 | TREFMLKLLINPYVSV | -0.0941 |
| 1699 | DSVIYYNNNINVPMTG | 0.2091 |
| 1570 | LNLINAFANGNFSQAV | 0.2914 |
| 1116 | LGKVSYQELIRQAINE | 0.0094 |
| 938 | LEGIFSGLIRIIFMRP | -0.4577 |
| 258 | VDIIKLLNDTNSVLGT | 0.2659 |
| 18 | ANLYAKLQDHIQRQTR | 0.3803 |
| 1415 | QIINKTYILLTKPYNV | 0.4989 |
| 1185 | FRNNIKGLDLDTIQKS | 0.5423 |
| 305 | GLKVEQYLQSKNWAEF | 0.7527 |
| 2156 | SEYINQALHELVRTIR | -0.1915 |
| 1104 | SEMFEDPSLQQELGKV | -0.0198 |
| 444 | DLSLIGFYTNAAAREE | 0.7365 |
| 299 | KALQKVGLKVEQYLQS | 0.492 |
| 2411 | RNLIFITNIQRLLRLR | 0.3842 |
| 1514 | PRTYANVNDAANNLRR | 0.1611 |
| 1444 | NIQINNNPQPSERFEQ | 0.8755 |
| 382 | QRLDRKHILMEFLNKS | 0.5673 |
| 1796 | MSRYNKEPLMPFSLSL | 1.0742 |
| 2297 | WFMTNILNIPAIIMTP | 0.1706 |
| 975 | NVIYQHFNLEYGEQEA | 1.0589 |
| 905 | AAELYFRLPRLAEFYQ | 0.1467 |
| 944 | GLIRIIFMRPIELINI | 0.2636 |
| 1708 | INVPMTGLSVGQAALR | 0.8003 |
| 1166 | ETVITGLNLLSAIYVL | 0.4868 |
| 485 | KNLYDSCSRLLQIIDF | 0.3052 |
| 277 | GKVLSYTLCNLGIAAS | 0.6523 |
| 1096 | VQKFRKQLSEMFEDPS | -0.5258 |
| 4 | RGSSTSSRPPLSSEAN | 0.5928 |
| 31 | QTRPFSGGGYFNGGGDKNPVQH | -0.0273 |
| 56 | YHIDSVSS | 0.8076 |
| 99 | KQLPDPRAG | 0.7043 |
| 139 | KLIDTTDGAAS | 0.7876 |
| 171 | DVHGSIE | 0.2737 |
| 265 | NDTNSVLGTKNFGKVL | 0.5112 |
| 312 | LQSKNWAE | 1.1112 |
| 362 | SCAKKGGDEEKTPLD | 0.5626 |
| 395 | NKSTQAYNDFL | 0.1925 |

| | | | |
|---|---|---|---|
| | 433 | SRINDMG | 1.4791 |
| | 478 | SKIDPNFKNLYDSCSR | 0.8279 |
| | 508 | KYGGGEDCECTRVG | 0.5751 |
| | 558 | IYSNLTHNKQEF | 0.4969 |
| | 597 | ARINSPAV | -0.05 |
| | 609 | GHVGPNPGGAQE | 0.9005 |
| | 697 | FPTSAGNDSNVFTDNAPAGHYY | -0.0263 |
| | 727 | QGRSVGT | 0.7949 |
| | 749 | IGRSLSN | -0.081 |
| | 783 | PMSPLQ | 0.8908 |
| | 805 | GLKNLNQSEIGG | 1.1203 |
| | 958 | NIGDYSE | 1.2509 |
| | 1026 | TMNDFGMMNQTNYSILPDEDGYTQSSQLLPSDRFISPSTQPTPKWRPALYNIDSVDV | 0.6081 |
| | 1086 | MLQPNSQWD | 0.434 |
| | 1107 | FEDPSLQQ | 0.4423 |
| | 1147 | IQGSESLADTDV | 0.4128 |
| | 1184 | NFRNNIKGLDL | 1.5646 |
| | 1222 | LGRKHGA | 0.3081 |
| | 1232 | IRNPGL | -0.2176 |
| | 1248 | RVYPDPTTNATAPQDQNL | 0.5918 |
| | 1278 | VGIPAGGGVR | 0.6593 |
| | 1371 | YENRSNPGSFY | 0.7468 |
| | 1392 | IKPPTDAGGRPLPGGELG | 0.1018 |
| | 1432 | QLRGGVQR | -0.8326 |
| | 1446 | QINNNPQPSER | 1.2039 |
| Chou & Fasman Beta-Turn Prediction (IEDB) | 1472 | ALENNSGLR | -0.0841 |
| | 1495 | IRTNNAQE | 0.1606 |
| | 1504 | NTLSYWDNMAP | 0.5861 |
| | 1516 | TYANVNDAANNLRR | 0.2834 |
| | 1531 | RLYGSDYGIQNNRS | 0.3696 |
| | 1561 | YDAPSGKIY | 0.0354 |
| | 1576 | FANGNFS | 1.2901 |
| | 1590 | YTHPDLAR | 0.154 |
| | 1600 | IAFGHRGDPTE | 1.8193 |
| | 1627 | KDTNRQGLS | 0.1356 |
| | 1694 | LGANNDSVIYYNNNINVPMTGLSV | 0.4455 |
| | 1724 | GIGGVFR | 0.8158 |
| | 1737 | PLGDAQNNTSDVV | 0.6625 |
| | 1760 | IIRGSHTL | 0.4989 |
| | 1794 | NYMSRYNKEP | 0.6757 |
| | 1805 | MPFSLSL | 1.6461 |
| | 1827 | PLLYPNLESGSPE | 0.6856 |
| | 1844 | YGTRKLLGNDPVQLSDMPG | 0.0201 |
| | 1912 | NENTFGGVNLISENRDDKP | 0.613 |
| | 1971 | VSPKGQTRTLGSNRE | 0.5916 |
| | 2013 | NIYNYDYS | 0.8879 |
| | 2040 | NTAAPQPD | 0.1028 |
| | 2053 | NIPNRPPMNT | -0.0378 |
| | 2079 | ITQYGNELMSKGSAGYMS | 0.4029 |
| | 2099 | FRGDNALNMGRPKFL | 0.4141 |
| | 2125 | SLYPTQFDYDEAGPGLA | 0.9549 |
| | 2148 | REQWGQPLSEYI | -0.5391 |
| | 2214 | EIQNNPTP | 1.0973 |
| | 2246 | GNIGQPNSM | 0.3214 |
| | 2277 | HGIPVPHPR | 0.7386 |
| | 2314 | TDLANDL | 0.0822 |
| | 2336 | RWLGPSTGR | 0.2918 |
| | 2354 | PRDMRHPISYTENSV | 0.5277 |
| | 2377 | REEGPWSI | 1.3844 |
| | 2401 | GKDRFD | 0.3737 |
| | 2438 | LVSPDHIINPSITEYGFSITGPSETFSDKQYDSDI | 0.4961 |
| | 4 | RGSSTSSRPPLSSEA | 0.5822 |
| | 28 | IQRQTRPFSGGGYFNGGGDKNPV | 0.1788 |
| | 59 | DSVSSKAK | 1.1248 |
| | 83 | KVDTKQPIEDILKDIKKQLPDPRAGSTFVKNAEKQET | 0.1516 |
| | 136 | GQDKLIDTTDG | 0.4522 |
| | 174 | GSIENTLENIKLLND | 0.3846 |
| | 200 | EVTKAAPNEE | 0.3043 |
| | 224 | LLNEQNL | 0.9065 |
| | 243 | LTPTQKELDKLQTDE | 0.3414 |
| | 265 | NDTNSVLGTKNFGK | 0.61 |
| | 296 | KINKALQKV | -0.529 |
| | 307 | KVEQYLQSKN | 0.4165 |
| | 348 | LRQTFNERHK | 0.911 |
| | 359 | LENSCAKKGGDEEKTPLDRRIEAQRLDRK | 0.3042 |
| | 395 | NKSTQA | 0.4551 |
| | 406 | ENVKKIG | 0.0677 |
| | 422 | TPNITRLRDALSRINDMG | 0.0821 |
| | 457 | REERETFL | 0.5965 |
| | 474 | LEEQSKIDPNFKNLYDSCSR | 0.8642 |
| | 505 | VQKKYGGGEDC | -0.0247 |
| | 533 | SKAARSQV | -0.2096 |
| | 564 | HNKQEFQSYEEN | 0.5904 |
| | 592 | DTEKNARINSPA | -0.0079 |
| | 611 | VGPNPGGAQEA | 0.4001 |
| | 652 | KNITKT | -0.0289 |
| | 661 | NIDSIQTVQ | 0.6686 |
| | 683 | TEATGDT | -0.1499 |
| | 696 | SFPTSAGNDSNV | 0.369 |
| | 725 | IQQGRSVGTLRPVRASQAKNIRDLIGRSLS | 0.2763 |
| | 771 | DMLGGEELR | 1.1718 |
| | 807 | KNLNQSEIGGQR | 0.7862 |
| | 823 | RTPEEA | 0.1105 |
| | 841 | DALSTRWETED | 0.923 |
| | 892 | GGADELEPE | 0.8061 |
| | 960 | GDYSETEIRQ | 0.8982 |
| | 985 | YGEQEATKK | 0.0809 |
| | 1010 | TRTEWEKFQ | 0.044 |
| | 1022 | QEARTMN | 0.6736 |
| | 1042 | PDEDGYTQSSQLLPSDR | 0.0551 |
| | 1061 | SPSTQPTPKWRP | 0.9506 |
| | 1077 | IDSVDVQTG | 1.5413 |
| | 1087 | LQPNSQ | 1.0542 |

| | | | |
|---|---|---|---|
| Karplus & Schulz Flexibility Prediction (IEDB) | 1098 | KFRKQLS | -0.3866 |
| | 1108 | EDPSLQQELGKV | 0.1898 |
| | 1131 | ELKKEHTDK | 0.0312 |
| | 1145 | KLIQGSESLADTDVN | 0.2482 |
| | 1185 | FRNNIKGLDLDTIQKS | 0.5423 |
| | 1231 | EIRNPG | -0.1238 |
| | 1239 | VKENDV | 1.1085 |
| | 1250 | YPDPTTNATAPQDQNLVTET | 0.4552 |
| | 1282 | AGGGVRAEQE | 1.2071 |
| | 1327 | PETSTA | 0.1159 |
| | 1360 | RPHIDK | -4.4436 |
| | 1372 | ENRSNPGS | 1.1014 |
| | 1390 | KLIKPPTDAGGRPLPGGEL | -0.3742 |
| | 1434 | RGGVQRRD | 1.0429 |
| | 1447 | INNNPQPSERFEQYG | 0.8342 |
| | 1473 | LENNSGL | 0.1198 |
| | 1496 | RTNNAQEENT | 0.4246 |
| | 1533 | YGSDYGIQNNR | 0.7649 |
| | 1592 | HPDLAR | -0.8175 |
| | 1605 | RGDPTEQS | 1.1089 |
| | 1626 | IKDTNRQGLS | 0.7855 |
| | 1669 | SQGELLK | -0.8381 |
| | 1739 | GDAQNNTSDVVRKR | 0.5813 |
| | 1798 | RYNKEP | 1.7109 |
| | 1818 | RIENNEVYDPL | 0.2982 |
| | 1831 | PNLESGSPEF | 0.9895 |
| | 1845 | GTRKLLGNDPV | -0.7699 |
| | 1877 | REQITPTR | 0.9647 |
| | 1913 | ENTFGG | 0.726 |
| | 1923 | SENRDDKP | 0.7753 |
| | 1945 | RKTLQD | 0.4189 |
| | 1956 | ESSYQEEQI | 0.6544 |
| | 1972 | SPKGQTRTLGSNRERER | 0.6468 |
| | 2033 | AEKVRSLNTAAPQPD | 0.3971 |
| | 2054 | IPNRPPMNTR | 0.5829 |
| | 2080 | TQYGNELMSKGSA | 0.4545 |
| | 2108 | GRPKFLS | 1.1474 |
| | 2133 | YDEAGPG | 0.5829 |
| | 2145 | QRGREQWGQPLS | 0.0651 |
| | 2169 | TIRIPQKL | 0.0551 |
| | 2193 | TTIREQ | 0.8889 |
| | 2212 | TPEIQNNPTPE | 0.6836 |
| | 2229 | NWTQQY | 0.2473 |
| | 2247 | NIGQPNS | -0.0175 |
| | 2289 | QTDDEAT | 0.5255 |
| | 2312 | PFTDLANDLRT | 0.2645 |
| | 2335 | PRWLGPSTGR | 0.5575 |
| | 2373 | TEQNREEGP | 1.0089 |
| | 2401 | GKDRFDTRL | 0.4472 |
| | 2456 | ITGPSETFSDKQYDSD | 0.1476 |
| Parker Hydrophilicity Prediction (IEDB) | 4 | RGSSTSSRPPLSSEANL | 0.6024 |
| | 24 | LQDHIQRQTRPFSG | -0.0975 |
| | 39 | GYFNGGGDKNPVQHIK | -0.034 |
| | 56 | YHIDSVSSKAK | 1.1242 |
| | 84 | VDTKQPI | 0.0984 |
| | 97 | IKKQLPDPRAGSTFVKNAEKQETV | 0.4489 |
| | 139 | KLIDTTDGAASI | 0.8063 |
| | 174 | GSIENTLE | 0.5305 |
| | 197 | MVTEVTKAAPNEEVI | 0.2936 |
| | 246 | TQKELDKLQTDEVD | 0.024 |
| | 265 | NDTNSVLG | 0.9031 |
| | 292 | SVANKINKALQ | 0.0815 |
| | 315 | KNWAEFDK | 0.1821 |
| | 336 | ENIAEFEK | 0.0095 |
| | 351 | TFNERHKILE | 0.8829 |
| | 362 | SCAKKGGDEEKTPLDRRIE | 0.4497 |
| | 395 | NKSTQAY | 0.0467 |
| | 454 | AAAREERETF | 0.8104 |
| | 473 | VLEEQSKIDPNF | 1.3622 |
| | 486 | NLYDSCS | 0.1099 |
| | 504 | IVQKKYGGGEDCECTRVG | 0.7061 |
| | 534 | KAARSQVDL | 0.0953 |
| | 562 | LTHNKQEFQSYEENY | 0.6631 |
| | 592 | DTEKNARINSPAV | 0.1259 |
| | 610 | HVGPNPGGAQEADWKA | 1.0586 |
| | 660 | NNIDSIQT | 0.3858 |
| | 685 | ATGDTL | -0.1734 |
| | 697 | FPTSAGNDSNVFTDNAPAGHYYEKVAAEIQQGRSVGT | 0.2931 |
| | 738 | RASQAKNI | -0.1651 |
| | 808 | NLNQSEIGGQR | 0.724 |
| | 823 | RTPEEAAQ | 0.1476 |
| | 875 | FERPEP | -0.3861 |
| | 893 | GADELE | 0.9109 |
| | 926 | RDENVQ | 0.2601 |
| | 960 | GDYSETEI | 1.0392 |
| | 984 | EYGEQEATK | 0.4015 |
| | 1021 | VQEARTMND | 0.5162 |
| | 1042 | PDEDGYTQSSQLLP | -0.0584 |
| | 1062 | PSTQPTPKW | 0.3148 |
| | 1078 | DSVDVQT | 0.6146 |
| | 1086 | MLQPNSQ | 0.4722 |
| | 1110 | PSLQQELGKVSY | 0.6279 |
| | 1128 | AINELKKEHTDK | 0.0656 |
| | 1147 | IQGSESLADTDV | 0.4128 |
| | 1185 | FRNNIK | 0.1373 |
| | 1208 | TQAANVNR | 0.4931 |
| | 1225 | KHGAISEIRN | 1.0093 |
| | 1239 | VKENDVRL | 0.2154 |
| | 1249 | VYPDPTTNATAPQDQNLV | 0.5762 |
| | 1281 | PAGGGVRAEQELA | 1.0098 |
| | 1326 | FPETSTAQV | 0.2906 |
| | 1371 | YENRSNPG | 1.1771 |
| | 1395 | PTDAGGRPLP | 0.0408 |

| | | |
|---|---|---|
| 1435 | GGVQRRDAANIQINNNPQPSERFEQYG | 1.0399 |
| 1473 | LENNSGLR | -0.237 |
| 1495 | IRTNNAQEENT | 0.3447 |
| 1512 | MAPRTYANVNDAANNLR | 0.5238 |
| 1533 | YGSDYGIQNNRS | 0.7164 |
| 1561 | YDAPSG | 0.7471 |
| 1590 | YTHPDLARDNIA | 0.0413 |
| 1604 | HRGDPTEQ | 0.9122 |
| 1626 | IKDTNRQGL | 0.8 |
| 1695 | GANNDSVI | 0.1014 |
| 1739 | GDAQNNTSDVV | 0.5007 |
| 1765 | HTLADSAM | -0.0332 |
| 1786 | ETEEHF | 0.846 |
| 1795 | YMSRYNK | 0.2827 |
| 1818 | RIENNEVY | 0.1508 |
| 1833 | LESGSPE | 0.5346 |
| 1869 | NYNETVVA | 0.886 |
| 1912 | NENTFG | 0.0982 |
| 1923 | SENRDDKPI | 0.8355 |
| 1956 | ESSYQEEQIN | 0.7039 |
| 1971 | VSPKGQTRTLGSNRERER | 0.8021 |
| 2015 | YNYDYSF | 1.6852 |
| 2031 | ISAEKVRSLNTAAPQPDI | 0.5149 |
| 2078 | SITQYGNELMSKGSAGY | 0.3863 |
| 2099 | FRGDNALN | 0.3331 |
| 2130 | QFDYDEAGPGL | 0.6622 |
| 2144 | IQRGREQWG | 0.6388 |
| 2203 | RREVENMIQ | -0.0986 |
| 2213 | PEIQNNPTP | 1.0029 |
| 2228 | QNWTQQYRARV | 0.123 |
| 2248 | IGQPNS | -0.2883 |
| 2288 | LQTDDEAT | 0.35 |
| 2315 | DLANDL | -0.3517 |
| 2341 | STGRVA | 0.4355 |
| 2353 | APRDMRH | 0.8374 |
| 2361 | ISYTENSV | 1.245 |
| 2373 | TEQNREEGP | 1.0089 |
| 2401 | GKDRFD | 0.3737 |
| 2456 | ITGPSETFSDKQYDSD | 0.1476 |

**Table S7. Eight antigenic and four non-antigenic B cell linear epitopes predicted by VirusImmu for ASFV pp220 protein.**

| Start | End | Peptide | Peptide length | VirusImmunScore | Antigenicity (VirusImmu) | AntigenicityScore (VaxiJen) | Allergenicity (Allergen FP 1.0) | Toxicity (ToxinPred) | Method |
|---|---|---|---|---|---|---|---|---|---|
| 951 | 966 | MRPIELINIGDYSETE | 16 | 0.67432579 | Antigenic | 1.5232 | NON-ALLERGEN | Non-Toxin | ABCpred |
| 957 | 966 | INIGDYSETE | 10 | 0.62304061 | Antigenic | 1.7685 | NON-ALLERGEN | Non-Toxin | Bepipred2.0 (IEDB) |
| 477 | 486 | QSKIDPNFKN | 10 | 0.56450307 | Antigenic | 1.951 | NON-ALLERGEN | Non-Toxin | Bepipred2.0 (IEDB) |
| 618 | 625 | AQEADWKA | 8 | 0.4962204 | Antigenic | 1.4251 | NON-ALLERGEN | Non-Toxin | Accessibility (Bcepred) |
| 1600 | 1610 | IAFGHRGDPTE | 11 | 0.42618034 | Antigenic | 1.8193 | NON-ALLERGEN | Non-Toxin | Chou &Fasman Beta-Turn (IEDB) |
| 1806 | 1815 | PFSLSLYYLRDLRIEN | 10 | 0.4211895 | Antigenic | 1.3904 | NON-ALLERGEN | Non-Toxin | ABCpred |
| 102 | 109 | PDPRAGST | 8 | 0.40746678 | Antigenic | 1.4768 | NON-ALLERGEN | Non-Toxin | Hydrophilicity (Bcepred) |
| 980 | 990 | HFNLEYGEQEA | 11 | 0.40496708 | Antigenic | 1.6057 | NON-ALLERGEN | Non-Toxin | Bepipred2.0 (IEDB) |
| 473 | 484 | VLEEQSKIDPNF | 12 | 0.32176303 | Non-antigenic | 1.3622 | NON-ALLERGEN | Non-Toxin | Parker Hydrophilicity (IEDB) |
| 1233 | 1241 | RNPGLVVKE | 9 | 0.29628091 | Non-antigenic | 1.7014 | NON-ALLERGEN | Non-Toxin | Antigenic Propensity (Becpred) |
| 1065 | 1072 | QPTPKWRP | 8 | 0.28975268 | Non-antigenic | 1.341 | NON-ALLERGEN | Non-Toxin | Exposed Surface (Bcepred) |
| 1448 | 1456 | NNNPQPSER | 9 | 0.21236347 | Non-antigenic | 1.3001 | NON-ALLERGEN | Non-Toxin | Hydrophilicity (Bcepred) |

### S1. Materials and Methods

#### Dataset of training and test

Protein datasets were constructed by 100 antigens (Positive set) (Table S1) and 100 non-antigens (Negative set) (Table S2). An antigen is a protein that can cause an immune response in pertinent immunization test. Protected antigens were the validated protein antigens which were collected from the literatures containing tested immunogenic proteins data, and the corresponding protein sequences were collected from UniProt [1] and NCBI [2]. Protein with complete fragments were preferred. Unprotected protein sequences (non-antigens) were randomly selected from Viral Bioinformatics Resource Center [3]. A BLAST (Expectation value of 3.0) [4] was performed to confirm that non-antigens have no sequence identity to the antigens to ensure that negative sequences were not obviously related to the positive set at the sequence level.

A random sampling cross validation strategy was applied to derive test set from 20% of the positive datasets and negative dataset. Thus, every test set contained 20 antigens and 20 non-antigens and the training set consisted of 80 antigens and 80 non-antigens. 50 randomizations were performed.

#### Dataset of external evaluation

The external dataset was constructed to ensure the universality and accuracy of our models (Table S3). This dataset consisted of 59 antigens and 54 non-antigens after removal of some duplicate proteins. The protective antigens' sequences were manually curated from UniProt and the Protegen database [5] in the same way of training set. The non-antigens' sequences were selected randomly from UniProt, and were also not obviously related to the antigens at the sequence level after a BLAST (Expectation value of 3.0).

#### Descriptors

Z-descriptors, originally proposed by Hellberg [6], have been widely used for the characterization [7] and classification [8] of proteins. They are highly condensed descriptors and are derived from principal component analysis (PCA) of 29 experimental or calculated physicochemical properties of the twenty naturally occurring amino acids [3]. Each amino acid in the protein sequence was represented by three z descriptors: $Z1$ represents hydrophobicity, $Z2$ steric properties and $Z3$ polarity of the amino acid (Table S4).

E-descriptors, proposed by Venkatarajan and Braun [9], were also used to quantitatively characterize the protein sequences. They derived five numerical values for each of the 20 naturally occurring amino acids based on the PCA of 237 physicochemical properties [10]:$E1$represents hydrophobicity, $E2$ molecular size and the steric properties, $E3$ propensity for occurrence in α-helices, $E4$ partial specific volume, number of codons, and relative frequency, and $E5$ propensity for occurrence in β-strands (Table S4).

Each protein in our datasets was represented as a two-dimensional array ($5 \times N$) by using the E-descriptors, where 5 is the number of descriptors and N is the protein length. At the same time, we also used the Z-descriptors ($3 \times N$) to represent protein and compared the difference between the two descriptors in the process of viral immunogenicity prediction.

#### Auto-Cross Covariance (ACC)

As the proteins used in the study had different lengths, an auto-cross covariance (ACC) transformation, which introduced in 1993 [11], was used to transform them to a uniform equal-length vector. Auto covariance $A_{jj}(lag)$ and Cross covariance $C_{jk}(lag)$ were calculated by the following formulas:

$$A_{jj}(L) = \sum_{i}^{n-L} \frac{E_{j,i} \times E_{j,i+L}}{n-L} \tag{1}$$

$$C_{jk}(L) = \sum_{i}^{n-L} \frac{E_{j,i} \times E_{k,i+L}}{n-L} \tag{2}$$

Index $j$ and $k$ refer to the the Z-scales (1, 2, 3) or E-scales (1, 2, 3, 4, 5), $n$ is the number of amino acids in a virus protein sequence, index $i$ indicates the position of amino acid ($i$ = 1, 2, ...n) and $L$ is the lag ($L$= 1, 2, ...$L$). A short range of lag ($L$), which are the lengths of the frame of contiguous amino acids, were used to calculated $A_{jj}$ and $C_{jk}$, to investigate the influence of close amino acid proximity on protein antigenicity. Different descriptors and lags ($L$) produced varying feature numbers. These proteins were presented as numerical matrix of $3 \times N$ Z-descriptors and $5 \times N$ E-descriptors, respectively. $N$ is the number of amino acid residues. Matrices of different lengths were transformed into uniform vectors by ACC-transformation with different lags ($L$). The results of ACC of all protein sequences are also freely accessible at https://github.com/zhangjbig/VirusImmu.

#### Feature selection

Feature selection (FS) is one of the most important steps in the machine learning process for constructing predicting models, as identification and ranking of the most relevant features greatly affect the computational speed and predictive capability [12-14]. In this study, we chose Random Forest (RF) [15] as an embedded method to obtain feature importance. Each protein was transformed into a uniform vector, which consisted of ACC terms of different lengths. RF is used to calculate the Gini impurity of each ACC term [16]. Since the gap between the Gini impurity of each ACC term is small, we sorted them in descending order, and selected the top 20 to record. We recursed the above process 100 times, calculated the frequency of each ACC term, and obtained the final feature importance distribution.

#### Machine learning models

Machine learning (ML) techniques are used to determine the parameters of a data-driven model, which would translate a given input to the correct output [14]. Protein immunogenicity prediction is a classification problem in our study, as the input needs to be mapped to a discrete output and our purpose is an alignment-independent classifier model. As it is not known yet which models are more suitable for predicting the immunogenicity of the proteins, here, eight ML models were applied in the present study: Partial Least Squares (PLS) Regression [17, 18], K Nearest Neighbours (kNN) [19], Adaptive Boosting (Ada) [20], Logistic Regression (LR) [21], Support Vector Machine (SVM) [22], Random Forest (RF) [23-25], Extreme Gradient Boosting (XGBoost) [25, 26] and A Deep Learning Model (Multi-Layer Perceptron, MLP) [27]. All of the eight models have some applications in bioinformatics, for example, SVM and RF were applied to PubChem bioassay data to create classification models of inhibitors/non-inhibitors of flap endonuclease 1 (FEN1) [28]. Parameters of PLSR model were principal components = 2, maximum iterations = 100, iteration stop condition = 1e-06. Parameters of kNN model were neighbor number = 2, weight mode = uniform, leaves number = 30. Parameters of Adaboost model were maximum iterations = 100, learning rate = 1.0. Parameters of LR model were maximum iterations = 1000, penalty = L2. Parameters of SVM were error costs = 1, kernel function = RBF. Parameters of RF were maximum iterations = 100, feature selection method = gini, maximum depth = 6. Parameters of xgboost were base learning = tree, maximum depth = 6. Parameters of MLP were two hidden layers = 32 & 16, input layer size = input data size, output layer size = 2, activation function = relu, optimization function = adam, maximum iterations = 1000.

#### Soft voting ensemble approach

To ensure highly reliable predictions, we designed a soft voting ensemble classifier [29] based on the best three models. The soft voting ensemble method is the combination of multiple models in which decisions are made on the basis of individual decisions. Predictions could be weighted on the basis of classifier's importance and merged them to get the sum of weighted probabilities in this approach. In contrast of hard voting, soft voting gives better result and performance because it can give more importance and involvement to some specific learning models. The soft voting ensemble method covers up the weakness of individual base classifiers and outperforms the overall results by aggregating the multiple prediction models.

#### Evaluation criteria

The ML models used in the present study were trained by training set, validated by test set and then another layer of validation by the external test set. The predictive ability of the models was measured by Receiver Operating Characteristic (ROC) statistics [30], which includes accuracy, precision, sensitivity (Recall) and F1[31]. The ML models were also assessed by AUC, the area under the ROC curve. It is a quantitative measure of the predictive ability and varies from 0.5 for a random prediction to 1.0 for a perfect prediction. These measures were calculated by the scikit-learn of the python package [32].

#### Antigenic linear B-cell epitope prediction for pp220 protein of ASFV

A total of 17 widely used methods were used to predict linear B-cell epitopes for ASFV pp220 protein, including BCPreds [33], Immune Epitope Database (IEDB) server (Bepipred2.0, Bepipred, Emini Surface Accessibility, Kolaskar & Tongaonkar Antigenicity, Chou & Fasma Beta-Turn, Karplus & Schulz Flexibity, and Parker Hydrophilicity) [34], AAP [35], FBCPred [36], Bcepred (Hydrophilicity, Flexibility, Accessibility, Turns, Exposed Surface, Polarity, and Antigenic Propensity) [37], and ABCpred [38], resulting in a total of 1376 epitopes. 1064 of 1376 epitopes had at least eight amino acids. VaxiJen [39] was employed to evaluate the antigenicity of 1064 B cell epitopes with stringent criteria to filter out epitopes having antigenicity score ≤ 1.3. 29 epitopes were remained. Allergen FP 1.0 [40] and ToxinPred [41] were used to predict the allergenicity and toxicity of the 29 epitopes with 12 epitopes classified as non-allergen and non-toxin. VirusImmu was then applied to predict the antigenicity of the 12 epitopes, finding eight epitopes antigenic and four non-antigenic.

#### Animal experiments

All animal experiments were approved by and carried out in accordance with the guidelines of the Institutional Experimental Animal Welfare and Ethics Committee (the approval number is LL20220603). Five six-month old healthy pigs that had no detectable antibodies against ASFV were provided and housed under standard laboratory conditions in Military Veterinary Institution, Changchun. Blood samples of the five pigs were then collected. Pooled sera of five pigs infected with ASFV SY18ΔMGF/ΔCD2v ($10^4$ $TCID_{50}$/mL, $TCID_{50}$ represents Tissue Culture Infectious Dose) were provided by Jilin Heyuan Bioengineering Co., Ltd. The ASFV infected pigs had viral nucleic acid detected in blood and saliva during the observation period. The pooled sera from the healthy pigs and ASFV infected pigs were used for indirect ELISA assays, respectively.

#### Peptide Synthesis

The eight linear B-cell epitopes and two control peptides were synthesized by Scilight-Peptide Inc., Beijing, China via a practical approach of Fmoc solid-phase peptide synthesis. The unsophisticated peptides were purified using a Varian ProStar 218 high-performance liquid chromatography (HPLC) instrument with an Agilent Venusil MP C18 reversed phase column. Peptides were eluted with a linear gradient of water, H2O, and acetonitrile, CAN, (both having 0.05% TFA) at a flow rate of 1 mL/min. The separation was monitored at 220 nm using UV detection. Then peptides were subjected to Voyager-DE STR mass spectrometric (MS) analysis. The solvents for gradient elution HPLC are: solvent A, CAN 2%, TFA 0.05% and solvent B, CAN 90%, TFA 0.05%. Peptides were dissolved in deionized H2O at a final concentration of 20mg/ml and stored at -20$^{o}$C until further use.

#### ASFV-specific IgG assay in pig (Indirect ELISA)

96-well polystyrene microplates (Oriental Ocean Global Health, China) were coated with peptides 2 μg/mL (50μL/well) in carbonate bicarbonate buffer pH9.6 and the plates were incubated at 4$^{o}$C overnight. The plates were then blocked at 37 °C for one hour with PBS (Solarbio, China, cat no:A8020) pH 7.4 in 5% skim milk (blocking buffer) and washed with PBST(0.1% Tween-80) three times. Serial dilutions of sera in dilution buffer (Solarbio, China, cat no: P1010) were added to the plates and incubated at 37$^{o}$C for 30 minutes. HRP-conjugated goat anti-pig IgG (abcam, cat no:ab6915, 1:5000 dilution) was added to the plates, and the plates were incubated in incubator (ZHCHENG, China, ZXDP-B2050) at 37$^{o}$C for 30 minutes and washed with PBST three times. The assay was developed for 10min at 37$^{o}$C with 50 μL of TMB substrate solution (Solarbio,China, cat no: PR1200), stopped by the addition of 50 μL of stop solution (Solarbio, China, cat no:C1058), and then the optical density (OD) was measured at 450 nm (TECAN, INFINITE F50). The endpoint titre was defined as the highest reciprocal serum dilution that yielded an absorbance ≥2.1-fold over negative control serum values (OD value is set as 0.05, if it is less than 0.05). The indirect ELISA were performed at the biosafety level 2 facilities in Beijing Institute of Microbiology and Epidemiology, China.

### References


1. Consortium UP: **UniProt: a worldwide hub of protein knowledge**. *Nucleic acids research* 2018.
2. Coordinators NCIR: **Database resources of the National Center for Biotechnology Information**. *Nucleic Acids Research* 2000(D1):D1.
3. Flower DR, Doytchinova IA: **VaxiJen: a server for prediction of protective antigens, tumour antigens and subunit vaccines**. *BMC Bioinformatics* 2007, **8**(1):4.
4. Altschul SF, Madden TL, Schäffer AA, Zhang J, Zhang Z, Miller W, Lipman DJ: **Gapped BLAST and PSI-BLAST: a new generation of protein database search programs**. *Nucleic Acids Res* 1997, **25**(17):3389-3402.
5. Yang B, Samantha S, Xiang Z, He Y: **Protegen: a web-based protective antigen database and analysis system**. *Nucleic Acids Research* 2011(suppl_1):D1073-D1078.
6. Hellberg S, Sjoestroem M, Skagerberg B, Wold S: **Peptide quantitative structure-activity relationships, a multivariate approach**. *Journal of Medicinal Chemistry* 1987, **30**(7):1126-1135.
7. **Polypeptide sequence property relationships in Escherichia coil based on auto cross covariances**. *Chemometrics & Intelligent Laboratory Systems* 1995.
8. M., Lapinsh: **Classification of G-protein coupled receptors by alignment-independent extraction of principal chemical properties of primary amino acid sequences**. *Protein Science* 2002, **11**(4):795-805.
9. **New quantitative descriptors of amino acids based on multidimensional scaling of a large number of physical–chemical properties**. *Molecular modeling annual* 2001, **7**(12):445-453.
10. Dimitrov I, Zaharieva N, Doytchinova I: **Bacterial Immunogenicity Prediction by Machine Learning Methods**. *Vaccines (Basel)* 2020, **8**(4).
11. Wold S, Jonsson J, Sjrstrm M, Sandberg M, Rnnar S: **DNA and peptide sequences and chemical processes multivariately modelled by principal component analysis and partial least-squares projections to latent structures**. *Analytica Chimica Acta* 1993, **277**(2):239-253.
12. Shahlaei, Mohsen: **Descriptor Selection Methods in Quantitative Structure–Activity Relationship Studies: A Review Study**. *Chemical Reviews* 2013, **113**(10):8093-8103.
13. Arkadiusz Z. Dudek TA, Jorge Galvez: **Computational Methods in Developing Quantitative Structure-Activity Relationships (QSAR): A Review**. *Combinatorial Chemistry & High Throughput Screening* 2006, **9**(3):-.
14. Bonetta R, Valentino G: **Machine learning techniques for protein function prediction**. *Protns Structure Function and Bioinformatics* 2020.
15. Kuhn M, Johnson K: **Applied Predictive Modeling**: Springer New York; 2013.
16. **Sequence Based Prediction of DNA-Binding Proteins Based on Hybrid Feature Selection Using Random Forest and Gaussian Naive Bayes**. *PLOS ONE* 2014, **2014,9(1)**(-):-.
17. Mao HH, Chao S: **Advances in Vaccines**. *Adv Biochem Eng Biotechnol* 2020, **171**:155-188.
18. Abdi H: **Partial Least Squares (PLS) Regression**. 2003.
19. Hu L, Tao H, Shi X, Lu WC, Cai YD, Kuo-Chen C, Christos O: **Predicting Functions of Proteins in Mouse Based on Weighted Protein-Protein Interaction Network and Protein Hybrid Properties**. *Plos One* 2011, **6**(1):e14556.
20. Wu P, Zhao H: **Some Analysis and Research of the AdaBoost Algorithm**. In: *Intelligent Computing and Information Science: 2011// 2011; Berlin, Heidelberg*. Springer Berlin Heidelberg: 1-5.
21. Stoltzfus JC: **Logistic regression: a brief primer**. *Acad Emerg Med* 2011, **18**(10):1099-1104.
22. Boser BE: **A Training Algorithm for Optimal Margin Classifiers**. *Proceedings of Annual Acm Workshop on Computational Learning Theory* 2008, **5**:144--152.
23. Zhang, Zulkernine: **A hybrid network intrusion detection technique using random forests**. - 2006.

24. Breiman: **Random forests**. *MACH LEARN* 2001, **2001,45(1)**(-):5-32.
25. Fuyi, Li, Chen, Mingjun, Wang, Geoffrey, Webb, Yang, Zhang: **GlycoMine: a machine learning-based approach for predicting N-, C- and O-linked glycosylation in the human proteome**. *Bioinformatics* 2015.
26. **XGBoost: A Scalable Tree Boosting System**. In: *the 22nd ACM SIGKDD International Conference: 2016*.
27. **A deep learning-based method for grip strength prediction: Comparison of multilayer perceptron and polynomial regression approaches**. *PLOS ONE* 2021, **16**.
28. Deshmukh AL, Chandra S, Singh DK, Siddiqi MI, Banerjee D: **Identification of human flap endonuclease 1 (FEN1) inhibitors using a machine learning based consensus virtual screening**. *Mol Biosyst* 2017, **13**(8):1630-1639.
29. Sherazi S, Bae JW, Lee JY: **A soft voting ensemble classifier for early prediction and diagnosis of occurrences of major adverse cardiovascular events for STEMI and NSTEMI during 2-year follow-up in patients with acute coronary syndrome**. *PLOS ONE* 2021, **16**.
30. Carter JV, Pan J, Rai SN, Galandiuk S: **ROC-ing along: Evaluation and interpretation of receiver operating characteristic curves**. *Surgery* 2016:1638-1645.
31. Larrañaga P, Calvo B, Santana R, Bielza C, Robles V: **Machine learning in bioinformatics**. *Briefings in Bioinformatics* 2006, **7**(1):86-112.
32. Pedregosa F, Varoquaux G, Gramfort A, Michel V, Thirion B, Grisel O, Blondel M, Müller A, Nothman J, Louppe G: **Scikit-learn: Machine Learning in Python**. 2012.
33. El-Manzalawy Y, Dobbs D, Honavar V: **Predicting linear B-cell epitopes using string kernels**. *J Mol Recognit* 2008, **21**(4):243-255.
34. Peters B, Sidney J, Bourne P, Bui HH, Buus S, Doh G, Fleri W, Kronenberg M, Kubo R, Lund O *et al*: **The immune epitope database and analysis resource: from vision to blueprint**. *PLoS Biol* 2005, **3**(3):e91.
35. Chen J, Liu H, Yang J, Chou KC: **Prediction of linear B-cell epitopes using amino acid pair antigenicity scale**. *Amino Acids* 2007, **33**(3):423-428.
36. El-Manzalawy Y, Dobbs D, Honavar V: **Predicting flexible length linear B-cell epitopes**. *Comput Syst Bioinformatics Conf* 2008, **7**:121-132.
37. Saha S, Raghava GPS: **BcePred: Prediction of continuous B-cell epitopes in antigenic sequences using physico-chemical properties**. In: *ICARIS 2004, LNCS3239: 2004 2004*. Springer: 197-204.
38. Saha S, Raghava GP: **Prediction of continuous B-cell epitopes in an antigen using recurrent neural network**. *Proteins* 2006, **65**(1):40-48.
39. **Immunogenicity Prediction by VaxiJen: A Ten Year Overview**. *Journal of Proteomics & Bioinformatics* 2017, **10**(11).
40. Dimitrov I, Naneva L, Doytchinova I, Bangov I: **AllergenFP: allergenicity prediction by descriptor fingerprints**. *Bioinformatics* 2014, **30**(6):846-851.
41. Gupta S, Kapoor P, Chaudhary K, Gautam A, Kumar R, Open Source Drug Discovery C, Raghava GP: **In silico approach for predicting toxicity of peptides and proteins**. *PLoS One* 2013, **8**(9):e73957.

**S2: A more detailed description of relevant models in present study.**

**Machine learning models**

Machine learning (ML) techniques are used to determine the parameters of a data-driven model, which would translate a given input to the correct output[6]. Protein immunogenicity prediction is a classification problem in our study, as the input needs to be mapped to a discrete output and our purpose is an alignment-independent classifier model. Eight ML models were applied in the present study and are described below.

Partial Least Squares (PLS) Regression (PLSR) is a versatile algorithm that can be used for predictive modelling, which uses a regression method for classification[7]. PLSR generalizes and combines features from principal component analysis and multiple regression[8]. This method classifies samples based on a particular threshold (larger than the threshold:1, smaller than the threshold:0).

K Nearest Neighbours (kNN) algorithm is a non-parametric method which classifies a given observation through a majority vote of the labels of the closest k points in a given feature space[9]. The majority voting procedure suffers when the class distribution is skewed.

Adaptive Boosting (Ada) classifier is a meta-algorithm classifier. The adaBoost classifier uses the same base classifier (weak classifier), assigns different weight parameters based on the error rate of the classifier, and finally accumulates the weighted prediction results as the output[10].

Logistic Regression (LR) models are defined as statistical models, which describe the relationship between a qualitative dependent variable and an independent variable[11]. In LR, a sigmoid function is used as a squashing function to map a real-valued input to a range from 0 to 1.

Support Vector Machine (SVM) seeks to maximize the separation between points corresponding to different classes in some n-dimensional space, and therefore, determines a maximum-margin hyperplane[12], which is achieved by means of kernel functions.

Random Forest (RF) is an ensemble classification approach, which has been widely used for pattern recognition in bioinformatics[13]. It can provide the high prediction performance for classification task[14]. The algorithm of RF is based on the ensemble of a large number of decision trees[15], where each tree gives a classification and the forest chooses the final classification having the most votes (over all the trees in the forest).

Extreme Gradient Boosting (XGBoost) is an advanced implementation of decision-tree-based ensemble ML algorithm, which was first proposed by Tianqi Chen and Carlos Guestrin in 2011 and has been continuously optimized and improved in the follow-up study of many scientists[16, 17].

Multi-Layer Perceptron (MLP) is a typical neural network architecture with one or more hidden layers between the input and output layer. Goal of training is to learn appropriate values for the weights so as to obtain a correct output for a given input[18]. In the present study, we set up two hidden layers with 32 neurons and 16 neurons respectively.

#### References


1. Ghosh N, Sharma N, Saha I. Immunogenicity and antigenicity based T-cell and B-cell epitopes identification from conserved regions of 10664 SARS-CoV-2 genomes, Infect Genet Evol 2021;92:104823.
2. Chen HZ, Tang LL, Yu XL et al. Bioinformatics analysis of epitope-based vaccine design against the novel SARS-CoV-2, Infect Dis Poverty 2020;9:88.
3. Crooke SN, Ovsyannikova IG, Kennedy RB et al. Immunoinformatic identification of B cell and T cell epitopes in the SARS-CoV-2 proteome, Sci Rep 2020;10:14179.
4. Grifoni A, Sidney J, Zhang Y et al. A Sequence Homology and Bioinformatic Approach Can Predict Candidate Targets for Immune Responses to SARS-CoV-2, Cell Host Microbe 2020;27:671-680.e672.
5. Kar T, Narsaria U, Basak S et al. A candidate multi-epitope vaccine against SARS-CoV-2, Sci Rep 2020;10:10895.
6. Bonetta R, Valentino G. Machine learning techniques for protein function prediction, Protns Structure Function and Bioinformatics 2020.
7. Bacterial Immunogenicity Prediction by Machine Learning Methods, Vaccines 2020;8:709.
8. Abdi H. Partial Least Squares (PLS) Regression 2003.
9. Hu L, Tao H, Shi X et al. Predicting Functions of Proteins in Mouse Based on Weighted Protein-Protein Interaction Network and Protein Hybrid Properties, PLOS ONE 2011;6:e14556.
10. Wu P, Zhao H. Some Analysis and Research of the AdaBoost Algorithm. In: Intelligent Computing and Information Science. Berlin, Heidelberg, 2011, p. 1-5. Springer Berlin Heidelberg.
11. Stoltzfus JC. Logistic regression: a brief primer, Acad Emerg Med 2011;18:1099-1104.
12. Boser BE. A Training Algorithm for Optimal Margin Classifiers, Proceedings of Annual Acm Workshop on Computational Learning Theory 2008;5:144--152.
13. Zhang, Zulkernine. A hybrid network intrusion detection technique using random forests, - 2006.
14. Breiman. Random forests, MACH LEARN 2001;2001,45(1):5-32.
15. Fuyi, Li, Chen et al. GlycoMine: a machine learning-based approach for predicting N-, C- and O-linked glycosylation in the human proteome, Bioinformatics 2015.
16. XGBoost: A Scalable Tree Boosting System. In: the 22nd ACM SIGKDD International Conference. 2016.
17. Chen Y, Li Y, Rajiv N et al. Gene expression inference with deep learning, Bioinformatics:1832.
18. A deep learning-based method for grip strength prediction: Comparison of multilayer perceptron and polynomial regression approaches, PLOS ONE 2021;16.
